# The minimal intrinsic stochasticity of constitutively expressed eukaryotic genes is sub-Poissonian

**DOI:** 10.1101/2023.03.06.531283

**Authors:** Douglas E. Weidemann, Abhyudai Singh, Ramon Grima, Silke Hauf

## Abstract

Stochastic variation in gene products (“noise”) is an inescapable by-product of gene expression. Noise must be minimized to allow for the reliable execution of cellular functions. However, noise cannot be suppressed beyond an intrinsic lower limit. For constitutively expressed genes, this limit is believed to be Poissonian, meaning that the variance in mRNA numbers cannot be lower than their mean. Here, we show that several cell division genes in fission yeast have mRNA variances significantly below this limit, which cannot be explained by the classical gene expression model for low-noise genes. Our analysis reveals that multiple steps in both transcription and mRNA degradation are essential to explain this sub-Poissonian variance. The sub-Poissonian regime differs qualitatively from previously characterized noise regimes, a hallmark being that cytoplasmic noise is reduced when the mRNA export rate increases. Our study re-defines the lower limit of eukaryotic gene expression noise and identifies molecular requirements for ultra-low noise which are expected to support essential cell functions.

## Introduction

All gene expression is “noisy”. Gene products fluctuate stochastically, which is an inescapable consequence of the nature of gene expression (*1–4*). Gene expression, like other intracellular processes, relies on molecule-molecule (in the case of gene expression often protein-DNA) interactions. When both molecules in such a reaction are abundant, their interaction is frequent and not limiting for the reaction itself. However, for several stages of gene expression, one binding partner may be present in very low numbers. For example, to initiate transcription, proteins must bind a gene promoter, which typically only has one to four copies per cell. As a result, stochasticity becomes apparent and the intracellular number of mRNA molecules that has been transcribed from a specific gene varies significantly over time and between cells (*5–8*).

Gene expression noise differs substantially between genes. For genes with high variation, some cells in the population may contain dozens or hundreds of mRNA molecules expressed from that gene, and others none (*7, 9–13*). In a few cases, this large variation has been shown to be advantageous, since it allows “bet-hedging”, i.e. some cells in the population are well prepared for a change in environmental conditions, whereas others don’t produce gene products that are not currently needed (*14–17*). Highly variable expression of a single gene is also exploited in development, where it may specify cell fate and generate a mix of cells with different phenotypes (*18–20*). At the other end of the spectrum, “housekeeping” genes, i.e. genes required for core cellular processes and often required for survival, are expressed stably with low noise (*21*–*23*). Cell cycle genes have also been found to show low intrinsic noise (*24*).

High-noise regimes have been extensively studied. One contributing factor to noisy expression is toggling between an inactive and active transcription state, so that mRNAs are produced in “bursts”, leading to large variation in mRNA abundance (*25*). The large noise created by bursty expression can be modulated downstream of mRNA synthesis (*26, 27*). For example, slow mRNA export from the nucleus may buffer bursty mRNA synthesis and thereby lower noise in the cytoplasm relative to noise in the nucleus (*11, 28–30*). By contrast, molecular processes downstream of mRNA synthesis, such as multi-step mRNA decay, may also enhance noise created during transcription (*31*).

Less is known about the low-noise regime. Low noise genes are thought to be expressed from constitutive promoters that stay ‘on’, and do not toggle into an ‘off’ state (*32, 33*). Widely used models for this constitutive expression assume that mRNA is produced with a single rate-limiting step from an active promoter and that the wait time between synthesis events follows an exponential distribution (*33–36*). This has been experimentally confirmed for two constitutively expressed genes in budding yeast (*37*). Under these assumptions, the steady-state mRNA numbers in the cell population will follow a Poisson distribution, independent of the precise nature of the mRNA degradation process (*38*). A Poisson distribution is characterized by the variance being equal to the mean, or the Fano factor (the variance divided by the mean) being equal to 1. Consistent with this proposed expression model, several constitutive genes in yeast show distributions of mRNA numbers that closely follow a Poisson distribution (*6, 12, 39, 40*). This distribution is typically considered the “noise floor”, i.e. the lower limit of intrinsic stochasticity in gene expression (*11, 40–43*).

To shed additional light on low-noise gene expression, we have investigated a group of constitutively expressed *S. pombe* genes, that are important for cell division and show low mRNA and protein noise (*44, 45*). The protein products of these genes contribute to the spindle assembly checkpoint (SAC, also known as mitotic checkpoint). This signalling pathway operates during cell division to detect chromosomes that are not correctly attached to the mitotic spindle and halts the execution of anaphase as a response (*46–49*). SAC signalling involves several protein-protein interactions, and the relative ratio of SAC proteins is important for SAC function (*44, 50, 51*). Hence, low expression noise of these genes supports SAC function, making it important to understand how low noise is achieved.

Strikingly, rather than Poissonian mRNA distributions, we found sub-Poissonian mRNA distributions for these genes, i.e. Fano factors less than 1. In the cytoplasm, the Fano factor could be as low as 0.5. We also examined other, non-SAC, low noise genes, as well as re-examined published results, and found a spectrum of sub-Poissonian to Poissonian mRNA distributions for constitutively expressed genes. The cytoplasmic mRNA distributions of these genes can have Fano factors that are lower, the same, or higher than those of the nuclear mRNA distributions. Altogether, this suggests that constitutively expressed genes are not a homogenous group, but instead can differ substantially in mRNA noise. Importantly, we conclusively establish that the lower limit of intrinsic stochasticity for constitutively expressed genes is sub-Poissonian, not Poissonian.

## Results

### Gene expression is regulated independently for each *S. pombe* SAC gene

SAC genes need to cooperate in one signalling pathway, and their relative stoichiometries are important for function (*44, 50, 51*). Stoichiometries could be more easily maintained if expression of these genes was coupled. i.e. if higher or lower expression of one coincided with higher or lower expression of another. Yet, isolated observations suggest that their expression is independent (*44, 52*). We systematically tested for interdependence at both the mRNA and protein levels (Fig. 1). Except for the encoded protein itself, the levels of the SAC proteins Mad1, Mad2, Mad3, and Bub1 were largely unaffected by single deletion of *mad1*, *mad2*, *mad3*, or *bub1* (Fig. 1A), and so were the mRNA levels measured by quantitative PCR (Fig. 1B). Mad1 and Mad2 form a tight 2:2 complex that is central for SAC function (*53, 54*). Yet, *mad1* and *mad2* mRNA concentrations did not correlate in single cells (Fig. 1C); nor did those of *mad1* and *mad3* (Fig. 1D). This agrees with findings in budding yeast that the concentrations of constitutively expressed mRNAs coding for subunits of stable protein complexes do not necessarily correlate (*39*). We did observe a slight increase in mRNA concentration of *mad1, mad2*, and *mad3* by quantitative PCR when expression of *bub1* was increased by 2- to 3-fold, and a similar trend for *mad3* in a *mad1* overexpression (Fig. 1B). However, the changes were so subtle that we conclude that SAC genes are expressed mostly or wholly independently.

**Figure 1.**
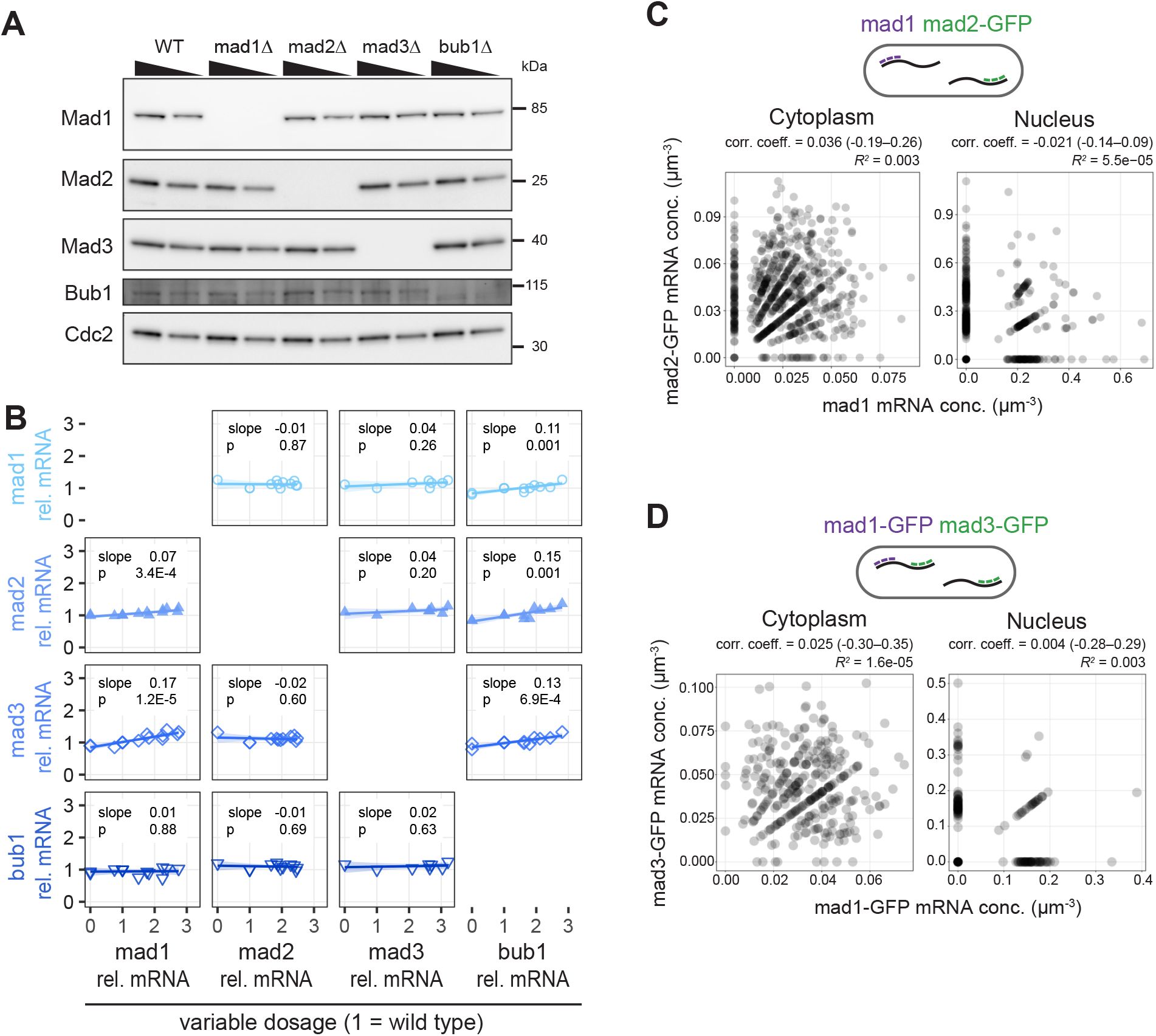
Expression of individual SAC genes is not correlated with other SAC genes. **(A)** Immunoblot of cell extracts from the indicated strains; 70 % of the extract is loaded in each second lane. Antibodies against the endogenous proteins. Cdc2 serves as loading control. **(B)** Strains in which SAC genes were either deleted (dosage = 0) or slightly overexpressed (dosage > 1) by expression from a second genetic locus were analyzed for the concentration of SAC gene mRNAs by quantitative PCR; the mRNA concentration is given relative to that in a wild-type strain. Parameters of linear regression are shown. **(C)** Both mad1 and mad2-GFP mRNA concentrations were determined in the same cells by single-molecule mRNA FISH (n = 734 cells). Cells expressed untagged mad1 and mad2-GFP from their respective endogenous locus; mad1 mRNA was targeted with mad1-specific probes, mad2 mRNA with probes against GFP. Nuclei were stained by DAPI. The correlation coefficient was calculated as Kendall’s tau-b; the 95 % confidence interval is given in brackets. **(D)** Both mad1-GFP and mad3-GFP mRNA concentrations were determined in the same cells by single-molecule mRNA FISH (n = 329 cells). Cells expressed mad1-GFP and mad3-GFP from their respective endogenous locus; mad1-specific probes and probes against GFP were used. GFP dots co-localizing with mad1 were considered mad1-GFP, all others were considered mad3-GFP. Nuclei were stained by DAPI. The correlation coefficient was calculated as Kendall rank correlation coefficient, tau-b; the 95 % confidence interval is given in brackets.

### SAC genes show sub-Poissonian distributions of mRNA numbers, more narrow than those of other low noise genes

Without cross-talk between SAC expression levels, it is important that expression of these genes is stable, so that levels do not deviate beyond the range that allows for SAC function. We therefore assessed variability in SAC gene mRNA numbers at the single cell level by single-molecule mRNA fluorescence in-situ hybridization, smFISH (*55*). The FISH spot intensities in the cytoplasm were homogenously distributed in a narrow range, consistent with representing single mRNA molecules (Fig. S1). Spots with higher intensities were almost exclusively observed for highly expressed genes and only in the nucleus, presumably representing the transcription site (Fig. S1, S9). We counted spots as a single mRNA if they had an intensity below the 95^th^ percentile of spot intensities in the cytoplasm (Fig. S1C). This avoids that technical noise from uneven labelling is interpreted as biological noise. For brighter spots, the number of mRNA contained in each spot was estimated as value of its intensity divided by the median intensity of a cytoplasmic spot. This ensures that several mRNAs at the same position are accurately counted as such.

The cell-to-cell variation in mRNA numbers is a result of intrinsic and extrinsic noise sources (*1, 56, 57*). A considerable extrinsic influence is cell size. Mean mRNA numbers increase with cell size (see Fig. 2B, S2 for examples), so that mRNA concentrations (number divided by cell volume) remain approximately constant as cells grow (*40, 43, 58–60*). To exclude the influence of size, we either considered only cells within a given size range (Fig. 2B, S2A) or mathematically corrected the variance for the size effect (*58*) (Fig. 2C, S2B; see Methods). When excluding the influence of cell size, all four SAC genes (*mad1 mad2*, *mad3*, and *bub1*) showed Fano factors of the mRNA distributions that were clearly below 1 and for *mad1* reaching as low as 0.5 (Fig. 2B,C; S2A,B; Supplemental Information, A.2). This was independent of whether we used cell length or cell volume as the size parameter (Fig. S2C), and for *mad1* was the case regardless of whether a GFP tag was present or not (Fig. 2B,C; S2A,B; Supplemental Information, A.2). Further statistical analysis showed that the Fano factor of the SAC genes being below 1 could not be explained by finite sample size effects (Supplemental Information, B.4). A Poisson distribution (Fano factor = 1, Fig. 2A) would have been expected if SAC genes were expressed with a single rate-limiting step, as is typically assumed for constitutively expressed genes. Hence, this observation suggests that other, potentially more complex mechanisms account for the ultra-low Fano factor of the SAC gene mRNA distributions.

**Figure 2.**
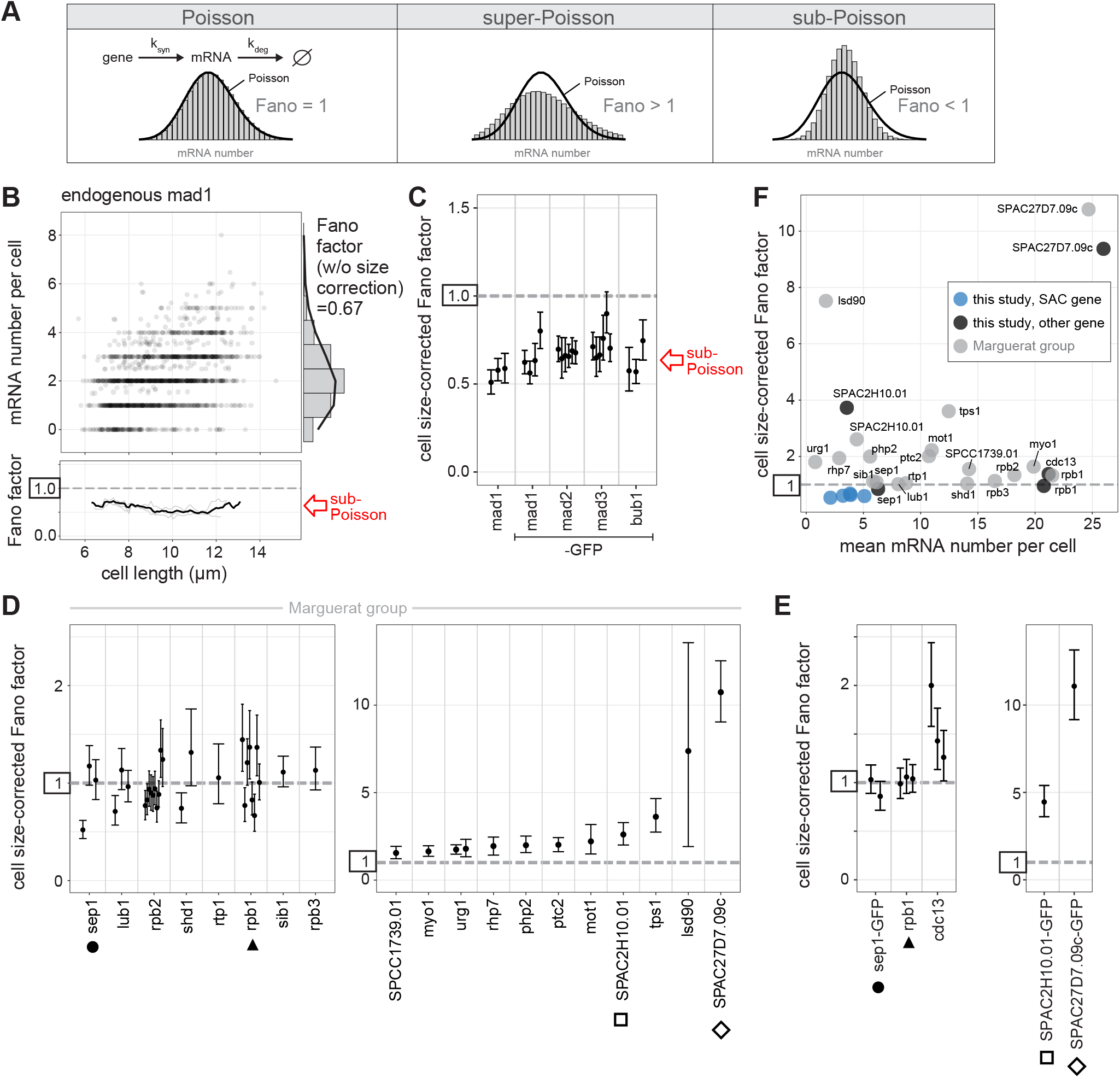
SAC genes show sub-Poissonian distributions of mRNA numbers. **(A)** Constitutive gene expression with a single rate-limiting step in transcription and mRNA degradation yields a Poisson distribution of mRNA numbers per cell (variance = mean, Fano factor = 1). Super-Poissonian distributions have a larger variance than the mean (Fano factor > 1), sub-Poissonian distributions have a lower variance than the mean (Fano factor < 1). Here, a negative binomial and a binomial distribution are used as illustration. Black lines are Poisson distributions with the same mean as the data in the histogram. **(B)** Top: Scatter plot of cell length versus mRNA number per cell for untagged mad1 (n = 1,619 cells; data from 3 replicates combined). Right: Histogram of mRNA number across all cells with fit to Poisson distribution (black line). Bottom: The Fano factor was determined in a sliding window spanning 1 µm of cell length. The Fano factors for single replicates are shown in light gray; the Fano factor for the pooled data in black. See Fig. S2 for other SAC genes. **(C)** Cell size-corrected Fano factors and their 95 % confidence interval were calculated from smFISH data as in Padovan-Merhar et al., Mol Cell 2015. At least three replicate experiments for each gene; the number of cells in single replicates ranged from 209 to 778. **(D)** Cell size-corrected Fano factors and their 95 % confidence interval were calculated as in (C) using published smFISH data by the Marguerat group (Saint et al., Nature Microbiology 2019; Sun et al., Curr Biol 2020). Genes that were also analyzed in this study (see E) are marked. Between 1 and 10 replicate experiments per gene; the number of cells in single replicates ranged from 103 to 441. **(E)** Cell size-corrected Fano factors and their 95 % confidence interval for non-SAC genes were calculated as in (C). Genes also analyzed by the Marguerat group (see D) are marked. Between 1 and 3 replicate experiments per gene; the number of cells in single replicates ranged from 236 to 676. **(F)** Cell size-corrected Fano factors were plotted against the mean mRNA number per cell for individual genes. Data from replicates for each gene were pooled to determine the cell size-corrected Fano factor. Genes analyzed in this study are shown in black; SAC genes from this study in blue; genes analyzed by the Marguerat group are shown in gray. Between 236 and 3,501 cells per gene for our data, and between 103 and 2,564 cells per gene for the Marguerat group’s data. For data from this study the “spot count” for mRNA number was used for better comparability to the Marguerat data; panels (B), (C), and (E) use the “hybrid count” for mRNA number (see Fig. S1).

The distribution of mRNA numbers in *S. pombe* for several other genes had previously been examined, and some genes showed mRNA distributions close to a Poisson distribution (*12, 40*). When we corrected these published data for cell size in the same way we had done for the SAC genes, five genes (*sep1*, *lub1*, *rpb2*, *shd1*, *rpb1*) showed single experiments with evidence of sub-Poissonian mRNA distributions, but none of them as consistently across multiple experimental replicates as we observed for the SAC genes (Fig. 2D). After averaging Fano factors from all experimental replicates, none of these genes had a Fano factor significantly below 1 (Supplemental Information Fig. A-2). We re-tested two of these genes (*sep1* and *rpb1*), as well as two higher noise genes (*SPAC2H10.01*, *SPAC27D7.09c*), with our workflow and found the mRNA numbers and variance to be highly consistent with those obtained previously, cross-validating the data (Fig. 2E, S2D). We did not observe a correlation between mean mRNA numbers and cell size-corrected Fano factors across these genes (Fig. 2F). SAC genes show low mean mRNA numbers (means of 2–5). Other genes with similarly low mean mRNA number can show considerably larger Fano factors. In summary, the SAC genes show the lowest Fano factor in mRNA number distributions among all *S. pombe* genes for which cell size and mRNA numbers have been reported.

### Characterization of SAC gene structure shows short promoters, variable 5’ UTR lengths, and typical 3’ UTR lengths

The sub-Poissonian mRNA number distribution of SAC genes raised the question whether these genes share any peculiar characteristics, possibly in the promoter or the mRNA. We first mapped the 5’ and 3’UTRs, because they may influence mRNA dynamics, and because the 5’UTR length provides insight into the position of the promoter (Fig. 3). The 5’UTRs of *mad1* and *mad2* are extremely short (∼10 nucleotides), below the 6^th^ percentile of all *S. pombe* genes (*61*). The 5’UTR of *mad3* (∼270 nucleotides) was longer than the interquartile range observed in *S. pombe*, whereas that of *bub1* (∼60 nucleotides) was within the interquartile range. Our results were highly consistent with a recent genome-wide study identifying transcription start sites in *S. pombe* (Fig. S3A) (*61*). The very short 5’UTRs were surprising, since short 5’UTRs have been shown to impair translation from the first start codon in the mRNA (*62*). The only gene ontologies that were enriched for genes with such short 5’UTRs were those of ribosomal proteins (Fig. S3B). Although the short 5’UTRs of *mad1* and mad2 are unusual, the strong differences in 5’UTR length between the SAC genes imply that there is no direct correlation with the sub-Poissonian mRNA distributions.

**Figure 3.**
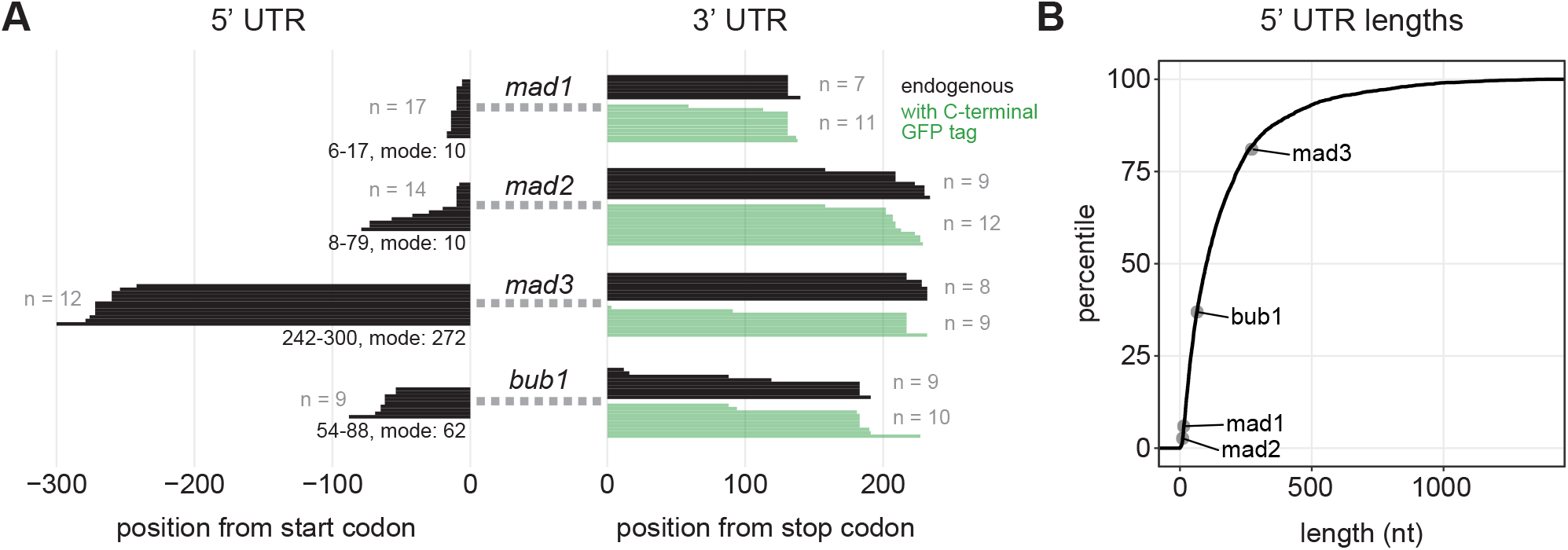
5’ UTR lengths of SAC genes range from untypically short to untypically long. **(A)** Lengths of 5’ and 3’ untranslated regions (UTRs) determined by RACE-PCR; black, endogenous gene; green, ymEGFP-tagged gene. For the 5‘ UTRs, the spread in lengths and the most frequent length is given at the bottom. **(B)** Lengths of SAC gene 5’ UTRs in comparison to genome-wide data from Thodberg et al., 2019, displayed as empirical cumulative distribution (4,710 transcripts).

Promoters have a large influence on gene expression noise (*15, 42, 63–68*). The location, size, and characteristics of most promoters in *S. pombe* remain poorly defined (*61, 69*). To map the promoter regions of the SAC genes, we transferred these genes with a variable length of sequence upstream of the transcription start site (3 to 770 nucleotides) to two different intergenic loci: (i) the 3.8 kb region next to *wis1*, one of the largest intergenic regions in *S. pombe*, and (ii) the 0.4 kb region next to *leu1*, which is a frequently used integration site in *S. pombe* (*70*) (Fig. 4A). We used two loci to be able to control for effects from the surrounding region. We found that expression only started to break down when less than 50 to 100 base pairs of upstream sequence were kept (Fig. 4B,C; S4A, S5). Monitoring mRNA levels by qPCR and protein levels by either immunoblotting or imaging gave similar results (Fig. 4B,C). This suggests that the expression breakdown indeed reflects the inability of the coding sequence to be transcribed. The results from the two intergenic loci were overall consistent, with some quantitative differences observed for *mad2* (Fig. 4B,C; S4A).

**Figure 4.**
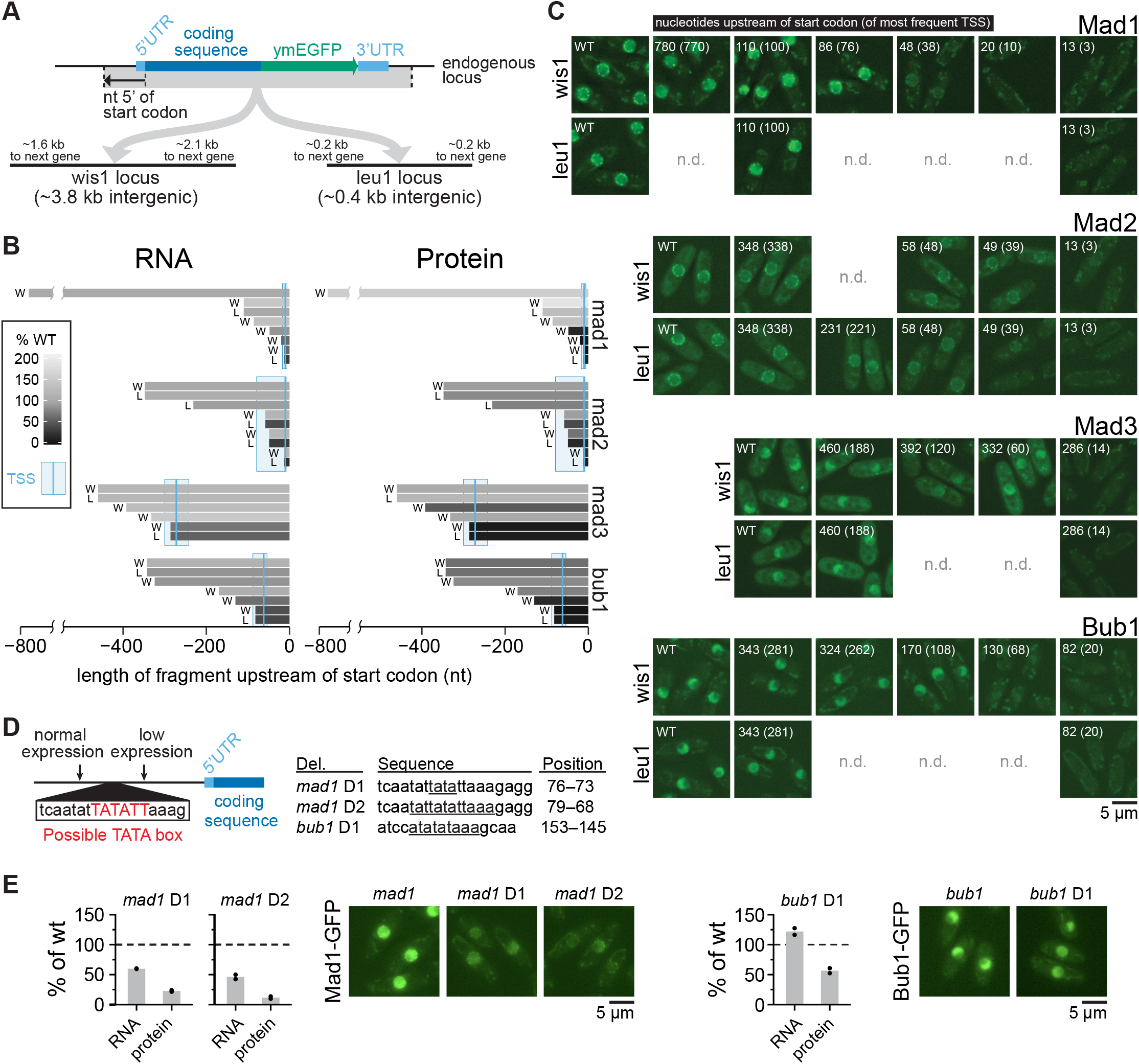
SAC genes have short promoter regions. **(A)** Diagram showing design of integrating SAC genes at two exogenous, intergenic loci. **(B)** Amount of RNA (by qPCR) and protein (by immunoblot quantification, see Fig. S4) from exogenous locus expression (W = wis1 locus, L = leu1 locus) relative to expression from the endogenous locus. The length of bars shows the length of the upstream region (promoter and 5’UTR), shades of gray show the mRNA or protein amount. qPCR primers and immunoblot antibody targeted the GFP tag. **(C)** Representative live-cell images showing GFP fluorescence from GFP-tagged genes at the endogenous locus (WT) or gene fragments inserted at an exogenous locus (leu1 or wis1). Numbers indicate distance in nucleotides that the exogenous locus gene fragment extends upstream of the start codon, or upstream of the most prominent TSS (in brackets). Gamma correction for images to make both high and low expression cells visible: Mad1 = 2.0, Mad2 = 1.8, Mad3 = 1.0, and Bub1 = 1.5. **(D)** Schematic (left) and table (right) of deletions to test the functionality of possible TATA box sequences. Deleted bases are underlined in the table and their positions are given in bp upstream of the start codon. **(E)** Relative expression of RNA (by qPCR) and protein (by immunoblot quantification, see Fig. S4) from deletion mutants in (D) normalized to strains with wild-type promoter sequences (two biological replicates each). One-sample t-tests: mad1 D1 RNA p = 0.007, mad1 D1 protein p = 0.01, mad1 D2 RNA p = 0.04, mad1 D2 protein p = 0.01, bub1 D1 RNA p = 0.16, bub1 D1 protein p = 0.06. Representative live-cell images showing GFP fluorescence of the wild-type gene or the TATA box mutants, all expressed from the exogenous leu1 locus (gamma correction: 1.5).

Based on the short length of the candidate promoter regions, expression is likely driven by a core promoter only. Consistently, *mad1* and *bub1* had TATA box-like sequences in the ∼40 base pair upstream sequence that, when removed, drastically lowered expression (Fig. S5). This was surprising, though, because TATA boxes are typically found in promoters causing large noise (*15, 63, 64, 71*). Mutating the candidate TATA box in *mad1* clearly lowered expression (Fig. 4D,E), suggesting it functions as a genuine TATA box. This was not the case for *bub1* (Fig. 4D,E), but *bub1* contains another TATA box-like sequence upstream (Fig. S5), which may have compensated. We conclude that the expression of SAC genes is likely driven by core promoter sequences only.

### Regulatory SAC gene sequences are sufficient for narrow mRNA distributions in the Poisson-range, but not for sub-Poissonian mRNA distributions

To test whether the identified promoters are sufficient for expression with low Fano factors, we placed the coding sequences of two high-noise genes under the *mad2* and *mad3* promoters at the exogenous locus (Fig. 5A). We used *rad21*, which is a cell cycle-regulated gene whose Fano factor can increase to around 12 during expression late in the cell cycle (Fig. 5B; S6A,B) (*72–74*), and *nmt1*, which is a thiamine-responsive, highly variable gene (*12, 45*). When expressing *rad21* and *nmt1* from the *mad2* and *mad3* promoter, the cell cycle-regulation of *rad21* disappeared (Fig. 5C; S6), and the Fano factor of the mRNA distributions of *rad21* and *nmt1* was lowered to a value slightly above 1 (Fig. 5D). This was close to, but not quite as low, as the *mad2* and *mad3* coding sequences expressed from their own promoter at the exogenous locus (Fig. 5D). Overall, this confirms that promoter sequences have a large influence (*67*), but suggests that a combination of gene elements is required for the very narrow mRNA distributions observed for SAC genes.

**Figure 5.**
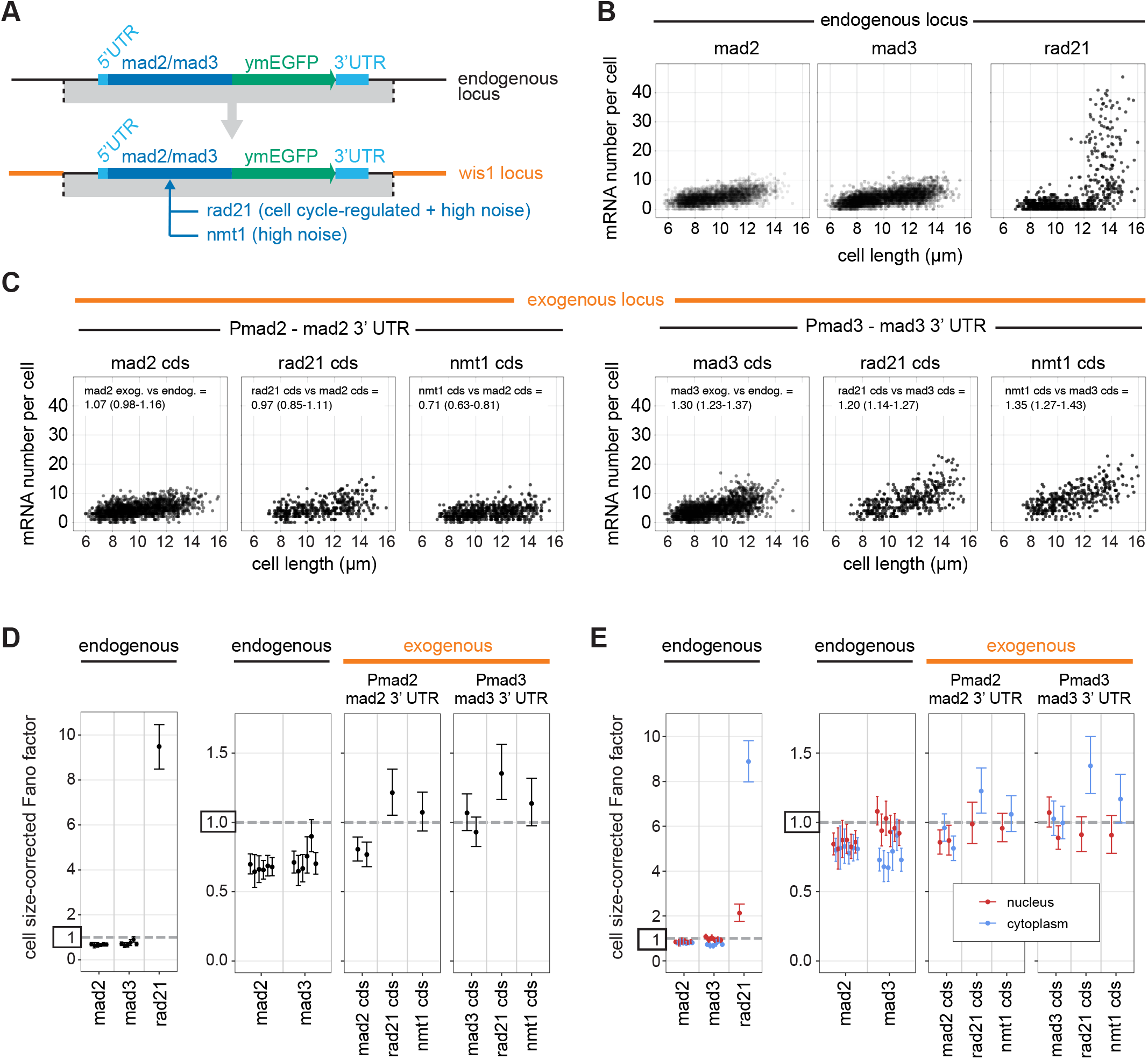
Small surrounding regions are sufficient for low-noise expression, but not for sub-Poissonian mRNA distributions. **(A)** Design of the experiment: *mad2*-GFP and the region surrounding it or *mad3*-GFP and the region surrounding it were integrated at the large intergenic locus adjacent to the wis1 gene. At this location, the coding-sequences of *mad2* or *mad3* were then replaced with those of *rad21*, a gene showing cell cycle-regulated and high-noise expression, or *nmt1*, a high-noise gene. The region around *mad2*-GFP ranged from 48 nucleotides 5’ of the most prominent TSS to 521 nucleotides 3’ of the stop codon; the region around *mad3*-GFP ranged from 270 nucleotides 5‘ of the most prominent TSS to 279 nucleotides 3’ of the stop codon. **(B)** mRNA number relative to cell length at the endogenous locus with the endogenous promoter (*mad2*: 3,501 cells from 6 independent experiments; *mad3*: 2,993 cells from 6 independent experiments; *rad21*: 678 cells from 1 experiment; data for *mad2* and *mad3* are the same as in Fig. 2). **(C)** mRNA number relative to cell length at the exogenous locus for expression from the *mad2* promoter (*Pmad2*) and *mad2* downstream region or *mad3* promoter (*Pmad3*) and *mad3* downstream region (*Pmad2-mad2* cds: 1,308 cells from 2 independent experiments; *Pmad2-rad21* cds: 375 cells from 1 experiment; *Pmad2-nmt1* cds: 557 cells from 1 experiment; *Pmad3-mad3* cds: 1,586 cells from 2 independent experiments; *Pmad3-rad21* cds: 368 cells from 1 experiment; *Pmad3-nmt1* cds: 342 cells from 1 experiment). Generalized linear mixed model estimates for the change in mRNA number per cell between conditions with 95 % confidence interval in brackets are given on top; also see Fig. S6C. **(D)** Cell size-corrected Fano factors and their 95 % confidence interval were calculated from smFISH data as in Fig. 2 using the data shown in (B) and (C). Between 1and 6 replicate experiments for each genotype; the number of cells in single replicates ranged from 209 to 794. **(E)** As in (D), but nuclei were segmented using DAPI staining, and the cell size-corrected Fano factor was calculated separately for mRNA localizing to the nucleus and localizing to the cytoplasm.

To explore the subtle differences in noise between endogenous and exogenous loci, and between the endogenous and exogenous coding sequences, we determined the size-corrected Fano factor separately for the nucleus, where mRNAs are produced, and for the cytoplasm, where mRNAs are ultimately translated. We assigned mRNAs to either the nucleus or the cytoplasm after segmenting nuclei in 3D based on DNA staining and correcting for chromatic aberration (Fig. S7A). This revealed interesting differences. For *mad3* at its endogenous locus, the Fano factor of the mRNA distribution was lower in the cytoplasm than in the nucleus, whereas it was similar for *mad2* (Fig. 5E). When *mad3* was expressed from the exogenous locus, though, its Fano factor now also became similar between nucleus and cytoplasm. Furthermore, *rad21* and *nmt1* expressed from the same regulatory sequences tended to show a higher Fano factor in the cytoplasm than in the nucleus (Fig. 5E). Together, this suggested that an interplay between promoter, coding sequence, and surrounding sequences may be necessary for sub-Poissonian mRNA distributions, and that the sub-Poissonian mRNA expression of SAC genes is maintained or further suppressed in the cytoplasm.

### Most SAC genes show lower mRNA Fano factors in the cytoplasm than in the nucleus

To test for the generality of this observation, we determined size-corrected nuclear and cytoplasmic Fano factors for all SAC genes and the other genes examined previously, all expressed from their endogenous locus (Fig. 6A; Supplemental Information A.3). Strikingly, all SAC genes, with the exception of *mad2*, showed lower Fano factors in the cytoplasm than in the nucleus (Fig. 6A, Supplemental Information Table A-I). Other low noise genes (*sep1* and *rpb1*) showed a similar or slightly higher Fano factor in the cytoplasm than in the nucleus, and high noise genes (*SPAC2H10.01* and *SPAC27D7.09c*) showed a considerably higher Fano factor in the cytoplasm than in the nucleus. These results suggest that for most SAC genes the Fano factor of their mRNA distribution is further lowered post-transcriptionally.

**Figure 6.**
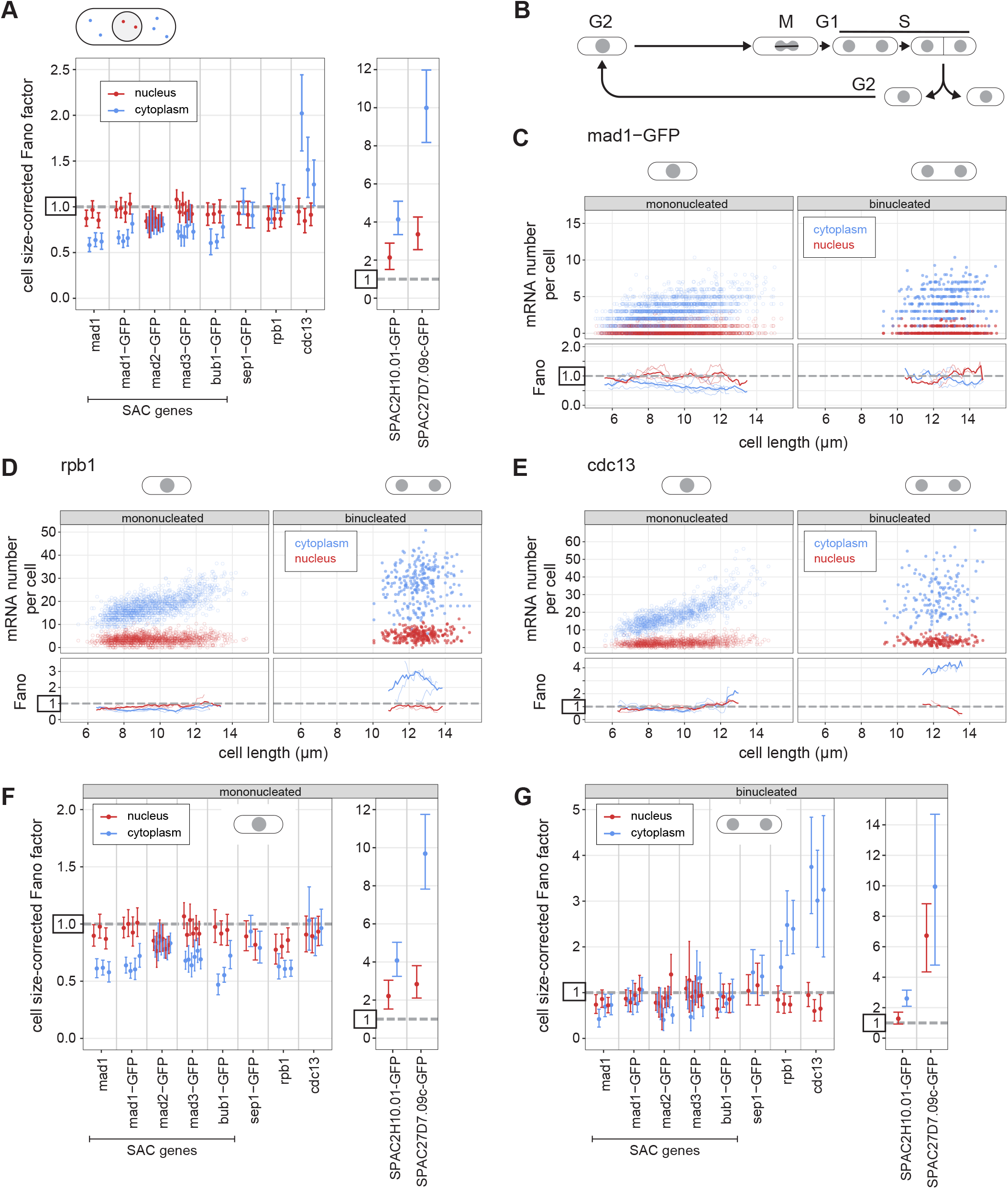
The Fano factor of the mRNA distribution can be lower in the cytoplasm than in the nucleus. **(A)** Cell size-corrected Fano factors and their 95 % confidence interval were calculated from smFISH data. Same experiments as in Fig. 2, but nuclei were segmented using DAPI staining, and the cell size-corrected Fano factor was calculated separately for mRNA localizing to the nucleus and localizing to the cytoplasm. Between 1and 6 replicate experiments for each genotype; the number of cells in single replicates ranged from 209 to 778. **(B)** Schematic of the *S. pombe* cell cycle. **(C-E)** Top: Scatter plot of cell length versus mRNA number in cytoplasm (blue) or nucleus (red). Mono- and binucleated cells are shown separately. Bottom: The Fano factor was determined in a sliding window spanning 1 µm of cell length. The Fano factors for single replicates are shown as thin lines; the Fano factor for the pooled data as thick line. See Fig. S8 for other SAC genes. **(F,G)** As in (A), but analyzed separately for mononucleated (F) and binucleated (G) cells.

These data with segmented nuclei also allowed us to distinguish mononucleated and binucleated cells (Fig. 6B-G; Supplemental Information A.4). In *S. pombe*, the nuclear envelope does not break down during cell division, and chromosome segregation is followed by splitting of the nucleus into two, and ultimately cell division (*75*). G1 phase in *S. pombe* is short and DNA replication occurs mostly prior to cell division during the binucleated stage (*76*). Thus, binucleated cells are in late mitosis, G1, or S phase, whereas almost all mononucleated cells are in G2 (Fig. 6B). We observed significant Fano factor differences between mononucleated and binucleated cells for almost all genes, with stronger differences in the cytoplasm than the nucleus for the low noise genes (Fig. 6C-G, S8, Supplemental Information, Table A-IV). In mononucleated cells, the *rpb1* gene shows a pattern very similar to the SAC genes with a sub-Poissonian distribution of mRNA numbers in the cytoplasm and a lower Fano factor in the cytoplasm than the nucleus (Fig. 6F). The Fano factor suppression between nucleus and cytoplasm for SAC genes went away during the binucleated stage, and *rpb1* and *cdc13* showed drastic increases in cytoplasmic (but not nuclear) Fano factor in binucleated cells (Fig. 6F,G). Overall, this most likely reflects dramatic changes in cellular physiology during cell division and DNA replication, which could impact mRNA synthesis, nuclear export, mRNA degradation, or possibly all of those (*77–83*). It also illustrates that it is important to minimize cell cycle effects when analyzing gene expression noise and its mechanistic basis (*43, 84–86*).

### The Fano factor of the *mad1* mRNA distribution is lower in the cytoplasm than at the transcription site or in the nucleoplasm

Our observations suggest that the Fano factor of SAC gene mRNA distributions is further lowered post-transcriptionally. To strengthen this conclusion, we additionally examined mRNA directly at the transcription start site (Fig. 7A). Using a similar strategy to one previously employed in *E. coli* (*87*), we labelled the transcription start site by integrating lac operator repeats into an intergenic region 3.3 kb and 6.4 kb away from *mad1* and *rpb1*, respectively, and expressing a Lac inhibitor- NLS-GFP fusion in these cells (Fig. 7A). This resulted in typically one GFP spot in the nucleus (Fig. 7B). Both mean mRNA numbers and cell size-corrected Fano factors remained highly similar to cells without the transcription site labelled (Fig. S9A), suggesting that the integration did not alter expression from the endogenous locus.

**Figure 7.**
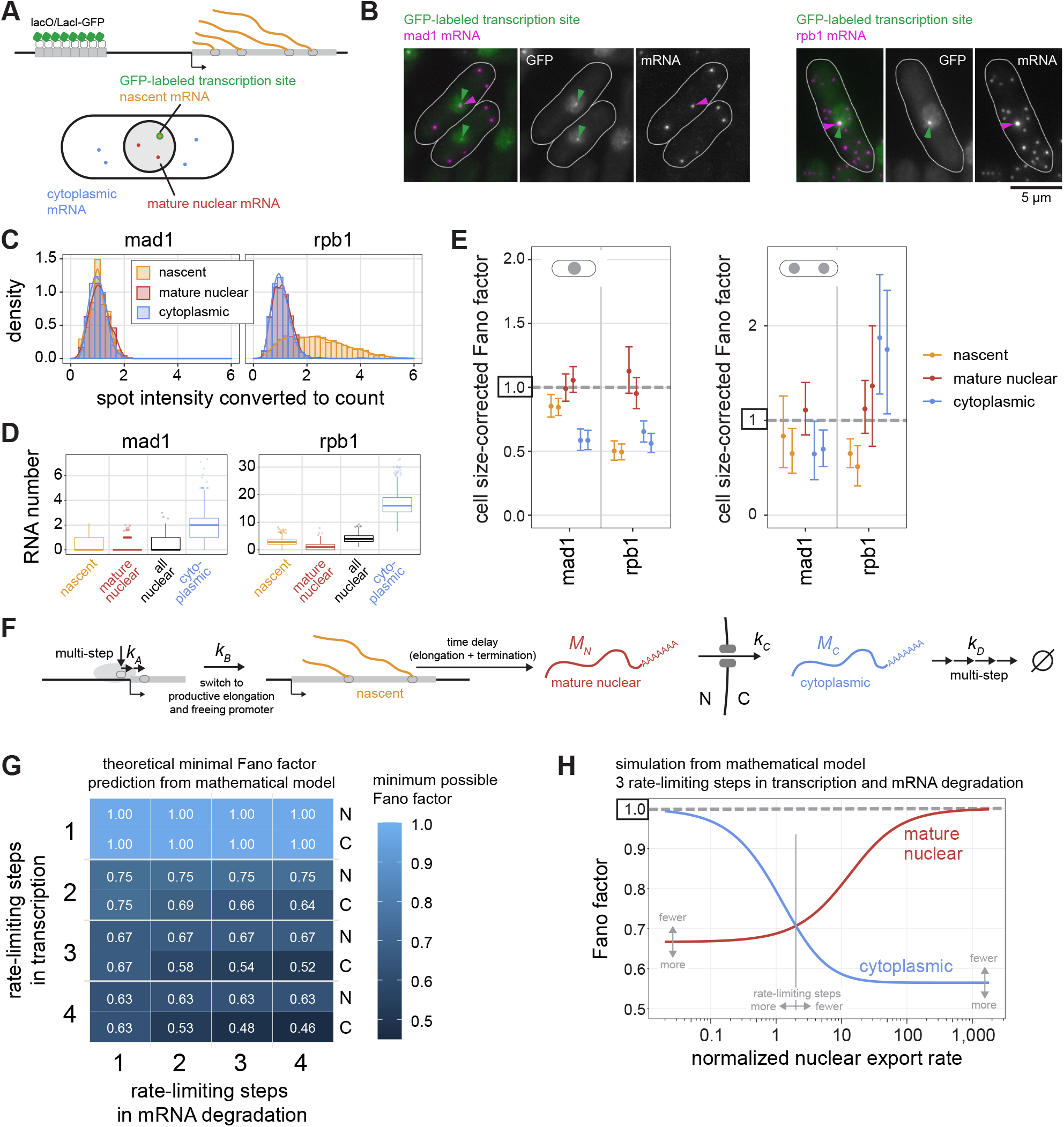
A sub-Poissonian mRNA distribution in the cytoplasm requires multiple rate-limiting steps in mRNA degradation and is modulated by nuclear export kinetics. **(A)** Diagram of transcription site labeling, and classifying mRNA into nascent (associated with the GFP-labeled gene), mature nuclear, or cytoplasmic. **(B)** Representative images of mad1 or rpb1 mRNA FISH in cells with the transcription site labeled by GFP. **(C)** Histogram and density distribution of FISH spot intensity at the transcription site (nascent), at other positions in the nucleus (mature nuclear), or in the cytoplasm. Data were normalized to the median of the spot intensity in the cytoplasm for each image. Pooled data from two independent experiments; n = 2,217 spots for mad1; n = 20,740 spots for rpb1. See Fig. S9 for the independent experiments. **(D)** Numbers of mRNA in the different compartments in mononucleated cells; n = 884 cells for mad1, n = 921 cells for rpb1. Boxplots show median and interquartile range; whiskers extend to values no further than 1.5 times the interquartile range from the first and third quartile, respectively. **(E)** Cell size-corrected Fano factors and their 95 % confidence interval were calculated from mRNA FISH data for nascent, mature nuclear, or cytoplasmic mRNA. Mono- and binucleated cells were analyzed separately. Two replicate experiments for each genotype; one of the experiments did not have enough binucleated cells with mature nuclear mad1 mRNA for a reliable estimate of the cell size-corrected Fano factor. The number of cells in single replicates ranged from 343 to 541 for mononucleated and 32 to 98 for binucleated cells. **(F)** Outline of the parameters in the mathematical model, including multiple rate-limiting steps in mRNA synthesis and degradation. **(G)** Analytical results from the mathematical model: minimum possible Fano factors in nucleus (N) and cytoplasm (C) depending on the number of rate-limiting steps in transcription and mRNA degradation. **(H)** Expectation for the nuclear and cytoplasmic Fano factors for a model with three rate-limiting steps in both transcription and mRNA degradation; *k_A_* = *k_B_* = 10; *k_D_* = 3. The x-axis shows the nuclear export rate *k_C_*, which is defined relative to the effective mRNA degradation rate. The nuclear export rate at which the Fano factors are equal scales inversely with the number of rate-limiting steps in transcription and mRNA degradation (Supplemental Information, Table B-XIII). Gray arrows illustrate in which direction the curves are expected to change for fewer or more rate-limiting steps in transcription and mRNA degradation.

We classified mRNAs in the direct vicinity of the GFP spot as nascent (Fig. 7A; S7B). Occasionally, a transcription site was just outside of the region that we had classified as nuclear based on DNA labelling and segmentation. In this case, we isotropically expanded the nucleus to include the transcription site (Fig. S7B) and adjusted our classification of nuclear and cytoplasmic mRNA accordingly. Excluding these cells from the analysis as an alternative method yielded similar results (compare Fig. 7E and Fig. S9B). For the highly expressed *rpb1* gene, this procedure captured all high-intensity mRNA spots as nascent (Fig. 7B,C; S9C), confirming the validity of the approach. This procedure then allowed us to distinguish nascent, mature nuclear, and cytoplasmic mRNAs (Fig. 7A). For both *mad1* and *rpb1*, the number of mature nuclear mRNAs was lower than the number at the transcription start site, indicating fast nuclear mRNA export (Fig. 7D), as has been previously found for budding yeast (*88*). Consequently, the number of all nuclear mRNAs is not a good proxy for the number of mature nuclear RNAs (Fig. 7D; Supplemental Information Table B-II).

The analysis of the cell size-corrected Fano factors in the different compartments showed that both *mad1* and *rpb1* are transcribed in a way that yields a sub-Poissonian distribution of mRNA numbers at the transcription site, even stronger for *rpb1* than for *mad1* (Fig. 7E; S9D; Supplemental Information A.2). The Fano factor of mature nuclear mRNA was close to 1. In agreement with our previous observations (Fig. 6F), the Fano factor of cytoplasmic *mad1* mRNA in mononucleated cells was clearly lower than either in the nucleoplasm or at the transcription site (Fig. 7E; Supplemental Information A.3). For *rpb1*, the Fano factor of cytoplasmic mRNA was lower than that of mature nuclear mRNA, but slightly larger than that at the transcription site. As we had seen before, suppression of the Fano factor in the cytoplasm for *mad1* disappeared in binucleated cells (Fig. 7E). The results were independent of whether we used cell length or cell volume as size parameter (Fig. S9E). Furthermore, the same trends were observed when binning cells by their size and analyzing the Fano factor of the mRNA distribution in cells of similar size (Fig. S9F). Overall, these results show that both these constitutively expressed genes show characteristics of transcription that lead to a sub-Poissonian distribution of mRNA numbers at the transcription start site. In addition, the results confirm that the Fano factor of the cytoplasmic mRNA distribution can be significantly lower than that in the nucleoplasm, or even at the transcription site.

### Multiple rate-limiting steps in transcription and degradation, combined with fast nuclear export, can explain the sub-Poissonian mRNA distributions

How can sub-Poissonian mRNA distributions be explained mechanistically? A simple, widely used model for constitutive gene expression assumes single rate-limiting steps for mRNA synthesis and degradation with exponential wait times between single synthesis or degradation events (*33–36*). Under these assumptions, the Fano factor of mature mRNA in both the nucleus and cytoplasm will be 1 (*38*). This may be consistent with data for some constitutive genes, but clearly not all.

To obtain a distribution of mRNA numbers that is narrower than the Poisson distribution, the wait times between single mRNA synthesis events need to be more homogeneous than in an exponential distribution (*89, 90*). Assuming multiple steps with similar timescales in mRNA synthesis, rather than just one rate-limiting step, will lead to such narrower than exponential wait times (*91–94*). Biologically, this assumption is well justified because RNA polymerase II (Pol II) binding to the promoter is known to be followed by several additional steps, such as promoter opening, promoter escape, promoter-proximal pausing and pause release (*95, 96*). We constructed a model of this type that also includes mRNA export from the nucleus to the cytoplasm and degradation of the mRNA in the cytoplasm (Fig. 7F). Degradation of mRNA in the cytoplasm is also known to involve multiple steps (*97, 98*), and it has been argued that assuming several rate-limiting steps is necessary to fit transcriptome-wide mRNA decay data of several budding yeast genes (*99*). Hence, the model includes the possibility of multiple rate-limiting steps in mRNA degradation.

We derived mathematical expressions for the Fano factor in steady-state conditions to determine the minimal Fano factors possible for this model (Fig. 7G and Supplemental Information). As expected, increasing the number of transcription initiation steps decreases the possible Fano factor and allows for Fano factors below 1 (Fig. 7G). Interestingly, increasing the number of rate-limiting steps in mRNA degradation allows for lower Fano factors in the cytoplasm than in the nucleus (Fig. 7G), fitting what we observed experimentally (Fig. 6,7). In this model, whether the cytoplasmic Fano factor is lower or higher than the nuclear Fano factor depends on the nuclear export rate (Fig. 7H). At slow nuclear export rates, the Fano factor in the cytoplasm is larger than that in the nucleus. Only at fast nuclear export rates will the cytoplasmic Fano factor become smaller than that in the nucleus (Fig. 7H). The threshold value at which the switch occurs becomes lower with an increasing number of rate-limiting steps in transcription and mRNA degradation (Fig. 7H; Supplemental Information Table B-XIII). Hence, our mathematical model shows that multiple rate-limiting steps in transcription and mRNA degradation, along with efficient nuclear export of mRNA, can explain both the sub-Poissonian mRNA distributions as well as the lower Fano factor in the cytoplasm than in the nucleus. Furthermore, nuclear export can be an efficient lever to tune cytoplasmic mRNA variability in this sub-Poisson regime.

### At least three rate-limiting steps in synthesis and mRNA degradation are required to explain all SAC gene data

To determine how many rate-limiting steps in transcription and mRNA degradation are required to explain the experimental data, we used Bayesian model selection. We found strong evidence that 3 or 4 rate-limiting steps in both transcription and mRNA degradation are required to explain the data for *mad1*, *mad1*-GFP and *bub1*-GFP (Supplemental Information Table B-XI). The expression of other genes (*mad2*-GFP, *mad3*-GFP, *rpb1*, *sep1*-GFP) could be equally well explained by other combinations of the number of rate-limiting steps, with the provision that there are at least 2 steps for transcription. For each gene, we then estimated the rate constants for promoter remodeling (*k­*), promoter freeing (*k_B_*), and nuclear export (*k_C_*) relative to the effective rate of cytoplasmic degradation, using a model with three rate-limiting steps in both transcription and mRNA degradation (Supplemental Information Table B-XII, Fig. B-1). This indicated differences between genes. The non-SAC genes, *rpb1* and *sep1*, were best fit with higher rates of promoter remodeling (*k_A_*); the two genes not showing a reduction of the Fano factor in the cytoplasm, *mad2* and *sep1*, were best fit with lower rates of nuclear export (*k_C_*) (Supplemental Information, Fig. B-2). Consistent with a lower nuclear export rate, *mad2* and *sep1* have a higher fraction of their mRNA in the nucleus (Fig. S8G), information that was not used in the fitting (Supplemental Information B.3). Altogether, our results imply that the sub-Poissonian mRNA distributions observed for some constitutively expressed genes can be biologically explained by multiple rate-limiting steps in transcription and mRNA degradation. Furthermore, our results indicate that constitutively expressed genes are not a homogenous group, but instead differ measurably in their expression characteristics with consequences on mRNA noise.

## Discussion

The expression characteristics of low noise genes have received much less attention than those of high noise genes. Here, we showed that the mRNA distribution of constitutively expressed, low noise genes can be narrower than a Poisson distribution, which establishes that the intrinsic “noise floor” of constitutively expressed genes is sub-Poissonian, not Poissonian. This reveals that low noise gene expression is more varied than previously thought and that constitutively expressed genes are not a homogenous group. Our findings imply that the widely used model for constitutive expression with single rate-limiting steps in transcription and mRNA degradation is an oversimplification. This knowledge is relevant because researchers have at times eliminated data from their analysis when the mRNA distributions appeared sub-Poissonian (*100*). Based on our results, this may have excluded valid and interesting data. We suggest that—in their most basic, unregulated form—the multi-step nature of transcription and mRNA degradation leads to sub-Poissonian noise. However, certain promoter architectures or added-on regulation, e.g. by a transcription factor or by an RNA-binding protein, can introduce additional intrinsic noise and thereby move the gene closer to a Poisson-type expression.

Our findings were surprising because prior reports of sub-Poissonian mRNA distributions are extremely sparse. In bacteria, one group has reported that expression from the *tetA* promoter results in a sub-Poissonian mRNA distribution (*101*), and that the wait times between synthesis events from this and two other bacterial promoters are more narrowly distributed than exponential, consistent with two or three rate-limiting steps in transcription (*101–103*). Others concluded that gene expression noise in bacteria is generally Poissonian or super-Poissonian (*5, 41, 104*). In eukaryotes, mRNA distributions of constitutively expressed genes have generally been reported to be Poissonian (*6, 39, 40*). However, some of these values were obtained without correcting for cell size, which may have made a sub-Poissonian distribution appear Poissonian (see Fig. S2A for an example). In addition, unrecognized cell cycle effects may have muddied the data. For example, the cytoplasmic mRNA distribution of *rpb1* (coding for the largest subunit of Pol II) has a Fano factor clearly below 1 in mononucleated *S. pombe* cells, but clearly above in binucleated (Fig. 7D), which averages to around 1 across the population (Fig. 6A)—and hence its expression was assumed Poissonian (*40*). Interestingly, one report provides evidence for a sub-Poissonian mRNA distribution of the orthologous *S. cerevisiae RPB1* gene (*105*). However, finite sample effects would need to be excluded for a formal conclusion. Furthermore, nascent mRNA distributions at the transcription site of budding yeast genes have been reported to be sub-Poissonian (*39, 106*). However, the transcription site is special in that it contains both full-length and shorter mRNAs. As a result, even if the time between synthesis events is exponentially distributed, the distribution of nascent mRNAs at the transcription site may appear sub-Poissonian (*91*).

Overall, although prior data from eukaryotes does not specifically support sub-Poissonian distributions, it does not exclude them either. Since we used one of the gold standards to assess mRNA numbers (*107*), which yielded solid evidence of sub-Poissonian distributions (Figs. 2,6,7; Supplemental Information), and since unrecognized biological or technical variability would only inflate the measured variability, we find it hard to escape the conclusion that the expression of constitutive genes can be sub-Poissonian.

Whether mRNA distributions are sub-Poissonian or Poissonian has at least two biological implications. For one, it makes a difference in noise strength. Whether this is functionally relevant, i.e. whether increasing sub-Poissonian to Poissonian expression lowers fitness, remains to be determined. Generally, noise characteristics of genes can be under natural selection (*22, 108*–*111*). It will be interesting to determine which classes of genes show sub-Poissonian expression and if and how this relates to their function. Secondly, the mRNA distributions provide a window into the gene expression process. A promoter transcribing with a single rate-limiting step must be fundamentally different from one exhibiting multiple rate-limiting steps. To what extent and how constitutive promoters differ from each other is still surprisingly little understood (*112*). Distinguishing sub-Poissonian and Poissonian low noise genes may help to classify promoters and identify their functionally important elements. It is worth noting, though, that the elements defining sub-Poissonian expression may not be neatly confined to the promoter (Fig. 5). This highlights the important role that chromatin context (*68, 113*) and non-coding features of the coding sequence (*114*) play in gene expression. Further, it remains a possibility that sub-Poissonian distributions are 16 brought about by negative feedback, as has been examined repeatedly in models (*115–120*). For the genes examined here, we are not aware of any known process establishing negative feedback, and we therefore favor multiple steps in transcription and mRNA degradation as the origin of their sub-Poissonian mRNA distributions.

Another relevant observation is that most SAC genes showing sub-Poissonian mRNA distributions also show a reduction of the Fano factor between nucleus and cytoplasm. Such a reduction of variance has been seen for bursty genes, where it was attributed to slow nuclear export (*11, 28, 29*). In contrast, our model predicts that the nuclear export rate needs to be sufficiently fast for the Fano factor in the cytoplasm to drop below the one in the nucleus (Fig. 7H). Our analysis also shows that the cytoplasmic Fano factor can become lower with an increasing number of rate-limiting steps in mRNA degradation (Fig. 7G,H), which seems at odds with published results that multiple rate-limiting steps in mRNA degradation can increase the Fano factor in the cytoplasm (*31*). The difference between these published results and ours is the underlying type of expression. Lowering of the cytoplasmic Fano factor by slow nuclear export or increase by multiple rate-limiting steps in mRNA degradation holds true for super-Poissonian expression (nuclear Fano factor larger than 1), but the relationships change in sub-Poissonian expression (nuclear Fano factor smaller than 1). Hence, the sub-Poissonian regime is not only quantitatively, but also qualitatively distinct, with its own rules.

Lastly, among the SAC genes we have studied, *mad2* is an interesting exception. It does show a sub-Poissonian mRNA distribution, but it does not show the Fano factor reduction between nucleus and cytoplasm seen for the other SAC genes (Fig. 6, S8). This suggests that its nuclear export or mRNA degradation characteristics differ qualitatively from that of the other SAC genes. Indeed, Bayesian inference suggested that the nuclear export rate of *mad2* is slower than that of other SAC genes (Supplemental Information Table B-XII). We speculate that this could be a consequence of co-translational assembly of the Mad1/Mad2 complex. The Mad1 and Mad2 proteins form an extremely tight 2:2 complex (*53, 54*). The Mad1 dimer at the core of this complex assembles co-translationally (*45, 121*). We consider it possible that the binding of Mad2 to the Mad1 dimer also needs to be co-translational. In this case, *mad2* mRNA may associate with Mad1 protein, which in turn may affect its export characteristics or the number of rate-limiting steps in its degradation.

In summary, we have established that the low noise regime of eukaryotic gene expression reaches lower than previously thought. This opens avenues to understand the underlying molecular differences and identify gene elements that minimize noise.

## Materials and Methods

### Tables

*S. pombe* strains are shown in Table S1, sgRNA sequences in Table S2, FISH probes in Table S3, and qPCR primers in Table S4.

### Strain construction

SAC genes *mad1*, *mad2*, *mad3*, and *bub1* were tagged with ymEGFP at the endogenous locus without inserting any other exogenous sequences (*45*). This was accomplished either by replacement of the counter-selectable rpl42-hphNT1 cassette in an rpl42::cyh^R^(sP56Q) strain (*122*) or by using CRISPR-mediated targeting (*123*). The *mad2*-ymEGFP strain contains a single, silent (AGG to AGA) PAM site mutation at amino acid position 173 of Mad2. The *mad3*-ymEGFP strain contains a single, silent (TTG to TTA) PAM site mutation at amino acid position 199 of Mad3. Yeast codon-optimized, monomeric enhanced GFP (ymEGFP) was derived from yEGFP (*124, 125*) by mutation of Alanine 206 to Arginine (A206R), which is expected to reduce dimerization (*126*). Tagging of *sep1*, *SPAC2H10.01*, *SPAC27D7.09c*, and *rad21* with ymEGFP was performed by conventional PCR-based gene targeting (*127*). Integration at the exogenous *wis1* and *leu1* loci used CRISPR-mediated targeting (CRISPR); sgRNA sequences are listed in Table S2.

For transcription site labeling, lacO repeats were integrated as described by Rohner and colleagues (*128*). Insertion close to *mad1* is between the adjacent *trz2* and *but1* genes (3.3 kb away from the *mad1* stop codon), insertion close to *rpb1* is between the adjacent *toa1* and *hrd3* genes (6.4 kb away from the *rpb1* start codon). We tried to be minimally disruptive to gene expression by choosing relatively large intergenic regions, not directly adjacent but sufficiently close to the genes of interest. A resistance cassette (hphMX4 from pAG32) was inserted at the respective locus, and then swapped for the lacO repeats and *LEU2* as resistance marker (from pSR13). The swap was facilitated by CRISPR/Cas9-targeting of the resistance cassette.

### Cell growth

YEA (yeast extract with adenine, 30 g/L glucose, 5 g/L yeast extract, 0.15 g/L adenine hemisulfate dihydrate) was used as rich medium; EMM (Edinburgh minimal medium, MP Biomedicals, 4110032) was used as minimal medium. Strains were thawed on YEA plates, and then grown for approximately 24 hr in liquid EMM, all at 30°C. When cultures were diluted to low densities, 50 % pre-conditioned medium (made by filtering EMM cultures) was added to EMM. L-Leucine (0.2 g/L) was added to EMM medium as needed for auxotrophic strains.

### Single-molecule mRNA fluorescence in-situ hybridization (smFISH)

Cultures were grown to ∼0.7-1.5 x 10^7^ cells/ml and 2 x 10^8^ cells were fixed with either 2 % or 4 % formaldehyde. After 30 min, formaldehyde was washed out with three washes of ice-cold Buffer B (1.2 M sorbitol, 100 mM potassium phosphate buffer pH 7.5). When not immediately continuing with the next step, cells were stored in 1 mL of Buffer B at 4°C. Fixed cells were resuspended in 1 mL spheroplast buffer (1.2 M sorbitol, 0.1 M potassium phosphate, 20 mM vanadyl ribonuclease complex [NEB S1402S], 20 µM beta-mercaptoethanol), and 1.2–5 µL 100T zymolyase (US Biological Z1005; 10 mg/mL) were added to digest the cell wall. Cells were kept at 30°C until the cell walls were sufficiently digested (determined by counting the fraction of cells that lysed when placed in deionized water). Typically, 30-50% of cells lysed was taken as evidence for sufficient digestion. Digestion was stopped by washing the cells three times with 1 mL of Buffer B. Next, cells were incubated for 20 min in 1 mL of 0.01 % Triton X-100 in 1x phosphate-buffered saline (PBS), followed by three more washes of Buffer B and one wash with 10 % formamide in 2x saline-sodium citrate (SSC) buffer (ThermoFisher AM9770). Cells were resuspended in the formamide/SSC solution and split evenly into two replicate samples for hybridization. For each sample, 3.75 pmol of Stellaris^TM^ RNA FISH probes (CAL Fluor red 610 probes targeting ymEGFP or cdc13, or Quasar 570 probes targeting mad1 or rpb1; Biosearch Technologies, LGC) were combined with 2 µL each salmon-sperm DNA (Life Technologies, 15632-011) and yeast tRNA (Thermo Fisher, AM7119) and diluted in Buffer F (20 % formamide, 10 mM sodium phosphate buffer pH 7.2) to a final volume of 50 µL. This mixture was heated to 95°C for 3 min, allowed to cool to room temperate, and added to an additional 50 µL of Buffer H (4x SSC buffer, 4 mg/mL acetylated BSA [Sigma, B8894], 20 mM vanadyl ribonuclease complex). Each sample was resuspended in 100 µL of this hybridization solution. When possible, as a positive control, one of the replicates of each sample was hybridized with probes for a higher abundance mRNA which would be expected to show FISH spots in all cells if the experiment was successful. After overnight incubation in the dark at 37°C, the hybridization solution was removed and cells were incubated for 6 min in 10 % formamide/2x SSC heated to 37°C, 6 min in 0.1 % Triton X-100/2x SSC, and finally 10 min in 1 µg/ml DAPI in 1x PBS. After DAPI staining, the cells were washed once with 1x PBS before final resuspension in 1x PBS. Cells were stored in the dark at 4°C until imaging.

For imaging, cells were mounted in SlowFade Diamond Antifade Mountant (Thermo Scientific, S36972) with #1.5 glass coverslips and RNase-free slides. Slides were imaged with a Zeiss AxioImager M1 equipped with Xcite Fire LED illumination (Excelitas), a Zeiss Plan FLUAR 100x/1.45 oil objective, and an ORCA-Flash4.0LT sCMOS camera (Hamamatsu). FISH optimized red or gold filters (Chroma 49306 and 49304, respectively) were used to image the FISH probes, while standard GFP, CFP, and DAPI filters were used to capture images of GFP-labeled transcription sites, cell boundaries, and nuclei, respectively. The images for each channel consisted of a 6 µm z-stack containing 31 images at 0.2 µm intervals.

### Image processing of smFISH

Images were dark noise-subtracted and flatfield-corrected. Cells were segmented in two dimensions using a custom FIJI macro based on using trainable WEKA segmentation (*129*) while nuclei were segmented in 3D dimensions from DAPI images using a custom FIJI macro adapted from https://github.com/haesleinhuepf/cca_benchmarking (Robert Haase, MPI-CBG, Dresden). Cell segmentation was manually corrected and cells missing nuclear segmentation, or whose nuclei were not fully contained within the image stack, were removed. Cell length was measured from the segmented cell outlines using the bounding box method. RNA FISH spots and GFP transcription site spots were detected with FISHquant v3a (*130*). FISH or GFP images were filtered with the default 3D_LoG filter (size = 5, sigma = 1). Spots were initially found using local maximum detection and an automatically determined minimum intensity threshold; the quality score filtering option was not used. The default setting was used for minimum distance between spots (160 nm). After point spread function (PSF) fitting, final thresholds were set manually for the PSF sigma xy, sigma z, and z position to exclude non-RNA or non-transcription site spots such as hot pixels. Spot detection accuracy was checked manually for a subset of cells from each image. Additional R scripts were used to classify each spot as nuclear or cytoplasmic based on the 3D nuclear DAPI segmentation.

In transcription-site labeling experiments, cells with a GFP spot in the cytoplasm, distant from the nucleus, were excluded from the analysis (only 3 cells across both strains and experiments). In addition, cells without a GFP spot or with more than two GFP spots per nucleus were excluded from the analysis. This removed 3 – 13 % of the mononucleated cells and 15 – 37 % of the binucleated cells. In the binucleated cells, this was typically due to the absence of a detectable GFP spot in one or both of the two nuclei. To identify nascent mRNA FISH spots, the 3D distance of each FISH spot to its nearest transcription site (TS) spot was determined. The distribution of distances was bimodal (Fig. S7B), and a distance cutoff was used to separate nascent from mature mRNA. The distance cutoff was calculated separately for each image generated by a clustering method that locates the minimum between the two distance distributions.

In cells where a transcription site was located just outside the region initially segmented as nuclear based on DNA staining, the segmentation of the nucleus was isotropically expanded to include the transcription site (NucSegmentMethod = 3DNucExpand). This seemed justified, since nuclei in cells where this was the case were on average slightly smaller than those in cells where this was not the case, suggesting that the initial nuclear segmentation had not captured the entire nucleus. Alternatively, these cells were excluded from the analysis (NucSegmentMethod = CF, cells filtered), which did not change the overall results or conclusions.

The number of mRNA molecules per cell were calculated with three different approaches. “Spot counts” are the number of FISH spots detected, not taking into account their intensity. “Intensity counts” were obtained by dividing the amplitude of each spot’s PSF by the median amplitude of the cytoplasmic spots in each image. Since for the genes in this study the vast majority of spots in the cytoplasm can be assumed to consist of single mRNA molecules, the result is a normalized intensity measure where a typical single-RNA FISH spot will have a value of 1 (Fig. S1C). These normalized intensities were summed to produce an intensity-based estimate of the number of mRNA molecules per cell. The third method (“hybrid counts”) combines features of both these methods: the number of mRNAs was calculated by partially converting the normalized intensity estimates into integers (Fig. S1). If a spot had a normalized intensity below the 95^th^ percentile of the cytoplasmic spots in that sample, it was assumed to contain one mRNA molecule. For brighter spots, the number of mRNA molecules per spot was set equal to the normalized intensity (Fig. S1C). The numbers for all spots were summed to obtain the number of mRNA molecules per cell. We consider “hybrid count” to be the most reliable method, since it ignores random intensity noise in “single mRNA” spots from differences in labeling efficiency, but counts more than one mRNA molecule when the intensity of a spot is clearly higher than the typical “single mRNA” spot (as is the case at *rpb1* transcription sites, for example). We have used the “spot count” method for comparison to the Marguerat group data, since we did not have intensity values available for their data.

### Size-corrected Fano factor

Cell-size corrected Fano factors were calculated as by Padovan-Merhar and colleagues (*58*), using linear regression to assess the influence of cell size. We either used cell length as a proxy for cell size, or we calculated volume from cell length and width, making the assumption that *S. pombe* cells are shaped like a cylinder with hemispheres at the two ends.

To determine the Fano factor in a sliding window across cell sizes, cells within a window of 1 µm cell length were combined, and mRNA variance, mean, and Fano factor were calculated. The window was moved by 0.1 µm across all cell sizes. For mono- and binucleated cells combined, data were retained when there were at least 50 cells in the window for pooled data, or 40 cells in the window for single replicates, for at least 20 consecutive windows. For mononucleated cells, these threshold numbers were: 40, 30, 10; for binucleated cells, they were 30, 20, 6.

### Analysis of published smFISH data

We re-analyzed smFISH data of *S. pombe* by the Marguerat group (*12, 40*), who had also determined both mRNA numbers and cell size. Genes that appeared cell cycle-regulated were excluded (*ace2, fkh2, mid2, SPAPB17E12.14c, SPAPB1E7.04c*). One dataset (wt.1) for *rpb1* from Sun *et al.* was excluded because 5 % of the cells lacked any mRNA, whereas the highest percentage of cells without *rpb1* mRNA in any other dataset was 0.6 %. A few cells across all datasets had highly unusual cell widths (smaller than 1.5 µm or larger than 4 µm) and were excluded. Cells with a length above 16 µm were excluded as well. The size-corrected Fano factor was calculated in the same way as for our data.

### Immunoblotting

Between 6 x 10^7^ and 1 x 10^8^ cells were washed with water, pelleted, resuspended in 500 µL 0.22 M NaOH, 0.12 M beta-mercaptoethanol, and incubated on ice for 15 min. Trichloroacetic acid was added to a final concentration of 6 % followed by another 10 min incubation on ice. The samples were pelleted, optionally washed with acetone, and either resuspended in 100 µl of 1x HU buffer pH 6.8 (4 M urea, 2.5 % SDS, 100 mM Tris-HCl, 0.5 mM EDTA, 0.005 % bromophenol blue, 10 % glycerol, 0.1 M DTT) or in 1.5x NuPage LDS sample buffer (ThermoFisher NP0007) with 5 % beta-mercaptoethanol. Samples were heated to 70°C for 7 min, and further lysed with glass beads (Sigma G8772) for 1–2 min at 30 s^-1^ in a Retsch MM 400 ball mill. Extracts were collected by centrifugation and incubated at 70°C for another 3 min. Samples were run on NuPAGE 4-12 % Bis-Tris polyacrylamide gels (Invitrogen) with MOPS running buffer and NuPAGE antioxidant. Proteins were transferred to Immobilon-P PVDF membrane using an Amersham Semi-Dry Transfer Unit and transfer buffer (39 mM glycine, 48 mM Tris-base, 5–10 % methanol) containing 0.01 % SDS and 1:1,000 NuPAGE antioxidant. The membrane was blocked with 4 % nonfat milk in TBS-T (150 mM NaCl, 20 mM TRIS, 0.05 % Tween-20) for 15–30 min at room temperature. Antibodies were also in 4 % nonfat milk in TBS-T, and membranes were washed with TBS-T between antibody incubation steps. Primary antibodies were: mouse anti-GFP (Roche, 11814460001, 0.4 µg/mL), rabbit anti-Cdc2 (Santa Cruz, SC-53, 0.2 µg/mL), rabbit anti-Mad1 (*44*) (2 µg/mL), rabbit anti-Mad2 (*131*), rabbit anti-Mad3 (*132*) (serum at 1:500), rabbit anti-Bub1 (peptide: CKPKAGGPGRRRSSN, serum at 1:2,000); secondary antibodies were: goat anti-mouse HRPO or goat anti-rabbit HRPO (Dianova, 115-035-003 and 111-035-003, at 1:10,000). For imaging, membranes were incubated SuperSignal West Dura Extended Duration Substrate or SuperSignal West Pico PLUS Chemiluminescent Substrate (ThermoFisher) and chemiluminescence was recorded with a ChemiDoc XRS+ (Bio-Rad). Protein expression was quantified relative to the control sample using the control sample dilution series and Cdc2 as a loading control.

### Quantitative PCR

RNA extraction, cDNA library preparation, and quantitative PCR (qPCR) were performed as previously described (*45*). Briefly, cells were grown in either EMM (TATA-box mutant experiments), EMM with 0.2 g/L leucine (one *mad1* deletion strain), or YEA (all other) to concentrations of 0.7-1.5 x 10^7^ cells/ml and then flash frozen in liquid nitrogen. RNA extraction was followed by DNase-treatment and SuperScript IV reverse transcription with oligo d(T)_20_ primers. *act1* and *cdc2* were used as reference genes in all qPCRs.

### RACE-PCR

Cells were grown in YEA and snap frozen in liquid nitrogen (∼1 x 10^8^ cells). RNA was extracted using acidic phenol chloroform and treated with DNase (Roche DNase I) to remove any remaining DNA contamination. For 3’ UTR sequencing, DNase-treated RNA was reverse transcribed with the FirstChoice RLM-RACE kit (Invitrogen AM1700). Either the FirstChoice RLM-RACE kit with SuperScript IV (Invitrogen) or the GeneRacer kit (Invitrogen L150201) with SuperScript III (Invitrogen) was used to prepare the cDNA for 5’ UTR sequencing. cDNA was amplified with two rounds of nested PCR targeting the gene of interest. The PCR product was inserted into a pBlueScript vector using Gibson assembly (NEB, E2611) and transformed into NEB 5-alpha competent *E. coli* (NEB, C2992). Vectors were sequenced by Sanger sequencing. Sequences were discarded if the UTR appeared to begin (5’ UTR) or end (3’ UTR) in the coding sequence of the gene. When possible, to limit the analysis to mature mRNA, sequences were checked for the presence of introns and discarded if introns were found.

### SAC protein-GFP imaging in live cells

Cells were grown in EMM or EMM with 0.2 g/L leucine to between 4 x 10^6^ and 1.5 x 10^7^ cells/ml and mounted in Ibidi chambers (8-well, 80827) coated with lectin (Sigma, L1395, at 50 µg/mL). Cells were imaged on a DeltaVision Elite microscope equipped with a PCO edge sCMOS camera. Imaging was performed with an Olympus 60x/1.42 Plan APO oil objective and GFP filters. Images were taken as 7.2 µm z-stacks with 0.1 µm step size. To bleach autofluorescence, an initial GFP image stack was acquired with 0.1 sec exposures and discarded, before the main GFP image stack was acquired using 0.25-0.40 sec exposures (depending on the protein being imaged). GFP images were deconvolved using SoftWoRx software with two cycles of the ratio method (conservative) and a camera intensity offset of 50. Representative regions of cells were selected and single image slices from the z-stack were extracted at approximately the midplane of the cells. Image intensity was further scaled in ImageJ and Photoshop, being consistent for each GFP-tagged protein.

### GO enrichment analysis

GO-slim enrichment analysis was performed with the PANTHER overrepresentation test (http://www.pantherdb.org (*133*)) (released 2022-02-02), using the GO Ontology database (DOI:10.5281/zenodo.6399963; released 2022-03-22). From the data reported by Thodberg and colleagues (*61*) only transcripts of non-mitochondrial protein-coding genes were kept. For genes with multiple TSSs, only the TSS with the highest pooled TPM value was retained. Start codon positions were downloaded from Pombase (*134*) on 2022-06-22. All genes for which the TSS was located downstream of the currently annotated start codon were excluded. This resulted in a final list of 4,648 genes, which was used as reference data for the GO enrichment analysis. Of those genes, 296 had a 5’ UTR of 15 nucleotides or shorter, and 451 had a 5’ UTR of 20 nucleotides or shorter. Only the most specific subclass from the GO-slim analysis is shown. Fisher’s exact test with FDR correction was used to identify statistically significant results.

## Acknowledgments

We thank Andrea Ciliberto and Peter Swain for comments on the manuscript, Eric Esposito, Anupreet Kour, Anna Makarov, and Jessie Rogers for contributing experimental data, James Holehouse and Svitlana Braichenko for data processing and useful discussions, Tatiana Boluarte for constructing yeast strains, Hunter Haynie, and Haoyun Yang for preliminary experiments, Erod Keaton Baybay for help in establishing the image analysis pipeline, Yimeng Liu for implementing nuclear thresholding, Ben Unruh and Shihoko Kojima for support in running qPCRs, Samir Chethan for help with image quantification, and Daniel Zenklusen for sharing their yeast FISH protocol. Research reported in this publication was supported by NIGMS (NIH) under award number R35GM119723 (DEW, SH) and R35GM148351 (AS), by NSF grant ECCS-1711548 (AS), by the Leverhulme Trust RPG-2020-327 (RG), and by a Dean’s Discovery Fund of Virginia Tech’s College of Science (DEW, AS, SH).

## Author contributions

Conceptualization, DEW and SH; Software, DEW, RG and AS; Formal analysis, DEW and RG; Investigation, DEW; Writing–Original Draft, SH; Writing–Review & Editing, DEW, RG and AS; Visualization, DEW, RG and SH; Supervision, SH; Funding acquisition: SH, RG and AS.

