## Supplemental text + Tables S1-S4 for "The minimal intrinsic stochasticity of constitutively expressed eukaryotic genes is sub-Poissonian"

#### Supplementary Information

Table of Contents

Section A: Statistical tests of size-corrected Fano factors 2

A.1 Statistical methodology 2

A.2 Comparison of hierarchical bootstrapping with bootstrapping from pooled data 2

A.3 Comparison of Fano factors in cellular compartments 4

A.4 Comparison of Fano factors between mononucleated and binucleated cells 6

A.5 Comparison of Fano factors between endogenous and exogenous locus and with different coding sequences 7

Section B: Modeling and Inference 8

B.1 Model specification 8

B.2 General properties of the model 9

B.3 Bayesian model selection and parameter inference 11

B.4 Testing the ansatz of Poisson fluctuations 21

Tables S1 – S4 23

### Section A: Statistical tests of size-corrected Fano factors

#### A.1 Statistical methodology

For statistical conclusions on the size-corrected Fano factors, we used a hierarchical bootstrapping approach. Experimental replicates (shown separately in the main figures) and cells within replicates were resampled with replacement. From this, the statistic of interest (average Fano factor across experimental replicates, difference between genotypes, average difference in Fano factor between mono- and binucleated cells, or average difference in Fano factor between cellular compartments) was calculated. To estimate the statistic's distribution, 5,000 bootstrap replicates were performed, and the mean and 95 % confidence interval (2.5$^{th}$ to 97.5$^{th}$ percentile) were calculated from the bootstrap results. Up to 0.5 % of bootstrap replicates were allowed to fail to calculate (i.e., returned NaN), and these were discarded before calculating the mean and confidence interval. If > 0.5 % of bootstrap replicates failed, no statistical results were reported for that analysis. The latter was only the case for mature nuclear mRNA from *mad1* in binucleated cells (TS-labelled data), due to a low cell number in one replicate. A genotype was considered to have a size-corrected RNA distribution significantly different from Poisson if the bootstrap 95 % confidence interval excluded 1, and differences were considered significant if the confidence interval excluded 0.

#### A.2 Comparison of hierarchical bootstrapping with bootstrapping from pooled data

As a comparison to hierarchical bootstrapping, cells from replicate experiments were combined and the cell-size corrected Fano factor calculated from the pooled set of cells. This yielded very similar results to the hierarchical bootstrapping results (Fig. A-I, A-III), except for some genes in published data (*1*, *2*) with a larger variation between experimental replicates (Fig. A-II).

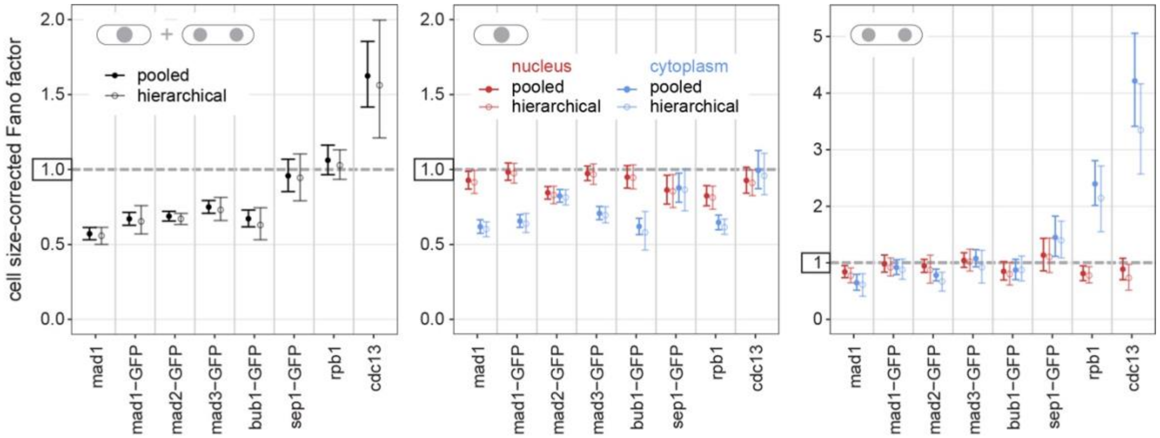


*Figure A-I: Fano factors from pooled and hierarchical analysis (non-TS labelled)*

Related to Fig. 2 and Fig. 6F and G.

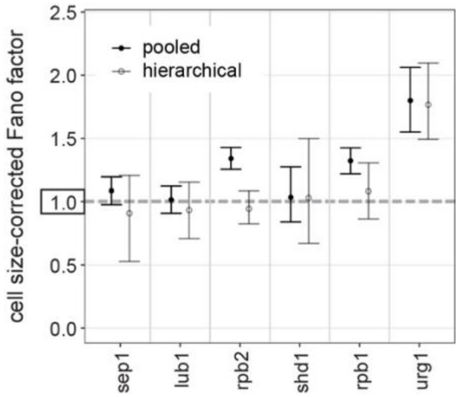


*Figure A-II: Fano factors from pooled and hierarchical analysis of Marguerat group data*

Related to Fig. 2D. Only genes with more than one experimental replicate are shown.

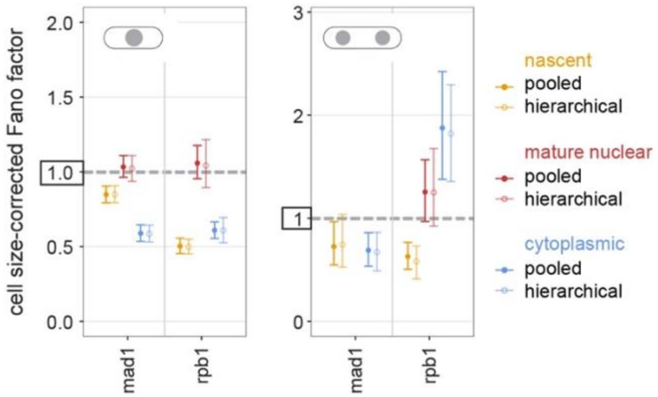


*Figure A-III: Fano factors from pooled and hierarchical analysis (TS-labelled)*

Related to Fig. 7E.

#### A.3 Comparison of Fano factors in cellular compartments

We tested if size-corrected Fano factors differ between cellular compartments using hierarchical bootstrapping. Mean and 95 % confidence intervals from bootstrapping the mean difference between compartments are provided. Confidence intervals that exclude 0 are interpreted as significantly different Fano factors between compartments.

*Table A-I: Comparison between nuclear and cytoplasmic Fano factors (non-TS labelled)*

Comparing nuclear and cytoplasmic Fano factors in experiments without transcription-site labelling (see Fig. 6 and A-I). A positive mean Fano factor difference indicates a larger Fano factor in the cytoplasm than the nucleus, while a negative mean Fano factor difference indicates a smaller Fano factor in the cytoplasm than the nucleus.

| Gene | Nuclei Count | Mean Fano Factor Difference | Lower 95 % Confidence Interval | Upper 95 % Confidence Interval | Effect |
|---|---|---|---|---|---|
| mad1 | 1 | -0.31 | -0.39 | -0.23 | smaller |
| mad1 | 2 | -0.16 | -0.38 | 0.06 | same |
| mad1 | all | -0.29 | -0.37 | -0.20 | smaller |
| mad1-GFP | 1 | -0.34 | -0.42 | -0.25 | smaller |
| mad1-GFP | 2 | -0.04 | -0.25 | 0.18 | same |
| mad1-GFP | all | -0.29 | -0.37 | -0.20 | smaller |
| mad2-GFP | 1 | -0.02 | -0.09 | 0.07 | same |
| mad2-GFP | 2 | -0.20 | -0.52 | 0.05 | same |
| mad2-GFP | all | -0.04 | -0.10 | 0.03 | same |
| mad3-GFP | 1 | -0.27 | -0.37 | -0.18 | smaller |
| mad3-GFP | 2 | -0.11 | -0.53 | 0.26 | same |
| mad3-GFP | all | -0.22 | -0.33 | -0.12 | smaller |
| bub1-GFP | 1 | -0.37 | -0.52 | -0.20 | smaller |
| bub1-GFP | 2 | 0.07 | -0.25 | 0.42 | same |
| bub1-GFP | all | -0.26 | -0.38 | -0.13 | smaller |
| sep1-GFP | 1 | 0.01 | -0.13 | 0.15 | same |
| sep1-GFP | 2 | 0.29 | -0.11 | 0.69 | same |
| sep1-GFP | all | 0.06 | -0.11 | 0.22 | same |
| rpb1 | 1 | -0.20 | -0.29 | -0.10 | smaller |
| rpb1 | 2 | 1.37 | 0.72 | 1.99 | larger |
| rpb1 | all | 0.18 | 0.05 | 0.30 | larger |
| cdc13 | 1 | 0.05 | -0.10 | 0.22 | same |
| cdc13 | 2 | 2.62 | 1.90 | 3.37 | larger |
| cdc13 | all | 0.66 | 0.30 | 1.09 | larger |
| SPAC2H10.01-GFP | 1 | 1.88 | 0.79 | 3.05 | larger |
| SPAC2H10.01-GFP | 2 | 1.31 | 0.61 | 1.95 | larger |
| SPAC2H10.01-GFP | all | 1.97 | 0.91 | 3.05 | larger |
| SPAC27D7.09c-GFP | 1 | 6.84 | 4.82 | 9.03 | larger |
| SPAC27D7.09c-GFP | 2 | 3.19 | -1.93 | 7.89 | same |
| SPAC27D7.09c-GFP | all | 6.63 | 4.74 | 8.67 | larger |

*Table A-II: Comparison between nuclear and cytoplasmic Fano factors at the exogenous wis1 locus*

Comparing nuclear and cytoplasmic Fano factors at the exogenous wis1 locus (see Fig. 5E). A positive mean Fano factor difference indicates a larger Fano factor in the cytoplasm than the nucleus, while a negative mean Fano factor difference indicates a smaller Fano factor in the cytoplasm than the nucleus. Change observed in mononucleated cells (as opposed to mononucleated and binucleated cells pooled = all) is indicated in the second effect column, without listing the mean Fano factor difference and confidence intervals.

| Gene | Mean Fano Factor Difference | Lower 95 % Confidence Interval | Upper 95 % Confidence Interval | Effect all | Effect mono-nucl. |
|---|---|---|---|---|---|
| mad2 endog | -0.04 | -0.11 | 0.03 | same | same |
| mad3 endog | -0.23 | -0.33 | -0.12 | smaller | smaller |
| rad21 endog | 6.74 | 5.82 | 7.71 | larger | larger |
| mad2 exog | 0.02 | -0.12 | 0.16 | same | same |
| Pmad2-rad21 exog | 0.24 | 0.02 | 0.46 | larger | same |
| Pmad2-nmt1 exog | 0.10 | -0.06 | 0.26 | same | same |
| mad3 exog | 0.03 | -0.12 | 0.17 | same | same |
| Pmad3-rad21 exog | 0.50 | 0.27 | 0.73 | larger | same |
| Pmad3-nmt1 exog | 0.26 | 0.03 | 0.49 | larger | same |

*Table A-III: Comparison of Fano factors in different compartments (TS-labelled)*

Comparing nascent, mature nuclear, and cytoplasmic Fano factors in experiments with transcription-site labelling (see Fig. 7 and A-III). A positive mean Fano factor difference indicates a larger Fano factor in compartment 2 than in compartment 1, while a negative mean Fano factor difference indicates a smaller Fano factor in compartment 2 than in compartment 1.

| Gene | Nuclei Count | Compartment 1 | Compartment 2 | Mean Fano Factor Difference | Lower 95 % Confidence Interval | Upper 95 % Confidence Interval | Effect |
|---|---|---|---|---|---|---|---|
| mad1 | 1 | Nascent | Mature Nuclear | 0.17 | 0.07 | 0.28 | larger |
| mad1 | 1 | Nascent | Cytoplasm | -0.26 | -0.34 | -0.18 | smaller |
| mad1 | 1 | Mature Nuclear | Cytoplasm | -0.44 | -0.54 | -0.33 | smaller |
| mad1 | 2 | Nascent | Mature Nuclear | N/A | N/A | N/A | N/A |
| mad1 | 2 | Nascent | Cytoplasm | -0.08 | -0.45 | 0.23 | same |
| mad1 | 2 | Mature Nuclear | Cytoplasm | N/A | N/A | N/A | N/A |
| mad1 | all | Nascent | Mature Nuclear | 0.20 | 0.07 | 0.31 | larger |
| mad1 | all | Nascent | Cytoplasm | -0.23 | -0.32 | -0.15 | smaller |
| mad1 | all | Mature Nuclear | Cytoplasm | -0.42 | -0.52 | -0.32 | smaller |
| rpb1 | 1 | Nascent | Mature Nuclear | 0.54 | 0.40 | 0.71 | larger |
| rpb1 | 1 | Nascent | Cytoplasm | 0.11 | 0.02 | 0.21 | larger |
| rpb1 | 1 | Mature Nuclear | Cytoplasm | -0.43 | -0.57 | -0.31 | smaller |
| rpb1 | 2 | Nascent | Mature Nuclear | 0.67 | 0.29 | 1.21 | larger |
| rpb1 | 2 | Nascent | Cytoplasm | 1.23 | 0.75 | 1.72 | larger |
| rpb1 | 2 | Mature Nuclear | Cytoplasm | 0.57 | -0.14 | 1.19 | same |
| rpb1 | all | Nascent | Mature Nuclear | 0.53 | 0.42 | 0.65 | larger |
| rpb1 | all | Nascent | Cytoplasm | 0.48 | 0.28 | 0.70 | larger |
| rpb1 | all | Mature Nuclear | Cytoplasm | -0.06 | -0.28 | 0.17 | same |

#### A.4 Comparison of Fano factors between mononucleated and binucleated cells

We tested if size-corrected Fano factors differ between mono- and binucleated cells using hierarchical bootstrapping. Mean and 95 % confidence intervals from bootstrapping the mean difference are provided. Confidence intervals that exclude 0 are interpreted as a significant difference in Fano factors.

*Table A-IV: Comparison of Fano factors between binucleated and mononucleated cells (non-TS labelled)*

Comparing Fano factors from mono- and binucleated cells in experiments without transcription-site labelling (see Fig. 6 and A-I). A positive mean Fano factor difference indicates a larger Fano factor in binucleated than mononucleated cells, while a negative mean Fano factor difference indicates a smaller Fano factor in binucleated than mononucleated cells.

| Gene | Compartment | Mean Fano Factor Difference | Lower 95 % Confidence Interval | Upper 95 % Confidence Interval | Effect |
|---|---|---|---|---|---|
| mad1 | Nucleus | -0.14 | -0.27 | -0.02 | smaller |
| mad1 | Cytoplasm | 0.01 | -0.21 | 0.22 | same |
| mad1-GFP | Nucleus | -0.06 | -0.22 | 0.12 | same |
| mad1-GFP | Cytoplasm | 0.24 | 0.08 | 0.41 | larger |
| mad2-GFP | Nucleus | 0.05 | -0.22 | 0.33 | same |
| mad2-GFP | Cytoplasm | -0.14 | -0.34 | 0.04 | same |
| mad3-GFP | Nucleus | 0.06 | -0.13 | 0.31 | same |
| mad3-GFP | Cytoplasm | 0.22 | -0.04 | 0.52 | same |
| bub1-GFP | Nucleus | -0.14 | -0.37 | 0.09 | same |
| bub1-GFP | Cytoplasm | 0.29 | 0.06 | 0.57 | larger |
| sep1-GFP | Nucleus | 0.25 | -0.06 | 0.61 | same |
| sep1-GFP | Cytoplasm | 0.53 | 0.21 | 0.88 | larger |
| rpb1 | Nucleus | -0.04 | -0.20 | 0.15 | same |
| rpb1 | Cytoplasm | 1.53 | 0.91 | 2.11 | larger |
| cdc13 | Nucleus | -0.18 | -0.41 | 0.08 | same |
| cdc13 | Cytoplasm | 2.38 | 1.61 | 3.19 | larger |
| SPAC2H10.01-GFP | Nucleus | -0.93 | -1.80 | -0.13 | smaller |
| SPAC2H10.01-GFP | Cytoplasm | -1.47 | -2.57 | -0.46 | smaller |
| SPAC27D7.09c-GFP | Nucleus | 3.87 | 1.08 | 6.16 | larger |
| SPAC27D7.09c-GFP | Cytoplasm | 0.19 | -5.79 | 5.38 | same |

*Table A-V: Comparison of Fano factors between binucleated and mononucleated cells (TS-labelled)*

Comparing Fano factors from mono- and binucleated cells in experiments with transcription-site labelling (see Fig. 7 and A-III). A positive mean Fano factor difference indicates a larger Fano factor in binucleated than mononucleated cells.

| **Gene** | **Compartment** | **Mean Fano Factor Difference** | **Lower 95 % Confidence Interval** | **Upper 95 % Confidence Interval** | **Effect** |
|---|---|---|---|---|---|
| mad1 | Nascent | -0.11 | -0.32 | 0.18 | same |
| mad1 | Mature Nuclear | N/A | N/A | N/A | N/A |
| mad1 | Cytoplasm | 0.08 | -0.12 | 0.29 | same |
| rpb1 | Nascent | 0.08 | -0.09 | 0.24 | same |
| rpb1 | Mature Nuclear | 0.21 | -0.18 | 0.75 | same |
| rpb1 | Cytoplasm | 1.21 | 0.74 | 1.68 | larger |

#### A.5 Comparison of Fano factors between endogenous and exogenous locus and with different coding sequences

We tested if size-corrected Fano factors for *mad2* and *mad3* differ between endogenous and exogenous locus, and after exchanging the coding sequence, using hierarchical bootstrapping. Mean and 95 % confidence intervals from bootstrapping the mean difference are provided. Confidence intervals that exclude 0 are interpreted as a significant difference in Fano factors.

*Table A-VI: Comparison of Fano factors for mad2 and mad3 between endogenous and exogenous locus and with different coding sequences*

Comparing Fano factors for *mad2* and *mad3* between endogenous and exogenous locus, and at the exogenous locus after changing the coding sequence (see Fig. 5). A positive mean Fano factor difference indicates a larger Fano factor in Genotype 2 than Genotype 1. Change observed in mononucleated cells (as opposed to mononucleated and binucleated cells pooled = all) is indicated in the second Effect column, without listing the mean Fano factor difference and confidence intervals.

| **Genotype 1** | **Genotype 2** | **Compart-ment** | **Mean Fano Factor Difference** | **Lower 95 % Confidence Interval** | **Upper 95 % Confidence Interval** | **Effect all** | **Effect mono-nucl.** |
|---|---|---|---|---|---|---|---|
| mad2 endog | mad2 exog | Nucleus | 0.02 | -0.07 | 0.10 | same | same |
| mad2 endog | mad2 exog | Cytoplasm | 0.08 | -0.05 | 0.21 | same | same |
| mad2 exog | Pmad2-**rad21** exog | Nucleus | 0.12 | -0.04 | 0.29 | same | same |
| mad2 exog | Pmad2-**rad21** exog | Cytoplasm | 0.34 | 0.14 | 0.55 | larger | larger |
| mad2 exog | Pmad2-**nmt1** exog | Nucleus | 0.10 | -0.03 | 0.22 | same | same |
| mad2 exog | Pmad2-**nmt1** exog | Cytoplasm | 0.17 | 0.00 | 0.35 | same | same |
| mad3 endog | mad3 exog | Nucleus | 0.00 | -0.15 | 0.16 | same | same |
| mad3 endog | mad3 exog | Cytoplasm | 0.26 | 0.14 | 0.38 | larger | larger |
| mad3 exog | Pmad3-**rad21** exog | Nucleus | -0.07 | -0.26 | 0.11 | same | same |
| mad3 exog | Pmad3-**rad21** exog | Cytoplasm | 0.39 | 0.17 | 0.62 | larger | larger |
| mad3 exog | Pmad3-**nmt1** exog | Nucleus | -0.07 | -0.27 | 0.12 | same | same |
| mad3 exog | Pmad3-**nmt1** exog | Cytoplasm | 0.16 | -0.04 | 0.36 | same | larger |

#### Section B: Modeling and Inference

##### B.1 Model specification

Transcription is modelled by the set of (effective) reactions:

$$U_1 \rightarrow U_2 \rightarrow \cdots \rightarrow U_S \rightarrow U_1 + N$$

$$N \Rightarrow M_N$$

where $N$ is the nascent RNA and $M_N$ is the mature nuclear mRNA. We consider $S$ gene states, $U_i$, where $i$ = 1, …, $S$. The free promoter state $U_1$ denotes the state in which no RNA polymerase (RNAP) is bound to the promoter. Once RNAP binds the free promoter, a closed complex between the two is formed ($U_2$). Downstream steps ($U_3$ to $U_{S-1}$) may include several long-lived intermediate states related to promoter opening and escape (*3*–*8*). The last state $U_S$ is a promoter-proximal paused state; release from this state leads to the clearing of the promoter for new RNAP binding and the beginning of elongation of the nascent transcript (*9*). This is modelled by the reaction $U_S \rightarrow U_1 + N$. After a time delay, elongation and termination finish, leading to mature nuclear mRNA $M_N$. The rate of switching from $U_i \rightarrow U_{i+1}$ is given by $k_A$, and the rate of initiating nascent mRNA production, $U_S \rightarrow U_1 + N$, is given by $k_B$. The time for the nascent RNA $(N)$ to become mature mRNA $(M_N)$ is considered fixed at some value $T$, i.e. the reaction $N \Rightarrow M_N$ is a delayed reaction (denoted by the double right arrow). The fixed delay (deterministic elongation and termination) can be derived from a microscopic stochastic model of RNAP movement along the DNA template (*10*), and deterministic elongation has been observed in budding yeast (*11*). Similar models have been proposed (*6*, *12*–*14*). Note that in this model there is no transcriptionally inactive state in the sense that each state is associated with a particular stage of the transcription process.

Nuclear export is modelled by the reaction

$$M_N \rightarrow M_C^1$$

with rate $k_C$. This process is modelled by a single rate-limiting step. $M_C^1$ is to be interpreted as the mRNA upon its entrance into the cytoplasm.

Degradation of mRNA in the cytoplasm is modelled by the chain of reactions

$$M_C^1 \rightarrow M_C^2 \rightarrow \cdots \rightarrow M_C^R \rightarrow \emptyset$$

Cytoplasmic mRNA degradation in eukaryotes is a complex multi-step process (*15*, *16*) that we simplify to the chain of reactions above. We are not aware of specific data for the number of rate-limiting steps *R* in the degradation process. In Deneke et al., 2013 (*16*), it was shown that choosing R = 5 can fit data in yeast well, though other values may provide an equally good fit. The rate of switching from $M_C^i \rightarrow M_C^{i+1}$ is given by $k_D$. Note that the measured cytoplasmic mRNA is assumed to be the sum of cytoplasmic mRNA at any stage of its cytoplasmic lifecycle, i.e. $M_C = \sum_{i=1}^{R} M_C^i$.

##### B.2 General properties of the model

The model is characterized by four rate constants $k_A$, $k_B$, $k_C$ *and* $k_D$, the number of rate-limiting steps in the initiation process *(S)*, and the number of rate-limiting steps in cytoplasmic degradation *(R)*.

The mean time between two successive nascent mRNA production events is given by:

$$T_{init} = \frac{S-1}{k_A} + \frac{1}{k_B}$$

The mean time for nuclear export is:

$$T_{expt} = \frac{1}{k_C}$$

The mean time for cytoplasmic degradation is:

$$T_{degr} = \frac{R}{k_D}$$

The chemical master equation of the model is difficult to solve exactly, even in steady-state conditions. This is not unexpected since exact solutions for the joint distributions of molecule numbers are only known for a handful of special cases (*17*). However, because all reactions are first order, the propensities of the chemical master equation are linear in the molecule numbers. This implies that the means, variances and covariances of all species can be obtained exactly in closed-form using the linear-noise approximation (*17*). The matrix form of this method (*18*) is particularly useful because it makes computations using a computer algebra system such as Mathematica easy for any number of species (which can be arbitrarily large for our model depending on the values of *S* and *R*).

As in the telegraph model of gene expression (*19*), in steady-state conditions the moments of nuclear and cytoplasmic mRNA are functions not of the absolute values of the rate constants but rather of the rate constants normalized by the degradation rate, i.e. $k_A / k_D$, $k_B / k_D$, $k_C / k_D$. Hence, without loss of generality we set $k_D$ = R such that $T_{degr}$ is fixed to unity; in other words, this choice non-dimensionalizes time by dividing it by the mean time it takes for a cytoplasmic mRNA to decay.

It is straightforward to derive the first-order moments of nuclear and cytoplasmic mRNA numbers per gene copy. These are respectively given by: $\mu_N = \frac{k_A k_B}{k_C(k_A + k_B(S-1))}$ and $\mu_C = \frac{k_A k_B}{k_A + k_B(S-1)}$.

The closed-form solutions for the second-order moments of mRNA numbers were obtained using Mathematica. They are very complex and hence we do not show them here. Instead, we state properties of the stochastic model that follow from an analysis of these solutions:

For all parameters, the Fano factors of nuclear mRNA and of cytoplasmic mRNA are found to be less than or equal to 1. For $S$ = 1, they are exactly equal to 1. As shown in previous publications (*13*, *14*), the crucial ingredient of the model that leads to sub-Poisson noise is the switching of the promoter state upon production of a single nascent transcript, changing it from a state where no new RNAP can bind to a state where this is possible. This captures the observation that a promoter can only bind a new RNAP if there is no other bound RNAP within a short distance from it (*20*, *21*). Mathematically, the switching of the promoter state causes sub-Poisson noise in the transcript numbers because the coefficient of variation of the time between two successive nascent mRNA production events is less than 1, i.e. less than expected from an exponential distribution.

Note that if there was no state switching upon the production of a nascent transcript

$$U_S \to U_1 \to U_2 \to \cdots \to U_S \to U_S + N$$

then the Fano factors of nascent, nuclear and cytoplasmic mRNA would be greater than or equal to 1 (*22*, *23*). In models of this type, the interpretation is that $U_1$ to $U_{S-1}$ are transcriptional off states while $U_S$ is a transcriptionally active state. Nascent mRNA can be continuously produced from this state and there is no associated promoter clearing.

In steady-state conditions, the dynamics of nascent mRNA elongation and release do not influence the moments of the nuclear and cytoplasmic mRNA in our model. This is because we are assuming that release happens a certain fixed time $T$ after initiation of elongation, and hence the time between two subsequent nascent mRNA production events is the same as the time between two subsequent mature nuclear mRNA production events. In other words, the moments of nuclear and cytoplasmic mRNA in steady-state conditions remain the same if we instead modeled transcription as

$$U_1 \to U_2 \to \cdots \to U_S \to U_1 + M_N$$

where the last reaction has rate constant $k_B$.

In steady-state conditions, the Fano factor of cytoplasmic mRNA can either be larger or smaller than the Fano factor of nuclear mRNA. From the closed-form solutions it is found that in the limit $k_C$ goes to infinity, the Fano factor of cytoplasmic mRNA is smaller than that of nuclear mRNA. The exact value of $k_C$ when the two Fano factors are equal depends on the values of $k_A$, $k_B$, S and R.

The minimum Fano factor of nuclear mRNA is achieved when $k_B = k_A \gg k_C$, in other words when the rates for promoter remodeling and promoter freeing are in the same range and considerably faster than export. The minimum Fano factor of cytoplasmic mRNA is achieved in the limits $k_C \gg k_B = k_A \gg k_D$, in other words when the rates for promoter remodeling and promoter freeing are in the same range and considerably faster than degradation but slower than export. The minimum values possible for a model with S and R each between 1 and 4 are shown in Table B-I and Fig. 7G.

*Table B-I: Minimum Fano factor of nuclear and cytoplasmic mRNA*

Minimum Fano factors for the general model with S and R rate limiting steps in initiation and decay, respectively. The first number shows the nuclear Fano factor, the second number shows the cytoplasmic Fano factor.

| | $\boldsymbol{R = 1}$ | $\boldsymbol{R = 2}$ | $\boldsymbol{R = 3}$ | $\boldsymbol{R = 4}$ |
|---|---|---|---|---|
| $\mathbf{S = 1}$ | (1.00,1.00) | (1.00,1.00) | (1.00,1.00) | (1.00,1.00) |
| $\mathbf{S = 2}$ | (0.750,0.750) | (0.750,0.688) | (0.750,0.656) | (0.750,0.637) |
| $\mathbf{S = 3}$ | (0.667,0.667) | (0.667,0.583) | (0.667,0.542) | (0.667,0.516) |
| $\mathbf{S = 4}$ | (0.625,0.625) | (0.625,0.531) | (0.625,0.484) | (0.625,0.455) |

##### B.3 Bayesian model selection and parameter inference

While it is immediately clear from Table B-I that models with one rate limiting step for transcription will not be able to explain the sub-Poisson nature of the data, it is an open question which of the models with two or more rate-limiting steps in transcription and one or more rate-limiting steps in degradation best fits the data. To answer this question, we used a simple ABC (Approximate Bayesian Computation) rejection sampler (*24*) to perform Bayesian model selection (*25*).

For each gene and for a given model with certain values of S and R, the ABC algorithm randomly samples parameters $k_A$, $k_B$, $k_C$ from a prior distribution. Since we do not have prior knowledge about the 3 parameters, we simply choose the distribution to be uniform over a certain range. Stochastic simulations of the model with the selected parameters are run using Gillespie's exact algorithm (*26*) until steady-state is achieved. Then a distance function between the moments of the data and the model is evaluated. Note that the number of samples of Gillespie's algorithm equals the number of single-cell measurements for a given gene. This constraint makes sure that the simulation-generated moments have the same variability (due to finite number of samples) as the moments calculated from the data. Those parameters whose distance function is less than a certain threshold are accepted. The mean and standard deviation of the marginal distribution of these accepted parameters (the posterior) provides an estimate of the values of the parameters and their uncertainty. We also calculate the acceptance rate, which is the number of parameter sets that are accepted divided by the total number of parameter sets that were randomly generated by the ABC algorithm. We then select which model best selects the data for a given gene by comparing the acceptance rate of any two models. The Bayes factor of two models A and B is defined as the ratio of the acceptance rate of the two models (*25*). If the Bayes factor is larger than 10, we conclude that there is strong evidence in favor of model A; if the Bayes factor is less than 1/10, we conclude that there is strong evidence in favor of model B (*27*). Otherwise, if the Bayes factor is between 1/10 and 10, we will conclude that there is no strong evidence for model A or model B (there might be weak evidence).

The heart of the ABC algorithm involves accepting only those parameters which satisfy a user-defined criterion. In our case the criterion is that the following distance function is smaller than a preset threshold value:

$$d = \left(\frac{2\mu_C - \mu_C^e}{\mu_C^e}\right)^2 + \left(\frac{FF_C - FF_C^e}{FF_C^e}\right)^2 + \left(\frac{FF_N - FF_N^e}{FF_N^e}\right)^2$$

where $\mu_C$ and $FF_C$ are the simulation predictions for the mean and Fano factor (variance divided by the mean) of the cytoplasmic mRNA, $M_C$; and $FF_N$ is the simulation prediction for the Fano factor of the nuclear mRNA, $M_N$. The same variables but with an *e* superscript denote the experimental estimates of the mean and the Fano factors for non-transcription site (non-TS) labelled data from mononucleated cells. Note that in this section whenever we refer to the experimental mean of transcript numbers or the Fano factor, we always mean the volume-corrected values (see Methods). This is necessary since the model's predictions (with which the experimental values will be compared) are independent of the cell volume. Note also that the simulation predictions are for a single gene copy. Mononucleated *S. pombe* cells are almost exclusively in G2 phase of the cell cycle and hence have two (independent) gene copies (*28*, *29*).

Therefore, we multiply the model's estimate of the mean by 2 in the expression for the distance function. The Fano factor is unchanged because both the mean and variance are doubled.

Note that the distance function does not have information about the mean of nuclear mRNA but only about its Fano factor. This choice is motivated by a direct comparison of fitting to the data with or without the transcription site label for the two genes, *mad1* and *rpb1*. When the transcription site is not labelled (non-TS labelled), it is uncertain which fraction of the nuclear RNA is nascent RNA and which fraction is mature nuclear RNA. Not making this distinction leads to a considerably different estimate for the mean of nuclear RNA, $\mu_N$, from non-TS labelled data compared to that estimated from TS-labelled data (Table B-II). The relative error is around 200 %. In contrast, estimating the nuclear Fano factor, $FF_N$, is more robust (relative error less than 20 %). Furthermore, the estimates for both cytoplasmic mean and cytoplasmic Fano factor are similar, regardless of which data is used. We therefore excluded the mean of nuclear RNA, $\mu_N$, from the distance function.

*Table B-II: Comparison of the means (μ) and Fano factors of nuclear (N) and cytoplasmic (C) mRNA*

Parameters were estimated using both non-TS labelled and TS-labelled data for both genes. The largest discrepancies are observed for $\mu_N$ (red).

| | $\boldsymbol{\mu_N}$ | | $\boldsymbol{\mu_C}$ | | $\boldsymbol{FF_N}$ | | $\boldsymbol{FF_C}$ | |
|---|---|---|---|---|---|---|---|---|
| | non-TS labelled | TS-labelled | non-TS labelled | TS-labelled | non-TS labelled | TS-labelled | non-TS labelled | TS-labelled |
| *mad1* | 0.318 | 0.120 | 1.74 | 1.74 | 0.929 | 0.917 | 0.620 | 0.540 |
| *rpb1* | 3.56 | 1.15 | 17.9 | 15.7 | 0.826 | 0.939 | 0.647 | 0.560 |

The only remaining information to be specified for the ABC rejection sampler is the choice of the ranges over which the parameters are randomly sampled and the threshold values ($d_c$) of the distance function for each gene (Table B-III). We have not included *cdc13* in this analysis because its data is consistent with a Poisson distribution (Section A.4). Theoretically it is known that the distribution of accepted parameters from an ABC rejection sampler will be a good approximation to the true posterior distribution provided the threshold value is sufficiently small (*25*). A threshold value is hence chosen small enough so that repeating with a smaller value does not significantly change the posterior; this value will necessarily be different from one gene to another, particularly given the variable sample sizes. The range of the uniform prior distribution was initially chosen to be (0,250) for $k_A$, $k_B$, $k_C$. We noticed that in many cases initial runs of the ABC rejection sampler identified parameters satisfying the user-defined criterion only in small regions of this range. Hence to obtain a large number of accepted parameters within a computationally reasonable time (which is needed to obtain a well-defined posterior distribution) it was necessary to limit the range of the prior distributions to a subset of numbers within (0,250) which the initial exploration identified as containing the vast majority of the accepted parameters. We implemented the constraint that, for a given parameter set, the 12 models should be indistinguishable with respect to the mean nuclear and cytoplasmic mRNA. This is achieved by scaling $k_A$ by a factor proportional to $S-1$ (following from the equations for the first-order moments above).

*Table B-III: Priors for parameters $k_A$, $k_B$, $k_C$, and thresholds for the distance function ($d_c$)*
Note that $k_D = R$.

| **Gene** | $\boldsymbol{k_A}$ | $\boldsymbol{k_B}$ | $\boldsymbol{k_C}$ | $\boldsymbol{d_c}$ |
|---|---|---|---|---|
| $\mathrm{mad1}$ | $\frac{S-1}{2}(0.5,10.5)$ | $(0.5,10.5)$ | $(0.5,100.5)$ | 0.003 |
| $\mathrm{mad1-GFP}$ | $\frac{S-1}{2}(0.5,10.5)$ | $(0.5,40.5)$ | $(20,220)$ | 0.0001 |
| $\mathrm{mad2-GFP}$ | $\frac{S-1}{2}(0.5,35.5)$ | $(0.5,10.5)$ | $(0.5,20.5)$ | 0.005 |
| $\mathrm{mad3-GFP}$ | $\frac{S-1}{2}(0.5,20.5)$ | $(0.5,10.5)$ | $(20,220)$ | 0.00025 |
| $\mathrm{bub1-GFP}$ | $\frac{S-1}{2}(0.5,20.5)$ | $(0.5,75.5)$ | $(20,220)$ | 0.0001 |
| $\mathrm{rpb1}$ | $\frac{S-1}{2}(25,75)$ | $(0.5,30.5)$ | $(0.5,50.5)$ | 0.00075 |
| $\mathrm{sep1-GFP}$ | $\frac{S-1}{2}(0.5,100.5)$ | $(0.5,5.5)$ | $(0.1,30.1)$ | 0.00075 |

The Bayes factors were computed for all genes and for all 12 models with $2 \leq S \leq 4$ and $1 \leq R \leq 4$. The results are shown in Tables B-IV-X. For a given gene, the models highlighted in yellow are those which can explain the data, i.e. no other model was found to have an acceptance rate at least 10-times larger; or, in other words, all the 12 Bayes factors for this model are greater than or equal to 1/10. The model selection results are summarized in Table B-XI. We find strong evidence that *mad1*, *mad1*-GFP, and *bub1*-GFP are best described by models with 3 or 4 rate-limiting steps in transcription and 2 – 4 rate-limiting steps in degradation. In contrast, for the rest of the genes (*mad2*-GFP, *mad3*-GFP, *rpb1*, *sep1*-GFP), most of the 12 models fit the data equally well. We also note that the models with S = 3 and R = 3 or 4, as well as S = 4 and R = 2, are the *only* models that can explain the data from all genes.

*Table B-IV: Bayes factors for mad1*

The value in the $i^{th}$ row and $j^{th}$ column is the Bayes factor of models i and j. Bayes factors $\geq 10$ are shown in blue (model i explains the data better than model j); Bayes factors $\leq 1/10$ are shown in red (model j explains the data better than model i); Bayes factors in the range (1/10,10) are shown in black, which is interpreted as the two models being indistinguishable. Those models which explain the data best, i.e. those lacking Bayes factors $\leq 1/10$, are shaded in yellow.

| mad1 | S2_R1 | S2_R2 | S2_R3 | S2_R4 | S3_R1 | S3_R2 | S3_R3 | S3_R4 | S4_R1 | S4_R2 | S4_R3 | S4_R4 |
|---|---|---|---|---|---|---|---|---|---|---|---|---|
| **S2_R1** | 1.0E+00 | 1.0E+00 | 1.0E+00 | 2.3E+00 | 1.3E+00 | 5.8E+02 | 4.3E+03 | 8.1E+03 | 6.5E+01 | 9.0E+03 | 9.5E+03 | 4.8E+03 |
| **S2_R2** | 1.0E+00 | 1.0E+00 | 1.0E+00 | 2.3E+00 | 1.3E+00 | 5.8E+02 | 4.3E+03 | 8.1E+03 | 6.5E+01 | 9.0E+03 | 9.5E+03 | 4.8E+03 |
| **S2_R3** | 1.0E+00 | 1.0E+00 | 1.0E+00 | 2.3E+00 | 5.7E-01 | 5.8E+02 | 4.3E+03 | 8.1E+03 | 6.5E+01 | 9.0E+03 | 9.5E+03 | 4.8E+03 |
| **S2_R4** | 4.4E-01 | 4.4E-01 | 4.4E-01 | 1.0E+00 | 5.7E-01 | 2.5E+02 | 1.9E+03 | 3.5E+03 | 2.8E+01 | 3.9E+03 | 4.2E+03 | 2.1E+03 |
| **S3_R1** | 7.8E-01 | 7.8E-01 | 7.8E-01 | 1.8E+00 | 1.0E+00 | 4.5E+02 | 3.3E+03 | 6.3E+03 | 5.0E+01 | 7.0E+03 | 7.4E+03 | 3.8E+03 |
| **S3_R2** | 1.7E-03 | 1.7E-03 | 1.7E-03 | 4.0E-03 | 2.2E-03 | 1.0E+00 | 7.4E+00 | 1.4E+01 | 1.1E-01 | 1.6E+01 | 1.7E+01 | 8.4E+00 |
| **S3_R3** | 2.3E-04 | 2.3E-04 | 2.3E-04 | 5.3E-04 | 3.0E-04 | 1.3E-01 | 1.0E+00 | 1.9E+00 | 1.5E-02 | 2.1E+00 | 2.2E+00 | 1.1E+00 |
| **S3_R4** | 1.2E-04 | 1.2E-04 | 1.2E-04 | 2.8E-04 | 1.6E-04 | 7.1E-02 | 5.3E-01 | 1.0E+00 | 8.0E-03 | 1.1E+00 | 1.2E+00 | 6.0E-01 |
| **S4_R1** | 1.5E-02 | 1.5E-02 | 1.5E-02 | 3.5E-02 | 2.0E-02 | 8.9E+00 | 6.6E+01 | 1.2E+02 | 1.0E+00 | 1.4E+02 | 1.5E+02 | 7.5E+01 |
| **S4_R2** | 1.1E-04 | 1.1E-04 | 1.1E-04 | 2.5E-04 | 1.4E-04 | 6.4E-02 | 4.8E-01 | 9.0E-01 | 7.2E-03 | 1.0E+00 | 1.1E+00 | 5.4E-01 |
| **S4_R3** | 1.0E-04 | 1.0E-04 | 1.0E-04 | 2.4E-04 | 1.4E-04 | 6.0E-02 | 4.5E-01 | 8.5E-01 | 6.8E-03 | 9.4E-01 | 1.0E+00 | 5.1E-01 |
| **S4_R4** | 2.1E-04 | 2.1E-04 | 2.1E-04 | 4.7E-04 | 2.7E-04 | 1.2E-01 | 8.9E-01 | 1.7E+00 | 1.3E-02 | 1.9E+00 | 2.0E+00 | 1.0E+00 |

*Table B-V: Bayes factors for mad1-GFP*

| mad1-GFP | S2_R1 | S2_R2 | S2_R3 | S2_R4 | S3_R1 | S3_R2 | S3_R3 | S3_R4 | S4_R1 | S4_R2 | S4_R3 | S4_R4 |
|---|---|---|---|---|---|---|---|---|---|---|---|---|
| **S2_R1** | 1.0E+00 | 1.0E+00 | 1.9E+01 | 4.7E+01 | 1.3E+01 | 6.4E+02 | 9.9E+02 | 8.8E+02 | 4.0E+02 | 2.4E+02 | 7.4E+01 | 6.6E+01 |
| **S2_R2** | 1.0E+00 | 1.0E+00 | 1.9E+01 | 4.7E+01 | 1.3E+01 | 6.4E+02 | 9.9E+02 | 8.8E+02 | 4.0E+02 | 2.4E+02 | 7.4E+01 | 6.6E+01 |
| **S2_R3** | 5.1E-02 | 5.1E-02 | 1.0E+00 | 2.4E+00 | 6.6E-01 | 3.3E+01 | 5.1E+01 | 4.5E+01 | 2.0E+01 | 1.2E+01 | 3.8E+00 | 3.4E+00 |
| **S2_R4** | 2.1E-02 | 2.1E-02 | 4.2E-01 | 1.0E+00 | 2.7E-01 | 1.4E+01 | 2.1E+01 | 1.9E+01 | 8.4E+00 | 5.0E+00 | 1.6E+00 | 1.4E+00 |
| **S3_R1** | 7.8E-02 | 7.8E-02 | 1.5E+00 | 3.6E+00 | 1.0E+00 | 5.0E+01 | 7.7E+01 | 6.8E+01 | 3.1E+01 | 1.8E+01 | 5.8E+00 | 5.1E+00 |
| **S3_R2** | 1.6E-03 | 1.6E-03 | 3.0E-02 | 7.3E-02 | 2.0E-02 | 1.0E+00 | 1.5E+00 | 1.4E+00 | 6.1E-01 | 3.7E-01 | 1.2E-01 | 1.0E-01 |
| **S3_R3** | 1.0E-03 | 1.0E-03 | 2.0E-02 | 4.7E-02 | 1.3E-02 | 6.5E-01 | 1.0E+00 | 8.9E-01 | 4.0E-01 | 2.4E-01 | 7.5E-02 | 6.7E-02 |
| **S3_R4** | 1.1E-03 | 1.1E-03 | 2.2E-02 | 5.3E-02 | 1.5E-02 | 7.3E-01 | 1.1E+00 | 1.0E+00 | 4.5E-01 | 2.7E-01 | 8.4E-02 | 7.5E-02 |
| **S4_R1** | 2.5E-03 | 2.5E-03 | 4.9E-02 | 1.2E-01 | 3.3E-02 | 1.6E+00 | 2.5E+00 | 2.2E+00 | 1.0E+00 | 5.9E-01 | 1.9E-01 | 1.7E-01 |
| **S4_R2** | 4.3E-03 | 4.3E-03 | 8.3E-02 | 2.0E-01 | 5.5E-02 | 2.7E+00 | 4.2E+00 | 3.7E+00 | 1.7E+00 | 1.0E+00 | 3.2E-01 | 2.8E-01 |
| **S4_R3** | 1.3E-02 | 1.3E-02 | 2.6E-01 | 6.3E-01 | 1.7E-01 | 8.7E+00 | 1.3E+01 | 1.2E+01 | 5.3E+00 | 3.2E+00 | 1.0E+00 | 8.9E-01 |
| **S4_R4** | 1.5E-02 | 1.5E-02 | 2.9E-01 | 7.1E-01 | 1.9E-01 | 9.7E+00 | 1.5E+01 | 1.3E+01 | 6.0E+00 | 3.6E+00 | 1.1E+00 | 1.0E+00 |

*Table B-VI: Bayes factors for mad2-GFP*

| mad2-GFP | S2_R1 | S2_R2 | S2_R3 | S2_R4 | S3_R1 | S3_R2 | S3_R3 | S3_R4 | S4_R1 | S4_R2 | S4_R3 | S4_R4 |
|---|---|---|---|---|---|---|---|---|---|---|---|---|
| **S2_R1** | 1.0E+00 | 8.9E-01 | 7.4E-01 | 6.6E-01 | 6.0E-01 | 4.2E-01 | 3.5E-01 | 3.4E-01 | 5.3E-01 | 4.3E-01 | 3.8E-01 | 3.5E-01 |
| **S2_R2** | 1.1E+00 | 1.0E+00 | 8.4E-01 | 7.4E-01 | 6.8E-01 | 4.7E-01 | 3.9E-01 | 3.8E-01 | 6.0E-01 | 4.8E-01 | 4.2E-01 | 3.9E-01 |
| **S2_R3** | 1.3E+00 | 1.2E+00 | 1.0E+00 | 8.8E-01 | 8.1E-01 | 5.7E-01 | 4.7E-01 | 4.5E-01 | 7.1E-01 | 5.8E-01 | 5.1E-01 | 4.6E-01 |
| **S2_R4** | 1.5E+00 | 1.4E+00 | 1.1E+00 | 1.0E+00 | 9.2E-01 | 6.4E-01 | 5.3E-01 | 5.2E-01 | 8.1E-01 | 6.6E-01 | 5.8E-01 | 5.3E-01 |
| **S3_R1** | 1.7E+00 | 1.5E+00 | 1.2E+00 | 1.1E+00 | 1.0E+00 | 7.0E-01 | 5.8E-01 | 5.6E-01 | 8.8E-01 | 7.2E-01 | 6.3E-01 | 5.8E-01 |
| **S3_R2** | 2.4E+00 | 2.1E+00 | 1.8E+00 | 1.6E+00 | 1.4E+00 | 1.0E+00 | 8.3E-01 | 8.0E-01 | 1.3E+00 | 1.0E+00 | 9.0E-01 | 8.2E-01 |
| **S3_R3** | 2.9E+00 | 2.5E+00 | 2.1E+00 | 1.9E+00 | 1.7E+00 | 1.2E+00 | 1.0E+00 | 9.7E-01 | 1.5E+00 | 1.2E+00 | 1.1E+00 | 9.9E-01 |
| **S3_R4** | 3.0E+00 | 2.6E+00 | 2.2E+00 | 1.9E+00 | 1.8E+00 | 1.2E+00 | 1.0E+00 | 1.0E+00 | 1.6E+00 | 1.3E+00 | 1.1E+00 | 1.0E+00 |
| **S4_R1** | 1.9E+00 | 1.7E+00 | 1.4E+00 | 1.2E+00 | 1.1E+00 | 7.9E-01 | 6.6E-01 | 6.4E-01 | 1.0E+00 | 8.1E-01 | 7.1E-01 | 6.5E-01 |
| **S4_R2** | 2.3E+00 | 2.1E+00 | 1.7E+00 | 1.5E+00 | 1.4E+00 | 9.8E-01 | 8.1E-01 | 7.9E-01 | 1.2E+00 | 1.0E+00 | 8.8E-01 | 8.0E-01 |
| **S4_R3** | 2.7E+00 | 2.4E+00 | 2.0E+00 | 1.7E+00 | 1.6E+00 | 1.1E+00 | 9.2E-01 | 9.0E-01 | 1.4E+00 | 1.1E+00 | 1.0E+00 | 9.2E-01 |
| **S4_R4** | 2.9E+00 | 2.6E+00 | 2.2E+00 | 1.9E+00 | 1.7E+00 | 1.2E+00 | 1.0E+00 | 9.8E-01 | 1.5E+00 | 1.2E+00 | 1.1E+00 | 1.0E+00 |

*Table B-VII: Bayes factors for mad3-GFP*

| mad3-GFP | S2_R1 | S2_R2 | S2_R3 | S2_R4 | S3_R1 | S3_R2 | S3_R3 | S3_R4 | S4_R1 | S4_R2 | S4_R3 | S4_R4 |
|---|---|---|---|---|---|---|---|---|---|---|---|---|
| **S2_R1** | 1.0E+00 | 6.1E+01 | 6.5E+01 | 5.7E+01 | 8.3E+01 | 2.0E+01 | 2.1E+01 | 2.1E+01 | 2.7E+01 | 2.0E+01 | 2.0E+01 | 1.9E+01 |
| **S2_R2** | 1.6E-02 | 1.0E+00 | 1.1E+00 | 9.4E-01 | 1.4E+00 | 3.3E-01 | 3.4E-01 | 3.5E-01 | 4.4E-01 | 3.3E-01 | 3.3E-01 | 3.2E-01 |
| **S2_R3** | 1.5E-02 | 9.3E-01 | 1.0E+00 | 8.8E-01 | 1.3E+00 | 3.1E-01 | 3.2E-01 | 3.2E-01 | 4.1E-01 | 3.0E-01 | 3.1E-01 | 2.9E-01 |
| **S2_R4** | 1.7E-02 | 1.1E+00 | 1.1E+00 | 1.0E+00 | 1.5E+00 | 3.5E-01 | 3.6E-01 | 3.7E-01 | 4.7E-01 | 3.5E-01 | 3.5E-01 | 3.4E-01 |
| **S3_R1** | 1.2E-02 | 7.3E-01 | 7.9E-01 | 6.9E-01 | 1.0E+00 | 2.4E-01 | 2.5E-01 | 2.5E-01 | 3.2E-01 | 2.4E-01 | 2.4E-01 | 2.3E-01 |
| **S3_R2** | 5.0E-02 | 3.0E+00 | 3.3E+00 | 2.9E+00 | 4.2E+00 | 1.0E+00 | 1.0E+00 | 1.1E+00 | 1.3E+00 | 1.0E+00 | 1.0E+00 | 9.6E-01 |
| **S3_R3** | 4.8E-02 | 2.9E+00 | 3.1E+00 | 2.8E+00 | 4.0E+00 | 9.6E-01 | 1.0E+00 | 1.0E+00 | 1.3E+00 | 9.6E-01 | 9.7E-01 | 9.3E-01 |
| **S3_R4** | 4.7E-02 | 2.9E+00 | 3.1E+00 | 2.7E+00 | 3.9E+00 | 9.5E-01 | 9.9E-01 | 1.0E+00 | 1.3E+00 | 9.5E-01 | 9.6E-01 | 9.1E-01 |
| **S4_R1** | 3.7E-02 | 2.3E+00 | 2.4E+00 | 2.1E+00 | 3.1E+00 | 7.4E-01 | 7.7E-01 | 7.8E-01 | 1.0E+00 | 7.4E-01 | 7.5E-01 | 7.2E-01 |
| **S4_R2** | 5.0E-02 | 3.1E+00 | 3.3E+00 | 2.9E+00 | 4.2E+00 | 1.0E+00 | 1.0E+00 | 1.1E+00 | 1.4E+00 | 1.0E+00 | 1.0E+00 | 9.7E-01 |
| **S4_R3** | 4.9E-02 | 3.0E+00 | 3.2E+00 | 2.8E+00 | 4.1E+00 | 9.9E-01 | 1.0E+00 | 1.0E+00 | 1.3E+00 | 9.8E-01 | 1.0E+00 | 9.5E-01 |
| **S4_R4** | 5.2E-02 | 3.2E+00 | 3.4E+00 | 3.0E+00 | 4.3E+00 | 1.0E+00 | 1.1E+00 | 1.1E+00 | 1.4E+00 | 1.0E+00 | 1.1E+00 | 1.0E+00 |

*Table B-VIII: Bayes factors for bub1-GFP*

| bub1-GFP | S2_R1 | S2_R2 | S2_R3 | S2_R4 | S3_R1 | S3_R2 | S3_R3 | S3_R4 | S4_R1 | S4_R2 | S4_R3 | S4_R4 |
|---|---|---|---|---|---|---|---|---|---|---|---|---|
| **S2_R1** | 1.0E+00 | 2.8E+00 | 1.3E+01 | 5.4E+01 | 1.5E+01 | 8.8E+02 | 1.5E+03 | 1.5E+03 | 3.9E+02 | 1.0E+03 | 2.1E+02 | 1.6E+02 |
| **S2_R2** | 3.6E-01 | 1.0E+00 | 4.7E+00 | 1.9E+01 | 5.4E+00 | 3.1E+02 | 5.2E+02 | 5.2E+02 | 1.4E+02 | 3.6E+02 | 7.3E+01 | 5.6E+01 |
| **S2_R3** | 7.7E-02 | 2.1E-01 | 1.0E+00 | 4.1E+00 | 1.2E+00 | 6.8E+01 | 1.1E+02 | 1.1E+02 | 3.0E+01 | 7.8E+01 | 1.6E+01 | 1.2E+01 |
| **S2_R4** | 1.9E-02 | 5.2E-02 | 2.4E-01 | 1.0E+00 | 2.8E-01 | 1.6E+01 | 2.7E+01 | 2.7E+01 | 7.2E+00 | 1.9E+01 | 3.8E+00 | 3.0E+00 |
| **S3_R1** | 6.6E-02 | 1.9E-01 | 8.6E-01 | 3.5E+00 | 1.0E+00 | 5.8E+01 | 9.6E+01 | 9.7E+01 | 2.6E+01 | 6.7E+01 | 1.4E+01 | 1.0E+01 |
| **S3_R2** | 1.1E-03 | 3.2E-03 | 1.5E-02 | 6.1E-02 | 1.7E-02 | 1.0E+00 | 1.7E+00 | 1.7E+00 | 4.4E-01 | 1.1E+00 | 2.3E-01 | 1.8E-01 |
| **S3_R3** | 6.9E-04 | 1.9E-03 | 9.0E-03 | 3.7E-02 | 1.0E-02 | 6.1E-01 | 1.0E+00 | 1.0E+00 | 2.7E-01 | 7.0E-01 | 1.4E-01 | 1.1E-01 |
| **S3_R4** | 6.8E-04 | 1.9E-03 | 6.5E-01 | 3.7E-02 | 1.0E-02 | 6.0E-01 | 1.0E+00 | 1.0E+00 | 2.6E-01 | 6.9E-01 | 1.4E-01 | 1.1E-01 |
| **S4_R1** | 2.6E-03 | 7.2E-03 | 3.4E-02 | 1.4E-01 | 3.9E-02 | 2.3E+00 | 3.8E+00 | 3.8E+00 | 1.0E+00 | 2.6E+00 | 5.3E-01 | 4.1E-01 |
| **S4_R2** | 9.9E-04 | 2.8E-03 | 1.3E-02 | 5.3E-02 | 1.5E-02 | 8.7E-01 | 1.4E+00 | 1.4E+00 | 3.8E-01 | 1.0E+00 | 2.0E-01 | 1.6E-01 |
| **S4_R3** | 4.9E-03 | 1.4E-02 | 6.4E-02 | 2.6E-01 | 7.4E-02 | 4.3E+00 | 7.1E+00 | 7.1E+00 | 1.9E+00 | 5.0E+00 | 1.0E+00 | 7.7E-01 |
| **S4_R4** | 6.3E-03 | 1.8E-02 | 8.2E-02 | 3.4E-01 | 9.6E-02 | 5.6E+00 | 7.0E+00 | 9.2E+00 | 2.4E+00 | 6.4E+00 | 1.3E+00 | 1.0E+00 |

*Table B-IX: Bayes factors for rpb1*

| rpb1 | S2_R1 | S2_R2 | S2_R3 | S2_R4 | S3_R1 | S3_R2 | S3_R3 | S3_R4 | S4_R1 | S4_R2 | S4_R3 | S4_R4 |
|---|---|---|---|---|---|---|---|---|---|---|---|---|
| **S2_R1** | 1.0E+00 | 1.5E+02 | 9.5E+02 | 1.9E+03 | 8.4E+02 | 1.6E+03 | 1.1E+03 | 9.3E+02 | 1.9E+03 | 1.1E+03 | 1.0E+03 | 9.8E+02 |
| **S2_R2** | 6.6E-03 | 1.0E+00 | 6.3E+00 | 1.2E+01 | 5.5E+00 | 1.0E+01 | 7.3E+00 | 6.2E+00 | 1.3E+01 | 7.5E+00 | 6.8E+00 | 6.5E+00 |
| **S2_R3** | 1.1E-03 | 1.6E-01 | 1.0E+00 | 2.0E+00 | 8.8E-01 | 1.7E+00 | 1.2E+00 | 9.8E-01 | 2.0E+00 | 1.2E+00 | 1.1E+00 | 1.0E+00 |
| **S2_R4** | 5.3E-04 | 8.0E-02 | 5.0E-01 | 1.0E+00 | 4.5E-01 | 8.4E-01 | 5.9E-01 | 5.0E-01 | 1.0E+00 | 6.0E-01 | 5.4E-01 | 5.2E-01 |
| **S3_R1** | 1.2E-03 | 1.8E-01 | 1.1E+00 | 2.2E+00 | 1.0E+00 | 1.9E+00 | 1.3E+00 | 1.1E+00 | 2.3E+00 | 1.3E+00 | 1.2E+00 | 1.2E+00 |
| **S3_R2** | 6.3E-04 | 9.6E-02 | 6.0E-01 | 1.2E+00 | 5.3E-01 | 1.0E+00 | 7.1E-01 | 5.9E-01 | 1.2E+00 | 7.2E-01 | 6.5E-01 | 6.2E-01 |
| **S3_R3** | 9.0E-04 | 1.4E-01 | 8.5E-01 | 1.7E+00 | 7.5E-01 | 1.4E+00 | 1.0E+00 | 8.4E-01 | 1.7E+00 | 1.0E+00 | 9.2E-01 | 8.8E-01 |
| **S3_R4** | 1.1E-03 | 1.6E-01 | 1.0E+00 | 2.0E+00 | 9.0E-01 | 1.7E+00 | 1.2E+00 | 1.0E+00 | 2.1E+00 | 1.2E+00 | 1.1E+00 | 1.1E+00 |
| **S4_R1** | 5.2E-04 | 7.9E-02 | 4.9E-01 | 9.8E-01 | 4.4E-01 | 8.2E-01 | 5.8E-01 | 4.8E-01 | 1.0E+00 | 5.9E-01 | 5.3E-01 | 5.1E-01 |
| **S4_R2** | 8.8E-04 | 1.3E-01 | 8.4E-01 | 1.7E+00 | 7.4E-01 | 1.4E+00 | 9.8E-01 | 8.2E-01 | 1.7E+00 | 1.0E+00 | 9.0E-01 | 8.7E-01 |
| **S4_R3** | 9.8E-04 | 1.5E-01 | 9.3E-01 | 1.8E+00 | 8.2E-01 | 1.5E+00 | 1.1E+00 | 9.1E-01 | 1.9E+00 | 1.1E+00 | 1.0E+00 | 9.6E-01 |
| **S4_R4** | 1.0E-03 | 1.5E-01 | 9.7E-01 | 1.9E+00 | 8.6E-01 | 1.6E+00 | 1.1E+00 | 9.5E-01 | 2.0E+00 | 1.2E+00 | 1.0E+00 | 1.0E+00 |

*Table B-X: Bayes factors for sep1-GFP*

| sep1-GFP | S2_R1 | S2_R2 | S2_R3 | S2_R4 | S3_R1 | S3_R2 | S3_R3 | S3_R4 | S4_R1 | S4_R2 | S4_R3 | S4_R4 |
|---|---|---|---|---|---|---|---|---|---|---|---|---|
| **S2_R1** | 1.0E+00 | 8.0E-01 | 7.4E-01 | 7.1E-01 | 9.9E-01 | 8.4E-01 | 7.7E-01 | 7.3E-01 | 1.0E+00 | 8.6E-01 | 8.1E-01 | 7.5E-01 |
| **S2_R2** | 1.2E+00 | 1.0E+00 | 9.2E-01 | 8.8E-01 | 1.2E+00 | 1.1E+00 | 9.6E-01 | 9.1E-01 | 1.3E+00 | 1.1E+00 | 1.0E+00 | 9.4E-01 |
| **S2_R3** | 1.4E+00 | 1.1E+00 | 1.0E+00 | 9.6E-01 | 1.3E+00 | 1.1E+00 | 1.0E+00 | 9.9E-01 | 1.4E+00 | 1.2E+00 | 1.1E+00 | 1.0E+00 |
| **S2_R4** | 1.4E+00 | 1.1E+00 | 1.0E+00 | 1.0E+00 | 1.4E+00 | 1.2E+00 | 1.1E+00 | 1.0E+00 | 1.4E+00 | 1.2E+00 | 1.1E+00 | 1.1E+00 |
| **S3_R1** | 1.0E+00 | 8.1E-01 | 7.4E-01 | 7.1E-01 | 1.0E+00 | 8.5E-01 | 7.8E-01 | 7.3E-01 | 1.0E+00 | 8.7E-01 | 8.2E-01 | 7.6E-01 |
| **S3_R2** | 1.2E+00 | 9.5E-01 | 8.8E-01 | 8.4E-01 | 1.2E+00 | 1.0E+00 | 9.1E-01 | 8.6E-01 | 1.2E+00 | 1.0E+00 | 9.6E-01 | 9.0E-01 |
| **S3_R3** | 1.3E+00 | 1.0E+00 | 9.6E-01 | 9.2E-01 | 1.3E+00 | 1.1E+00 | 1.0E+00 | 9.5E-01 | 1.3E+00 | 1.1E+00 | 1.1E+00 | 9.8E-01 |
| **S3_R4** | 1.4E+00 | 1.1E+00 | 1.0E+00 | 9.7E-01 | 1.4E+00 | 1.2E+00 | 1.1E+00 | 1.0E+00 | 1.4E+00 | 1.2E+00 | 1.1E+00 | 1.0E+00 |
| **S4_R1** | 9.9E-01 | 7.9E-01 | 7.3E-01 | 7.0E-01 | 9.8E-01 | 8.3E-01 | 7.6E-01 | 7.2E-01 | 1.0E+00 | 8.5E-01 | 8.0E-01 | 7.5E-01 |
| **S4_R2** | 1.2E+00 | 9.3E-01 | 8.6E-01 | 8.2E-01 | 1.1E+00 | 9.8E-01 | 8.9E-01 | 8.4E-01 | 1.2E+00 | 1.0E+00 | 9.4E-01 | 8.7E-01 |
| **S4_R3** | 1.2E+00 | 9.9E-01 | 9.1E-01 | 8.7E-01 | 1.2E+00 | 1.0E+00 | 9.5E-01 | 9.0E-01 | 1.2E+00 | 1.1E+00 | 1.0E+00 | 9.3E-01 |
| **S4_R4** | 1.3E+00 | 1.1E+00 | 9.8E-01 | 9.4E-01 | 1.3E+00 | 1.1E+00 | 1.0E+00 | 9.6E-01 | 1.3E+00 | 1.1E+00 | 1.1E+00 | 1.0E+00 |

*Table B-XI: Summary of Bayesian model selection results*

The symbol X denotes a model which can explain the data for a particular gene, i.e. all the 12 Bayes factors for this model are greater than or equal to 1/10. Those models that explain data from all genes are shaded in yellow.

| | **S2_R1** | **S2_R2** | **S2_R3** | **S2_R4** | **S3_R1** | **S3_R2** | **S3_R3** | **S3_R4** | **S4_R1** | **S4_R2** | **S4_R3** | **S4_R4** |
|---|---|---|---|---|---|---|---|---|---|---|---|---|
| mad1 | | | | | | | X | X | | X | X | X |
| mad1 − GFP | | | | | | X | X | X | X | X | | |
| mad2 − GFP | X | X | X | X | X | X | X | X | X | X | X | X |
| mad3 − GFP | | X | X | X | X | X | X | X | X | X | X | X |
| bub1 − GFP | | | | | | X | X | X | X | X | X | X |
| rpb1 | | | X | X | X | X | X | X | X | X | X | X |
| sep1 − GFP | X | X | X | X | X | X | X | X | X | X | X | X |

Furthermore, from the posterior distributions we can estimate the value of the parameters and quantify their uncertainty. For this purpose, we used the model S = R = 3 since it is one of the few models which can explain the data from all genes. In Table B-XII we show the estimates of the three parameters and their uncertainty (mean and standard deviation of their posterior distributions), as well as two biologically meaningful functions of these parameters:

- $T_{init}/T_{degr}$, the ratio of the mean time between two successive transcription events and the mean time between two successive cytoplasmic degradation events, and
- $T_{expt}/T_{init}$, the ratio of the mean time for nuclear export and the mean time between two successive transcription events.

Computing percentage error as standard deviation divided by the mean, the percentage error is in the range 10–36 % for $k_A$, 4–69 % for $k_B$, 21–66 % for $k_C$, 1.4–3.3 % for $T_{init}/T_{degr}$, and 26–186 % for $T_{expt}/T_{init}$. Hence, the most clearly identifiable parameter is $T_{init}/T_{degr}$. The underlying reason is that this parameter is equal to the model's prediction of the inverse of the cytoplasmic mean which has been directly measured. The large errors in $T_{expt}/T_{init}$ are to be expected because this quantity is equal to the model's prediction of the nuclear mean which we do not use in our distance function since the estimated mean from experimental data without transcription site label is significantly different from that estimated from data with transcription site label. Nevertheless, the mean nuclear mRNA per gene copy predicted for *mad1* and *rpb1* from non-TS labelled data are in good agreement with those measured in the TS-labelled data (Table B-XII), which is remarkable given that the TS-labelled data was not used to estimate parameters.

Note that the posterior distributions are not uniform (as the priors), but peaked (Figure B-I), implying that the inference procedure was successful at finding optimal parameter values for each gene. The identified parameters varied across genes (Figure B-I,B-II).

*Table B-XII: Estimation of parameters from non-TS labelled data*

Estimation uses the S = R = 3 model with $k_D$=R. In brackets we show the estimated uncertainty; this is computed using the standard deviation of the accepted ABC sample estimates. $T_{expt}/T_{init}$ is equal to the model's prediction of the nuclear mean mature nuclear mRNA number. In the last column, for comparison, we show the mean mature nuclear mRNA number per gene copy from TS-labelled data (Fig. 7D). Note that the choice $k_D$=R implies that $k_A$, $k_B$, $k_C$ have units of inverse time where one time unit equals the mean time for one cytoplasmic mRNA molecule to decay.

| **Gene** | $\boldsymbol{k_A}$ | $\boldsymbol{k_B}$ | $\boldsymbol{k_C}$ | $\boldsymbol{T_{init}/T_{degr}}$ | $\boldsymbol{T_{expt}/T_{init}}$ | mean mature nuclear mRNA per gene copy |
|---|---|---|---|---|---|---|
| mad1 | 2.54 (0.504) | 3.64 (1.79) | 20.0 (9.08) | 1.15 (0.0366) | 0.0533 (0.0248) | 0.064 |
| mad1 − GFP | 3.49 (1.24) | 20.4 (9.92) | 111 (37.9) | 0.7 (0.0106) | 0.0145 (0.00559) | N/A |
| mad2 − GFP | 14.8 (3.46) | 1.84 (0.125) | 4.33 (2.33) | 0.689 (0.0230) | 0.453 (0.271) | N/A |
| mad3 − GFP | 13.3 (1.39) | 2.42 (0.0997) | 90.4 (31.5) | 0.567 (0.00804) | 0.0220 (0.00851) | N/A |
| bub1 − GFP | 5.62 (1.80) | 26.9 (18.5) | 45.5 (9.77) | 0.455 (0.00732) | 0.0510 (0.0133) | N/A |
| rpb1 | 49.2 (5.82) | 14.3 (1.07) | 27.0 (9.35) | 0.112 (0.00157) | 0.382 (0.164) | 0.604 |
| sep1 − GFP | 53.7 (19.1) | 3.08 (0.179) | 10.0 (6.61) | 0.368 (0.00983) | 0.636 (1.18) | N/A |

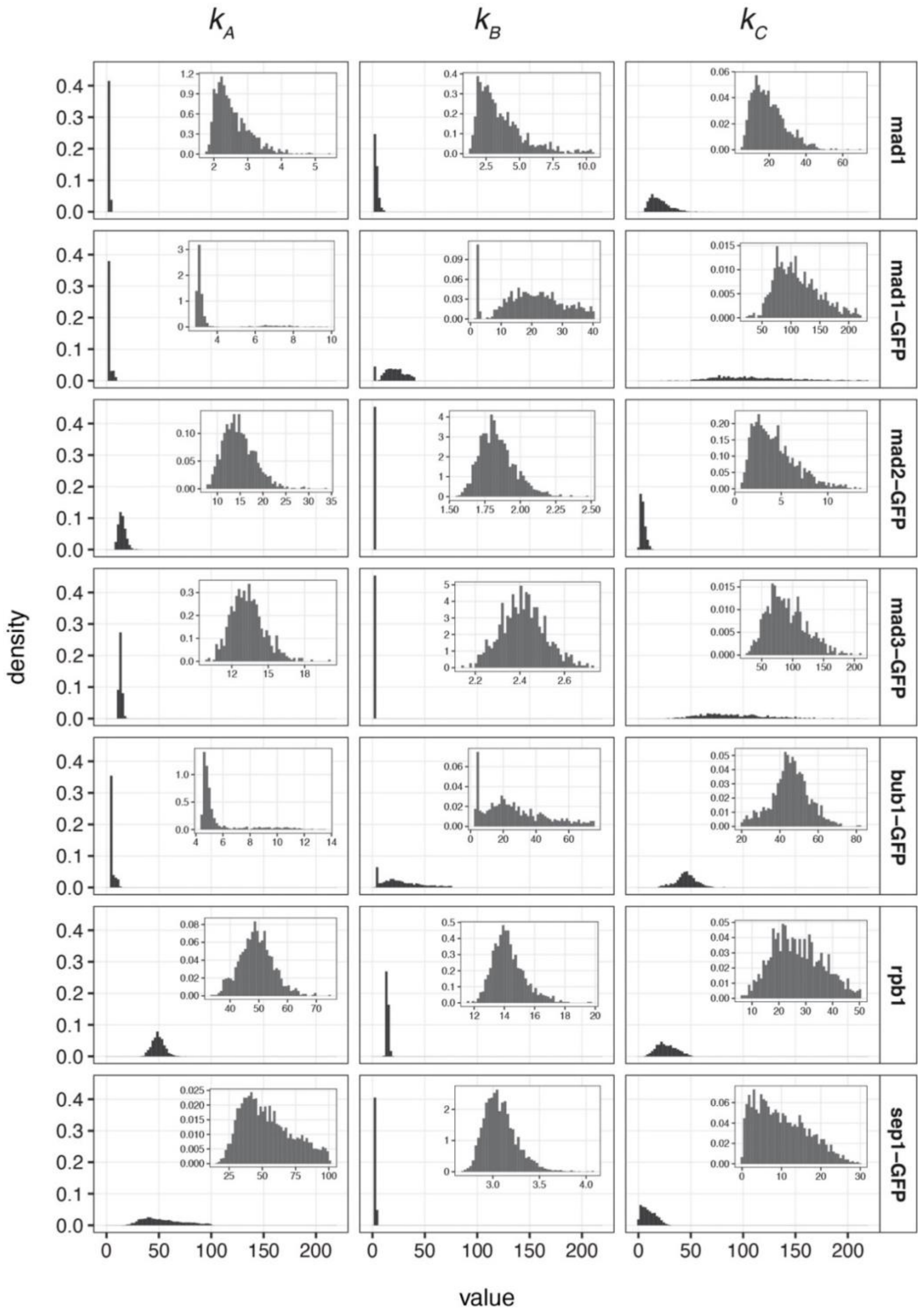


*Figure B-I: Posterior distributions of $k_A$, $k_B$, and $k_C$*

Posterior distributions were obtained using an ABC rejection sampler with the model with S = R = 3.

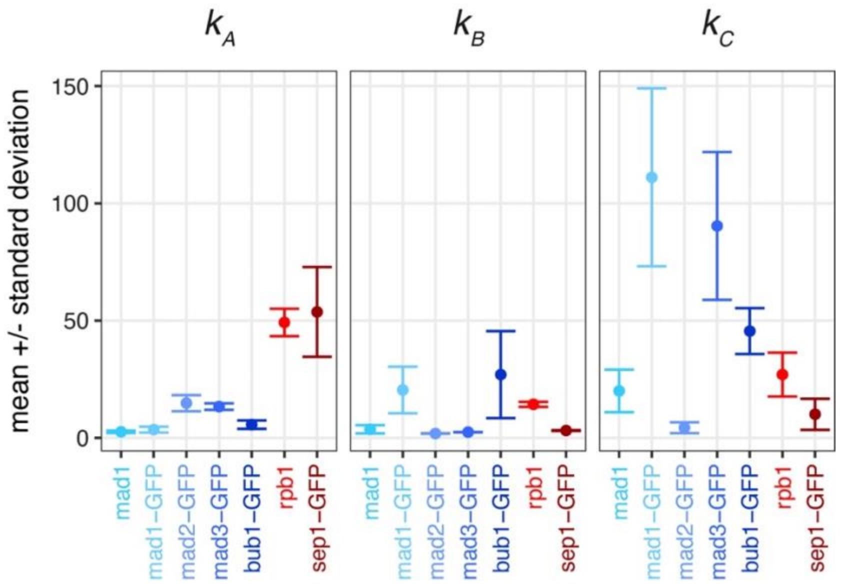


*Figure B-II: Overview on estimates of $k_A$, $k_B$, and $k_C$ from the posterior distributions*

Same data as in Figure B-I. Mean +/- standard deviation is shown.

*Table B-XIII: Nuclear export rate $k_C$ above which the Fano factor of cytoplasmic mRNA becomes smaller than the Fano factor of nuclear mRNA*

These are computed from the analytical expression of the model assuming $k_A = k_B = 10$.

| | $\boldsymbol{R = 1}$ | $\boldsymbol{R = 2}$ | $\boldsymbol{R = 3}$ | $\boldsymbol{R = 4}$ |
|---|---|---|---|---|
| $\mathbf{S = 2}$ | 4.58 | 2.60 | 2.14 | 1.94 |
| $\mathbf{S = 3}$ | 3.87 | 2.38 | 2.01 | 1.84 |
| $\mathbf{S = 4}$ | 3.39 | 2.21 | 1.89 | 1.75 |

#### B.4 Testing the ansatz of Poisson fluctuations

We considered the possibility that—while the measurements indicate sub-Poissonian mRNA distributions—they may instead result from a Poisson process and appear sub-Poissonian due to finite sample sizes. Specifically, if *n* Poisson-distributed random variables are generated (mimicking the mRNA number produced with a single rate-limiting step and measured in *n* cells), and the Fano factor (variance / mean) is calculated for this population of cells, then the Fano factor will vary around 1 and has a 50 % chance of being less than 1. In fact, it is known that the sampling distribution of the Fano factor is approximately a gamma distribution with shape parameter $(n-1)/2$ and scale parameter $2/(n-1)$ (*30*). Hence if the data from *n* cells were generated by a Poisson process, with probability 0.95 the computed Fano factor would lie in the interval $[\lambda_1, \lambda_2]$ where $\int_0^{\lambda_1} \Gamma(x; (n-1)/2, 2/(n-1))dx = 0.025$ and $\int_0^{\lambda_2} \Gamma(x; (n-1)/2, 2/(n-1))dx = 0.975$. In Table B-XIV we show the computation of the 95 % confidence interval for the Fano factor for all genes.

*Table B-XIV: Testing if the measured Fano factor could be due to finite sampling of Poisson noise*

For each gene we compute the confidence interval (CI) for its Fano factor given the number of cells and assuming mRNA fluctuations are Poissonian. Fano factor values in red indicate those which fall within the confidence intervals.

| Gene | Number of cells | Nuclear Fano factor | Cytoplasmic Fano factor | 95 % CI for Fano factor |
|---|---|---|---|---|
| **mad1** | 1382 | 0.929 | 0.620 | [0.927,1.08] |
| **mad1 − GFP** | 2155 | 0.983 | 0.657 | [0.941,1.06] |
| **mad2 − GFP** | 3064 | 0.845 | 0.824 | [0.951,1.05] |
| **mad3 − GFP** | 2661 | 0.974 | 0.725 | [0.947,1.05] |
| **bub1 − GFP** | 1142 | 0.951 | 0.621 | [0.920,1.08] |
| **sep1 − GFP** | 589 | 0.865 | 0.880 | [0.889,1.11] |
| **rpb1** | 1425 | 0.826 | 0.647 | [0.928,1.07] |
| **cdc13** | 1144 | 0.929 | 0.997 | [0.920,1.08] |

Note that *cdc13* is the only gene whose nuclear and cytoplasmic Fano factors both fall within the confidence intervals (also see Fig. A-I). On this basis, we cannot rule out that this gene has Poissonian expression characteristics, and hence we have not used data from this gene when fitting to the model. We note that for some genes their nuclear Fano factor falls within the confidence intervals whereas their cytoplasmic Fano factor does not (Table B-XIV). This might be expected due to the relationship between nuclear and cytoplasmic Fano factors when the nuclear export rate is high (Fig. 7H).

### Tables S1 – S4

#### Table S1: *S. pombe* strains

| Strain Number | Mating type | Genotype |
|---|---|---|
| JY001 | h- | |
| JY002 | h+ | |
| JY265 | h- | leu1 |
| JY743 | h- | leu1 ura4-D18 |
| SU161 | h- | leu1<<343nt-bub1+-ymeGFP-427nt |
| SU162 | h- | leu1<<82nt-bub1+-ymeGFP-427nt |
| SU163 | h- | leu1<<110nt-mad1+-ymeGFP-164nt |
| SU165 | h- | leu1<<13nt-mad1+-ymeGFP-164nt |
| SU167 | h+ | leu1<<49nt-mad2+-ymeGFP-521nt |
| SU169 | h- | leu1<<460nt-mad3+-ymeGFP-279nt |
| SU170 | h- | leu1<<286nt-mad3+-ymeGFP-279nt |
| SU171 | h+ | wis1+<<86nt-mad1+-ymEGFP-164nt |
| SU172 | h+ | wis1+<<48nt-mad1+-ymEGFP-164nt |
| SU173 | h+ | wis1+<<20nt-mad1+-ymEGFP-164nt |
| SU174 | h+ | wis1+<<13nt-mad1+-ymEGFP-164nt |
| SU184 | h- | wis1+<<286nt-mad3+-ymEGFP-279nt |
| SU185 | h+ | wis1+<<130nt-bub1+-ymEGFP-427nt |
| SU186 | h+ | mad3Δ::ymEGFP |
| SU186' | h+ | mad3Δ::ymEGFP |
| SU189 | h+ | wis1+<<170nt-bub1+-ymEGFP-427nt |
| SU190 | h- | rad21+-ymEGFP<<kanR |
| SU192 | h+ | wis1+<<392nt-mad3+-ymEGFP-279nt |
| SU193 | h+ | wis1+<<460nt-mad3+-ymEGFP-279nt |
| SU194 | h+ | wis1+<<332nt-mad3+-ymEGFP-279nt |
| SU194' | h+ | wis1+<<332nt-mad3+-ymEGFP-279nt |
| SU216 | h- | leu1 ura4-D18 mad2+-ymEGFP |
| SU228 | h+ | bub1+-ymEGFP |
| SU229 | h- | bub1+-ymEGFP |
| SU229' | h- | bub1+-ymEGFP |
| SU230 | h+ | bub1Δ::ymEGFP |
| SU231 | h- | bub1Δ::ymEGFP |
| SU499 | h- | leu1 ura4-D18 mad1+-ymEGFP |
| SU804 | h- | leu1 ura4-D18 mad3+-ymEGFP |
| SW130 | h+ | mad2+-ymEGFP |
| SW130' | h+ | mad2+-ymEGFP |
| SW132 | h+ | mad3+-ymEGFP |
| SW132' | h+ | mad3+-ymEGFP |
| SW139' | h- | mad2Δ::ymEGFP |
| SW140' | h+ | mad2Δ::ymEGFP |
| SW176 | h+ | mad1+-ymEGFP mad3+-ymEGFP |
| SW201 | h- | mad1Δ::ymEGFP |
| SW203 | h- | mad1Δ::ymEGFP |
| SW204 | h+ | mad1Δ::ymEGFP |
| SW205 | h+ | mad1+-ymEGFP |
| SW206 | h+ | mad1+-ymEGFP |
| SW222 | h+ | wis1+<<13nt-mad2+-ymEGFP-521nt |
| SW224 | h+ | wis1+<<49nt-mad2+-ymEGFP-521nt |
| SW225 | h+ | wis1+<<348nt-mad2+-ymEGFP-521nt |
| SW226 | h+ | wis1+<<58nt-mad2+-ymEGFP-521nt |
| SW226' | h+ | wis1+<<58nt-mad2+-ymEGFP-521nt |
| SW228 | h+ | wis1+<<780nt-mad1+-ymEGFP-164nt |
| SW229 | h+ | wis1+<<343nt-bub1+-ymEGFP-427nt |
| SW231 | h+ | wis1+<<82nt-bub1+-ymEGFP-427nt |
| SW232 | h+ | wis1+<<324nt-bub1+-ymEGFP-427nt |
| SW235 | h- | leu1+<<pDUAL-Pmad1-mad1+-ymEGFP |
| SW235' | h- | leu1+<<pDUAL-Pmad1-mad1+-ymEGFP |
| SW241 | h- | sep1+-ymEGFP<<hphNT1 |
| SW241' | h- | sep1+-ymEGFP<<hphNT1 |
| SW242 | h- | SPAC2H10.01+-ymEGFP<<hphNT1 |
| SW243 | h- | SPAC27D7.09c+-ymEGFP<<hphNT1 |
| SW244 | h- | leu1+<<pDUAL-Pbub1L-bub1+-ymEGFP |
| SW244' | h- | leu1+<<pDUAL-Pbub1L-bub1+-ymEGFP |
| SW245 | h- | leu1+<<pDUAL-Pbub1Lmut2-bub1+-ymEGFP |
| SW245' | h- | leu1+<<pDUAL-Pbub1Lmut2-bub1+-ymEGFP |
| SW249 | h+ | GFP-mad1+ |
| SW253 | h+ | leu1 his7+<<GFP-lacI-NLS but2+<<LEU2+<<lacO<<trz2+ |
| SW254 | h+ | leu1 his7+<<GFP-lacI-NLS hrd3+<<LEU2+<<lacO<<toa1+ |
| SW509 | h- | leu1<<13nt-mad2+-ymEGFP-521nt |
| SW510 | h- | leu1<<348nt-mad2+-ymEGFP-521nt |
| SW511 | h- | leu1<<231nt-mad2+-ymEGFP-521nt |
| SW513 | h+ | wis1+<<110nt-mad1+-ymEGFP-164nt |
| SW514 | h- | leu1<<58nt-mad2+-ymEGFP-521nt |
| SW642 | h+ | mad1+-ymEGFP |
| SW702 | h+ | wis1+<<58nt-Pmad2-rad21+-ymEGFP-521nt |
| SW703 | h+ | wis1+<<58nt-Pmad2-nmt1+-ymEGFP-521nt |
| SW704 | h- | leu1+<<pDUAL-Pmad1mut1-mad1+-ymEGFP |
| SW704' | h- | leu1+<<pDUAL-Pmad1mut1-mad1+-ymEGFP |
| SW705 | h- | leu1+<<pDUAL-Pmad1mut2-mad1+-ymEGFP |
| SW705' | h- | leu1+<<pDUAL-Pmad1mut2-mad1+-ymEGFP |
| SW712 | h+ | wis1+<332nt-Pmad3-nmt1+-ymEGFP-279nt |
| SW714 | h+ | wis1+<332nt-Pmad3-rad21+-ymEGFP-279nt |

**Table S2: sgRNA targeting sequences**

| Gene | Targeting sequence |
|---|---|
| mad2 | ATTGGGTAGACAGTGACCCT |
| mad2 | TATTCTTCATTAAGTTAGCA |
| mad2 | TGTTGTTCTATTATCACTTT |
| mad3 | GCAATTTACTCACCGTTGGT |
| wis1 intergenic | GTATGTGGCATACGCAGCCG |
| leu1 intergenic | GTAAGTACACAGCGACAACT |
| hph | TGCTCCATACAAGCCAACCA |

#### Table S3: FISH probes

| ymEGFP | mad1 | rpb1 | cdc13 |
|---|---|---|---|
| cagtgaataattcttcacctt | gaacggatccctaggagaatc | cgtaaggggacagaagaaggt | aactcgtcaagatgctggtt |
| tcaaccaaaattgggacaaca | aatctaggcaactgtgaacgc | cggggacaagattccaaattg | taggggaaaagtcctgttct |
| gaccattaacatcaccatcta | atttggtttcttaacgcttgt | cgcaacgctcattgaacgaat | tctagctctagacagtgact |
| ccttcaccggagacagaaaat | gcagaattaacagaaggcttc | ttcatccatggtctcaggaaa | ggcaagagcttttgagactt |
| gtcaatttaccgtaagtagca | agctagtttgggattttttgt | ggtttgacacttgaattgtcg | atagcctttgaccgaatcta |
| atggaactggcaatttaccag | ccttccgctttaaatcatttt | aaaatgaccaggacaatccgc | aacgtgaaggggtagtacgg |
| aaagtagtgactaaggttggc | actcaatttgttcacgctcaa | aactggctttgcaagttcaat | acgaattctgtcggggattg |
| tgttgtttcatatgatctggg | ttcttctgcaagttttctttg | ttttgcttaggaaaccgatgt | taggtgctggcgagttaaac |
| ctggcatggcagacttgaaaa | gctgtaacgaattcttctgtt | cgcaattccaacaaacgcatt | tttttgcacgggcaatatgg |
| gttctttcttgaacataacct | agttgcttttcaactagagtt | gatcacgataacgttgtgtgt | gaagacacagtggctttctt |
| agttaccgtcatctttgaaaa | gataagaagtagactgctcct | cagacattccaaactgcattt | caacgcatgacgcttcttag |
| ttgacttcagctctggtcttg | ccttttcatttcttacttctt | gtatcgcaaaccatttttgtc | agtgtgtctgacatttgtgt |
| taactaaggtatcaccttcaa | ttgcatcaagtagttcatgga | aaattatctgagcctgcagat | cacgacgggtactgacagaa |
| ataccttttaattcgattcta | ttcaactctgcaatctctttc | tattggcggagggattactaa | tcatctgttgcagggattat |
| cctaaaatgttaccatcttct | gatcattcttttctatctggg | tctttacgtattgtaggttgg | aaacagaaggttggcgacgc |
| gagagttatagttgtattcca | acttcatgattcttttcactt | caagaaccccataatctcagg | ttgaggtaacgaggggactg |
| gtcagccatgatgtaaacatt | tgcaaagcttgatttgagacc | aatctgattcatctttaccgc | gttgcttgatccttatggaa |
| actttgataccattcttttgt | gtttgtaagattggtatcctt | cgacagaaggcgtttttctgg | cgcgttcatcaacatctttc |
| gttgtgtctaattttgaagtt | ggaatccgcaaagagtttttc | atgtgtgaatattgtgtgcac | catcccaatcttgagattcg |
| attgaacagaaccatcttcaa | ttccttacacttcgtttcaag | actgctcattaagacctaagt | agggcatagtttcaatttcc |
| ttttgttgataatggtcagct | gaaagctcttgtaattgctgc | aataatcatccagtcaggtct | aagctctttttgacgatcca |
| agactggaccatcaccaattg | tcttccaattgctgattatga | cggaggaggaacaggtaagac | ccaatcggtaagtattccac |
| aagtaatggttgtctggtaac | aactagaaacctgcttgatgg | accgaaatactaggacggaca | aacaaggcagcaatgccaac |
| ggataaacgagattgagtgga | gcatttatttttccagttca | ttaaatcatcttcgccacgac | ccatccgccatatatacaaa |
| ctctcttttcgtttggatctt | cgctaatttgaagacgttgct | aacgttggcatttgcttttat | tcggcttgaagaatttcctc |
| aattctaacaagaccatgtgg | cgcagcttttaatttttccaa | ttcagagacaatatgcgcagg | tcataggattcgggtaagca |
| ggtaataccagcagcagtaac | gatagcttttcaattcgttct | tggcaacatggaactgaagta | agtcggcttttgagatacga |
| ttgtacaattcatccatacca | ccttgagaatttcaacattcc | cgctatttcattgtccatata | ctgttgggaagggggataag |
| | ttggactccaaatcgtttttc | tctgccagatttttgtaaagc | ctttttggaggcatacttct |
| | aaccttatccctatattcttc | cgtgcacggatacttttaaga | aatcttcatcagcgtcatcg |
| | tcattttcgagttcaagggta | gagctgaaaaatcaacccgtt | gaattatcggcgactcgttt |
| | gggttggtaactcgttagtaa | aatttggatcaccggtaatca | tatgcatctggcaatcttca |
| | acgagtttgtttgaaacagct | aacaccgagttcatctaacga | atcggcaagtagctcaacat |
| | ttagcattggtattctgtaga | agtgtcttagcaatactacgt | atattactccgactcatggt |
| | taaactagaaacgcgctctcc | aggtgtaacagtttcaggata | |

**Table S4: Primers for quantitative PCR**

| Target | Forward | Reverse |
|---|---|---|
| act1 | CCAAATCCAACCGTGAGAAGA | GTACGACCAGAGGCATACAAAG |
| cdc2 | GGTATCGTGCTCCTGAAGTATTG | CAGAGTCACCGGGAAATAATGG |
| mad1 | CCTAATGGGAGTGTTCGTGTTA | CCTGATGGATTACCAACCAATTTC |
| mad2 | TTAGAGCGGTGGCAGTTTAAT | CTCGCAGTTCATCTTCTTTGTTG |
| mad3 | CGGATGGTTCTGGAAAGGAT | CTACCCAAGAAGTAGCCGATATG |
| bub1 | TGTCACCAGCTATGCCTAAAG | TCGTGGCTACCGGATTACTA |
| ymEGFP | TGAAGGTGAAGGTGATGCTAC | CTAAGGTTGGCCATGGAACT |
