## Supplementary material for "The minimal intrinsic stochasticity of constitutively expressed eukaryotic genes is sub-Poissonian": Figures S1-S9

### Figure S1
Weidemann et al.

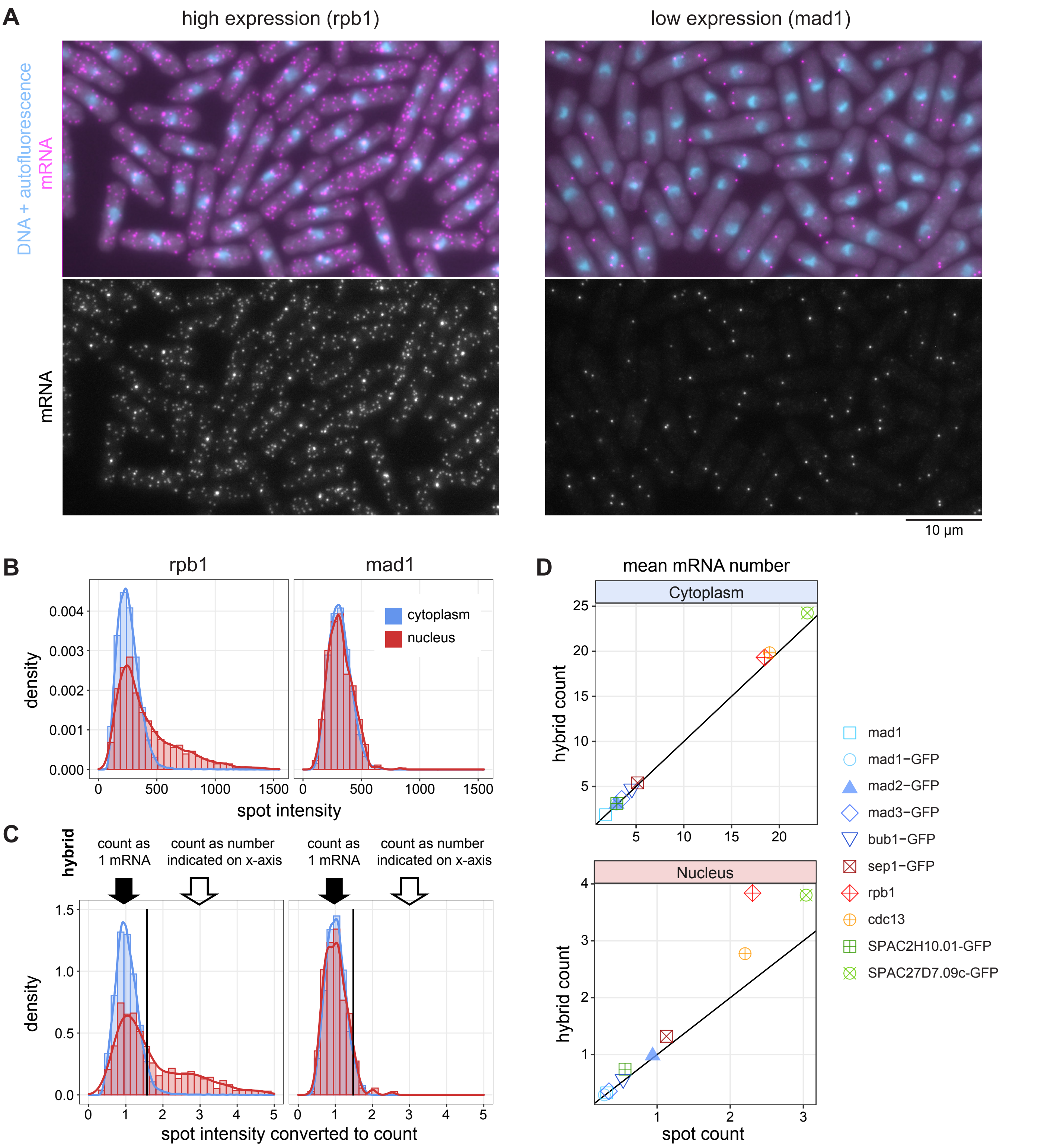


**Figure S1 - Comparison between "spot count" and "hybrid count" to determine mRNA number per cell**
**(A)** Representative images from smFISH experiments; DNA is stained with DAPI. The highly expressed gene (rpb1) shows higher intensity mRNA FISH spots in the nucleus, presumably representing transcription sites.
**(B)** Histogram and density distribution of FISH spot intensity ("AMP" from FISHquant) for rpb1 and mad1. Higher intensity spots are almost exclusively found for rpb1 and in nuclei. Pooled data from three independent experiments; n = 34,884 spots for rpb1; 3,424 spots for mad1.
**(C)** Single experiments from the data in (B) illustrating the "hybrid count" method. Data were normalized to the median of the spot intensity in the cytoplasm for each image. The vertical black line indicates the 95th percentile of intensities observed in the cytoplasm in this experiment and is used as cut-off between the count methods. n = 13,657 spots for rpb1; 1,511 spots for mad1.
**(D)** Comparison of mean mRNA counts per compartment obtained with the "spot count" method (which ignores spot intensities) and the "hybrid count" method illustrated in (C). Deviations are mostly observed for highly expressed genes and are most pronounced in the nucleus, as expected.

**Figure S2**
**Weidemann et al.**

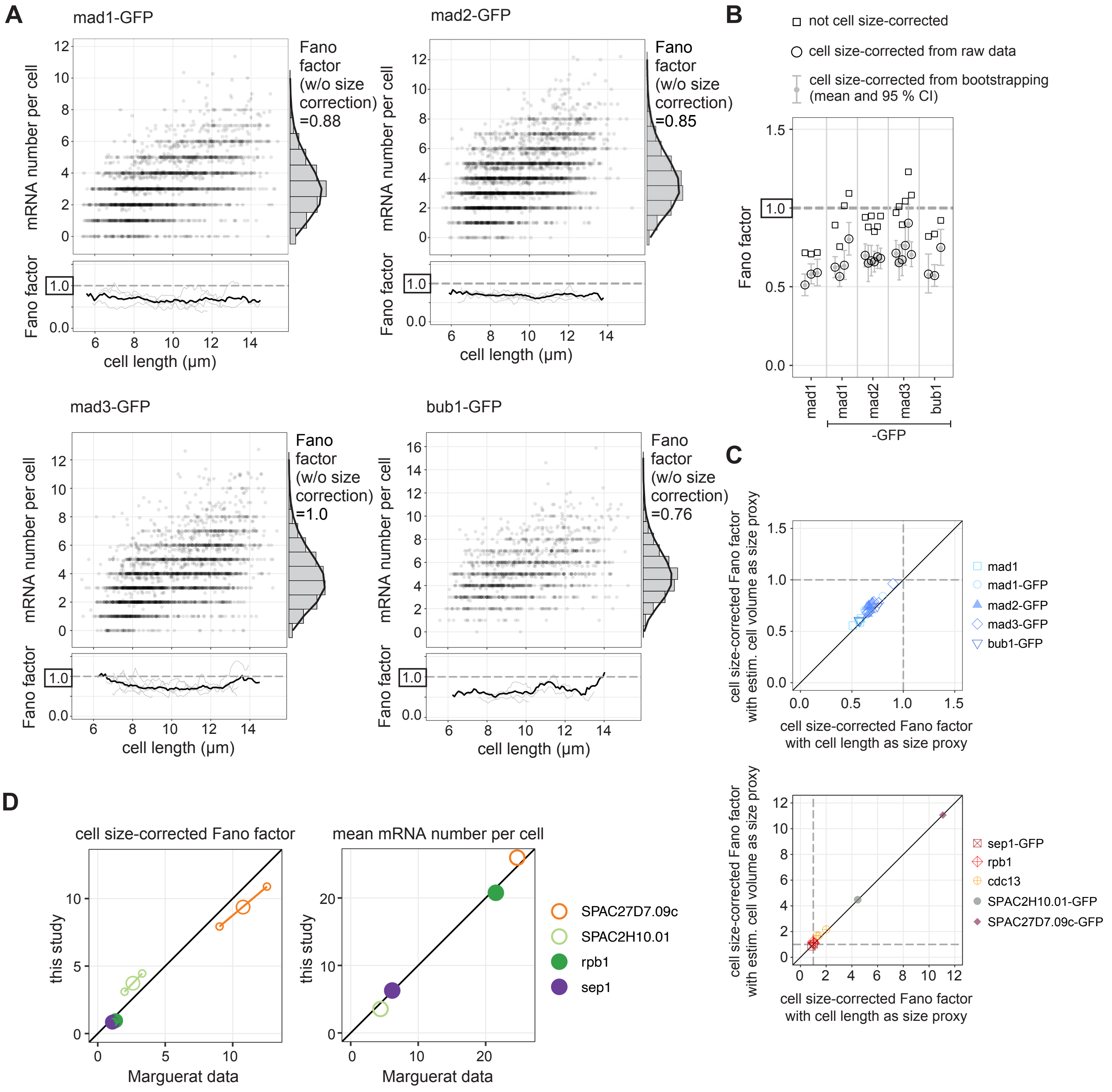


**Figure S2 - Additional data for sub-Poissonian mRNA distribution of SAC genes.**
**(A)** Scatter plots of cell length versus mRNA number per cell for mad1-GFP (n = 2,499 cells), mad2-GFP (n = 3,501 cells), mad3-GFP (n = 2,993 cells), and bub1-GFP (n = 1,358 cells), all expressed from their endogenous locus. Data from 3–6 replicates combined. Right: Histogram of mRNA number across all cells with fit to Poisson distribution. Bottom: The Fano factor was determined in a sliding window spanning 1 µm of cell length. The Fano factors for single replicates are shown in light gray; the Fano factor for the pooled data in black. Data for untagged mad1 is shown in Fig. 2.
**(B)** Comparison between not cell size-corrected and cell size-corrected Fano factors for SAC genes. The cell-size corrected Fano factors from bootstrapping (also shown in Fig. 2) and their 95 % confidence interval are shown in gray. 3 - 6 independent experiments for each gene are shown.
**(C)** Comparison of cell size-corrected Fano factors determined by either using cell length or cell volume as proxy for cell size. Cell volume is calculated from cell length, cell width, and the idealized assumption that an *S. pombe* cell is a cylinder with half-spheres at each end. Fano factors tend to be slightly higher when estimated cell volume is used, but they remain below 1.
**(D)** Direct comparison between values determined by the Marguerat group and in this study. Left: cell size-corrected Fano factors with 95 % confidence interval; right: mean mRNA numbers per cell. All data available for a single gene in each study were pooled. Note that SPAC27D7.09c, SPAC2H10.01, and sep1 were tagged with GFP in this but not in the Marguerat studies.

**Figure S3**
**Weidemann et al.**

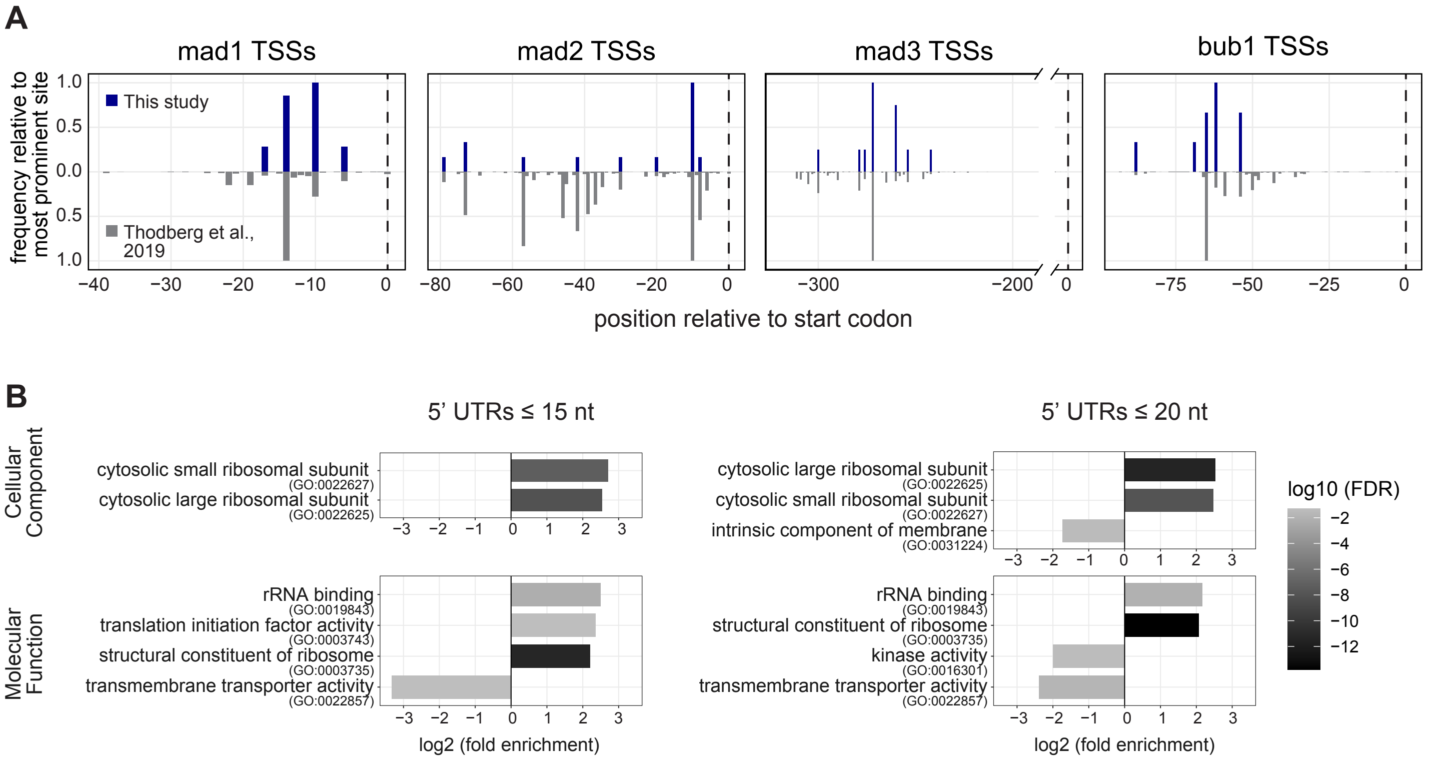


**Figure S3 - Additional data on SAC gene 5' UTRs.**
**(A)** Comparison of transcription start sites (TSSs) determined in this study by RACE-PCR, and determined in Thodberg et al., 2019, by cap analysis of gene expression (CAGE). Data are normalized by setting the most prominent TSS for each gene to 1. Data from this study is also shown in Fig 3.
**(B)** Gene Ontology (GO) enrichment analysis for genes with short 5' UTRs, as identified by Thodberg et al., 2019. Genes with 5' UTRs of 15 nucleotides or shorter (6.4th percentile), and 20 nucleotids or shorter (10th percentile) were analyzed. Ribosomal protein-coding genes are most prominently overrepresented. Value from Fisher's exact test with FDR correction is shown.

**Figure S4**
**Weidemann et al.**

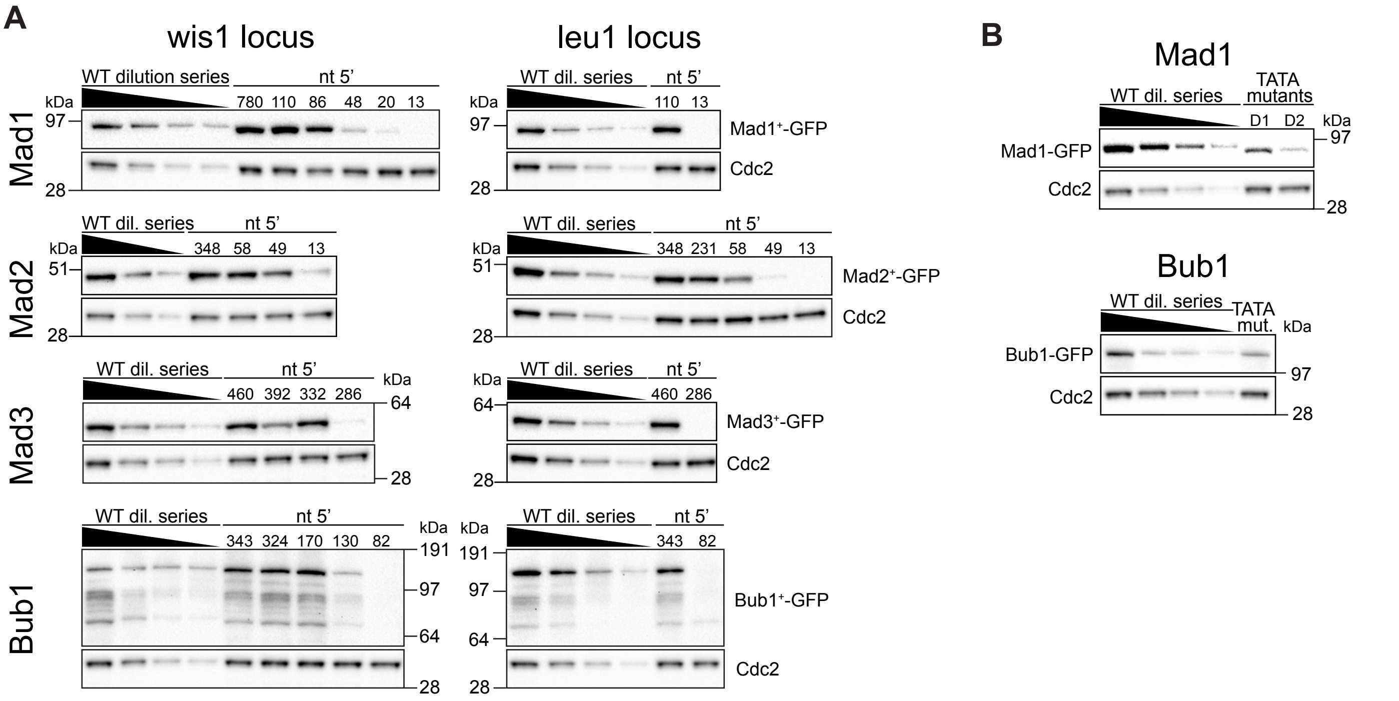


**Figure S4 - Additional data on SAC gene promoter mapping.**

**(A)** Immunoblots of protein extracts from cells with either the endogenous gene tagged with GFP (WT) or a genome fragment containing the GFP-tagged gene inserted at an exogenous locus (wis1 or leu1). The exogenous locus fragments for each gene contained a fixed sequence length 3' of the gene, but a variable number of nucleotides 5' of the start codon (nt 5'). A 1:1 serial dilution is loaded for WT extract. Quantification of these immunoblots is shown in Fig 4B.

**(B)** Examples of immunoblots quantified in Fig. 4E (one of two replicate experiments for each gene). Protein extracts from cells with either mad1-GFP or bub1-GFP inserted at the leu1 locus. The mad1 or bub1 promoter was either left intact (WT) or was mutated at the site of the proposed TATA box. A 1:1 serial dilution was loaded for the extract from cells with WT promoter.

### Figure S5
### Weidemann et al.

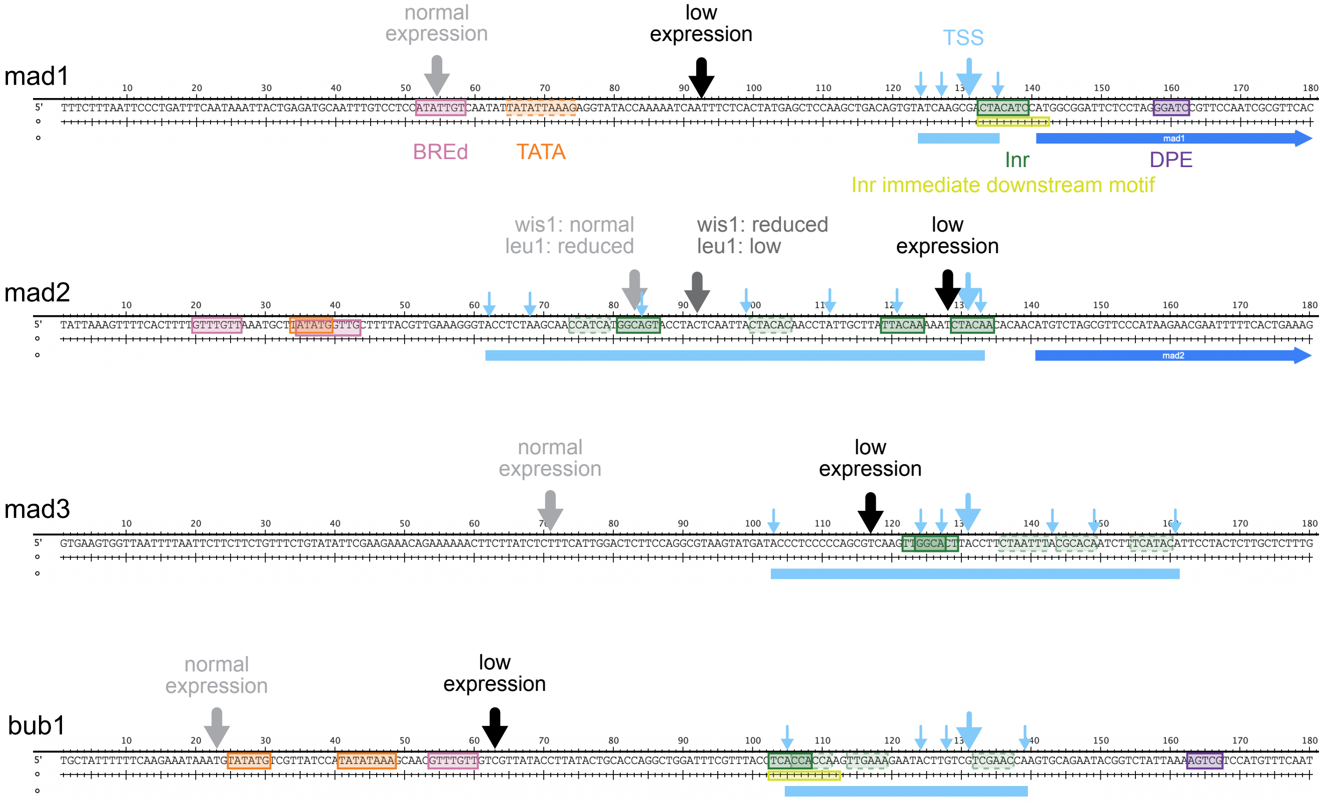


**Figure S5 - Potential core promoter motifs of *S. pombe* SAC genes.**
Sequence around the transcription start site (TSS) of SAC genes with potential core promoter motifs annotated. The range of **TSSs** observed is indicated by a **light blue** bar, individual TSSs by light blue arrows, the most frequent TSS by the larger arrow. Coding sequences for mad1 and mad2 are indicated in blue. Arrows in gray or black point to the 5’ ends of fragments that were or were not sufficient for expression (see Fig. 4). **TATA-box (orange)** identification used the consensus TATAWR (Vo Ngoc et al., Genes Dev 2017, PMID: 28808065). The region marked as dashed in mad1 has a mismatch to that consensus, but was included because it resembles the TATAWAWR consensus (Haberle and Stark, Nat Rev Mol Cell Biol 2018, PMID: 29946135), and because we found experimental evidence for its importance (Fig. 4). **BREd (pink)** identification used the RTDKKKK consensus (Vo Ngoc et al., Genes Dev 2017, PMID: 28808065; Haberle and Stark, Nat Rev Mol Cell Biol 2018, PMID: 29946135; Vo Ngoc et al., Genetics 2019, PMID: 31053615). Motifs within 10 nucleotides of a TATA box are shown, even when they were located upstream. BREu sequences (consensus SSRRGCC) were not found. **Inr sequences (green)** were annotated when located in the TSS region and matching one of the consensus sequences YYRNMM (identified in S. pombe, Li et al., RNA Biol. 2015, PMID: 25747261), YYANWYY (Roy and Singer, Trends Biochem Sci 2015, PMID: 25680757), or BBCABW (Haberle and Stark, Nat Rev Mol Cell Biol 2018, PMID: 29946135). Dark green boxes have an observed TSS in the correct position of the Inr, light green boxes do not. Sequences resembling an “Inr immediate downstream motif” (yellow), previously identified in S. pombe (Li et al., RNA Biol. 2015, PMID: 25747261), were found in mad1 and bub1, but not with the expected spacing relative to a TSS. **DPE (purple)** identification used the consensus RGWYV (Vo Ngoc et al., Genes Dev 2017, PMID: 28808065; Vo Ngoc et al., Genetics 2019, PMID: 31053615).

**Figure S6**
**Weidemann et al.**

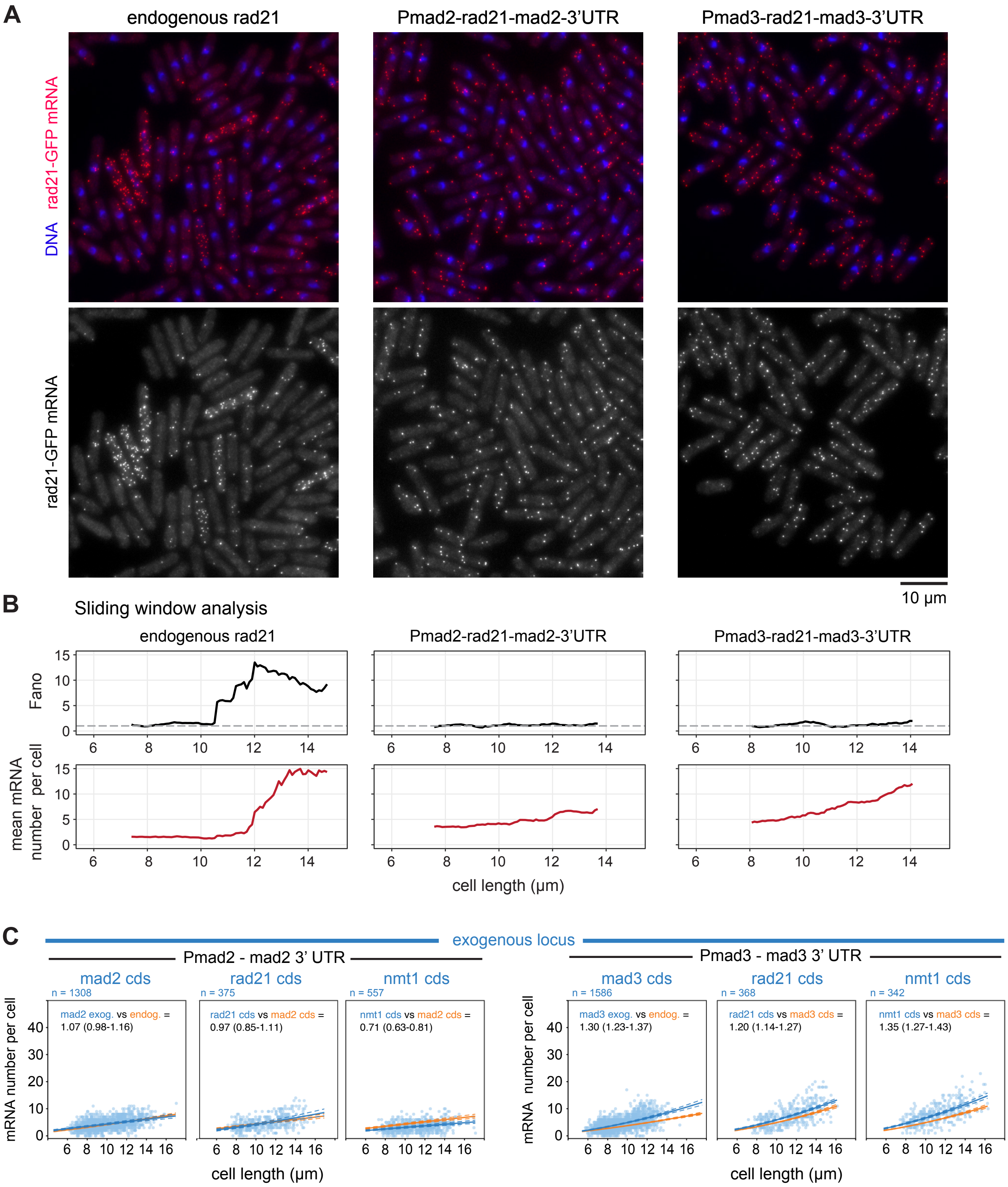


**Figure S6 - Expression of rad21 from mad2 or mad3 regulatory sequences reduces expression level and Fano factor.**
**(A)** Example images of rad21-GFP mRNA staining by smFISH. DNA is shown in blue (stained by DAPI). Expression from the endogenous rad21 locus (left), or from the wis1 locus with mad2 regulatory sequences (middle) or mad3 regulatory sequences (right).
**(B)** Same experiment as in (A). The Fano factor (black) and the mean mRNA number per cell (red) was determined in a sliding window spanning 1 µm of cell length. Only cell lengths, for which more than 35 cells were available for quantification are shown.
**(C)** Same experiment as in Fig. 5, except that "spot count", not "hybrid count" is shown. mRNA number relative to cell length at the exogenous locus for expression from the *mad2* promoter (*Pmad2*) and *mad2* downstream region or *mad3* promoter (*Pmad3*) and *mad3* downstream region. Solid blue lines are regression curves from generalized linear mixed model fits for the data shown, solid orange lines are regression curves for the reference data; dashed lines indicate the 95% bootstrap confidence bands for the regression curves. Left panel on each side: comparing mad2 or mad3 expression at the exogenous locus to the endogenous locus; middle and right panel on each side: comparing rad21 or nmt1 cds at the exogenous locus to mad2 or mad3 cds at the exogenous locus. Model estimates of the ratio with bootstrap 95% confidence interval in brackets are shown on the top.

### Figure S7
### Weidemann et al.

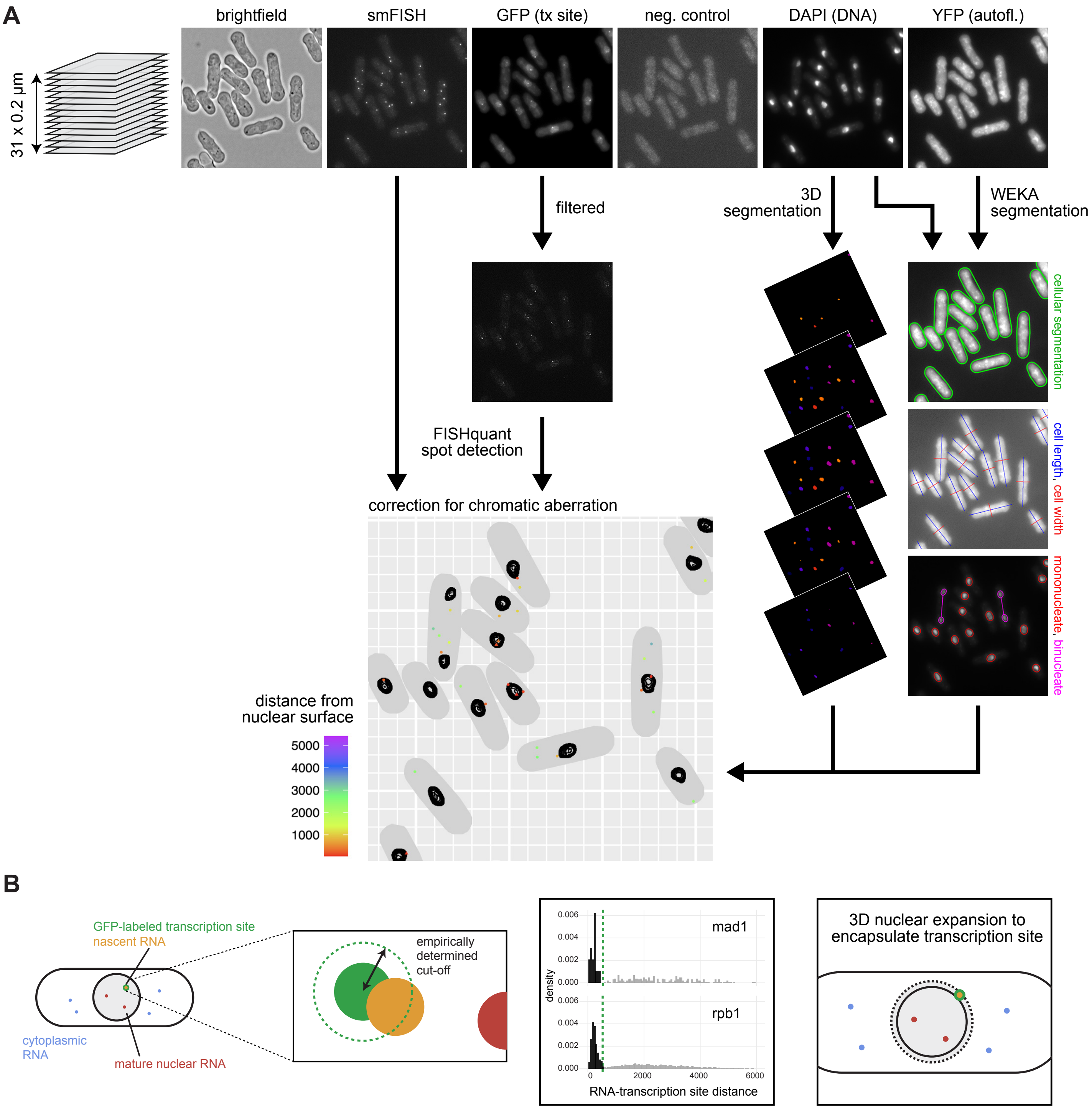


**Figure S7 - Segmentation and image processing steps to distinguish nascent, nuclear and cytoplasmic mRNA.**
**(A)** Image analysis pipeline. 31 Z-sections, spaced by 0.2 µm, were recorded in six channels. Sub-resolution fluorescent beads were imaged using the same settings (not shown), and this information was used to correct for chromatic aberration. Autofluorescence recorded with filters for yellow fluorescent protein (YFP) was used to segment cells in 2D by trainable WEKA segmentation. In addition, nuclei were segmented in 3D based on staining DNA with DAPI. Single-molecule mRNA FISH spots and GFP-labeled transcription sites were identified using FISHquant. After correcting for chromatic aberration, FISH spots were classified as nuclear if located within the 3D-segmented nucleus, and as cytoplasmic otherwise.
**(B)** FISH spots showed a bimodal distribution for distance to transcription sites, and this information was used as cut-off to distinguish nascent mRNA from mature nuclear or cytoplasmic mRNA. In around 16 % of the cells, the transcription site was located just outside of the segmented nucleus. In these cases, the nuclear border was expanded to incorporate the transcription site. In an alternative approach, we excluded these cells from the analysis, which did not change the conclusions (see Fig. 7E vs. S9B).

### Figure S8
### Weidemann et al.

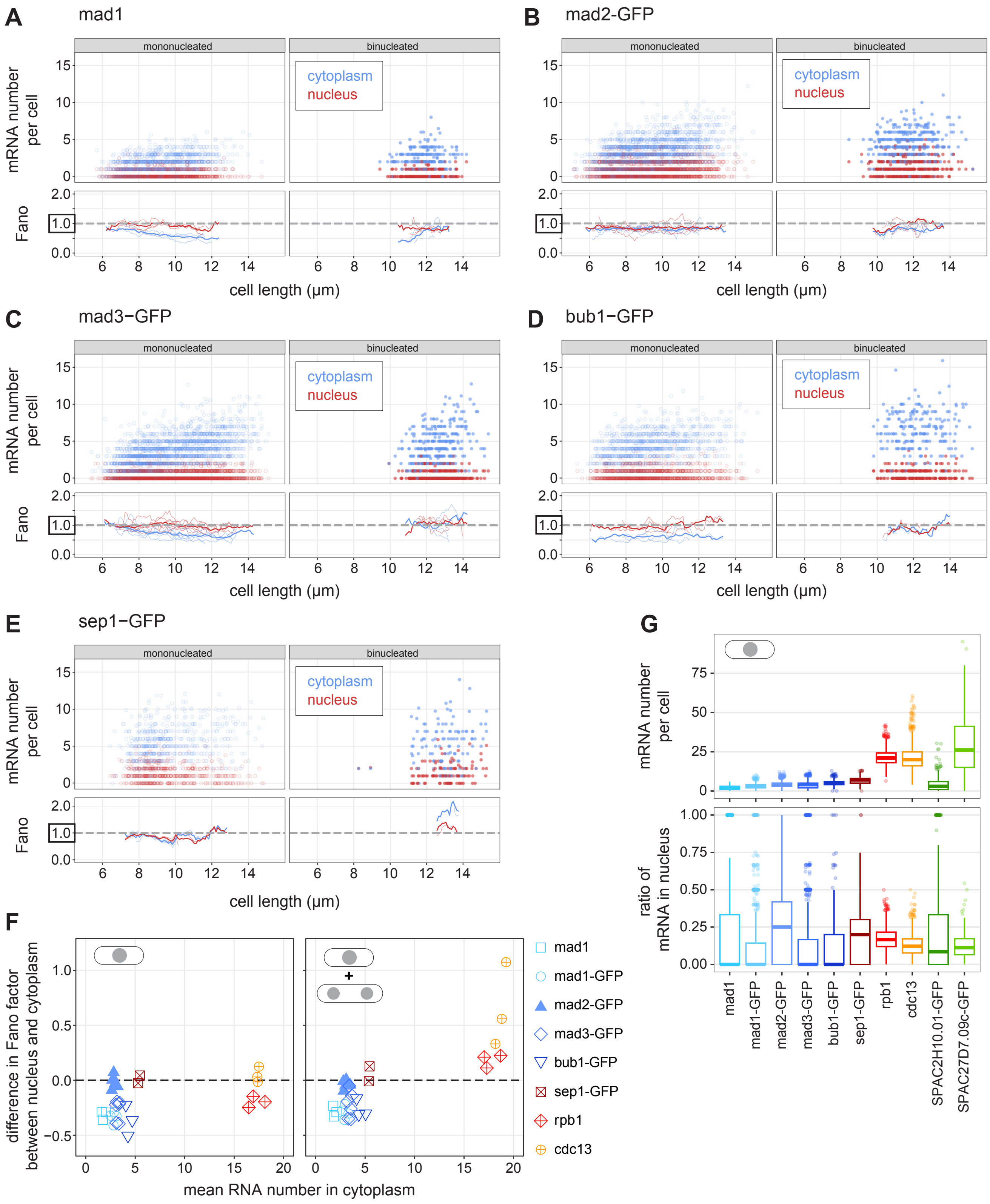


**Figure S8 - Additional data on Fano factors of the nuclear and cytoplasmic mRNA distributions.**
**(A-E)** Top: Scatter plot of cell length versus mRNA number in cytoplasm (blue) or nucleus (red). Mono- and binucleated cells are shown separately. Bottom: The Fano factor was determined in a sliding window spanning 1 μm of cell length. The Fano factors for single replicates are shown as thin lines; the Fano factor for the pooled data as thick line. See Fig. 6 for other genes. **(F)** For each experiment, the mean RNA number in the cytoplasm was calculated for cells with lengths in the interquartile range of all experiments (8.1–11.3 μm for all cells, 8.0–10.6 μm for mononucleated cells); mean RNA number is shown in relation to the difference between the Fano factor in the cytoplasm and the nucleus (data from Fig. 6A). **(G)** Number of mRNA molecules in mononucleated cells (top) and fraction of these mRNAs in the nucleus (bottom); number of cells per gene between 212 and 3,064. Boxplots show median and interquartile range; whiskers extend to values no further than 1.5 times the interquartile range from the first and third quartile, respectively.

### Figure S9
### Weidemann et al.

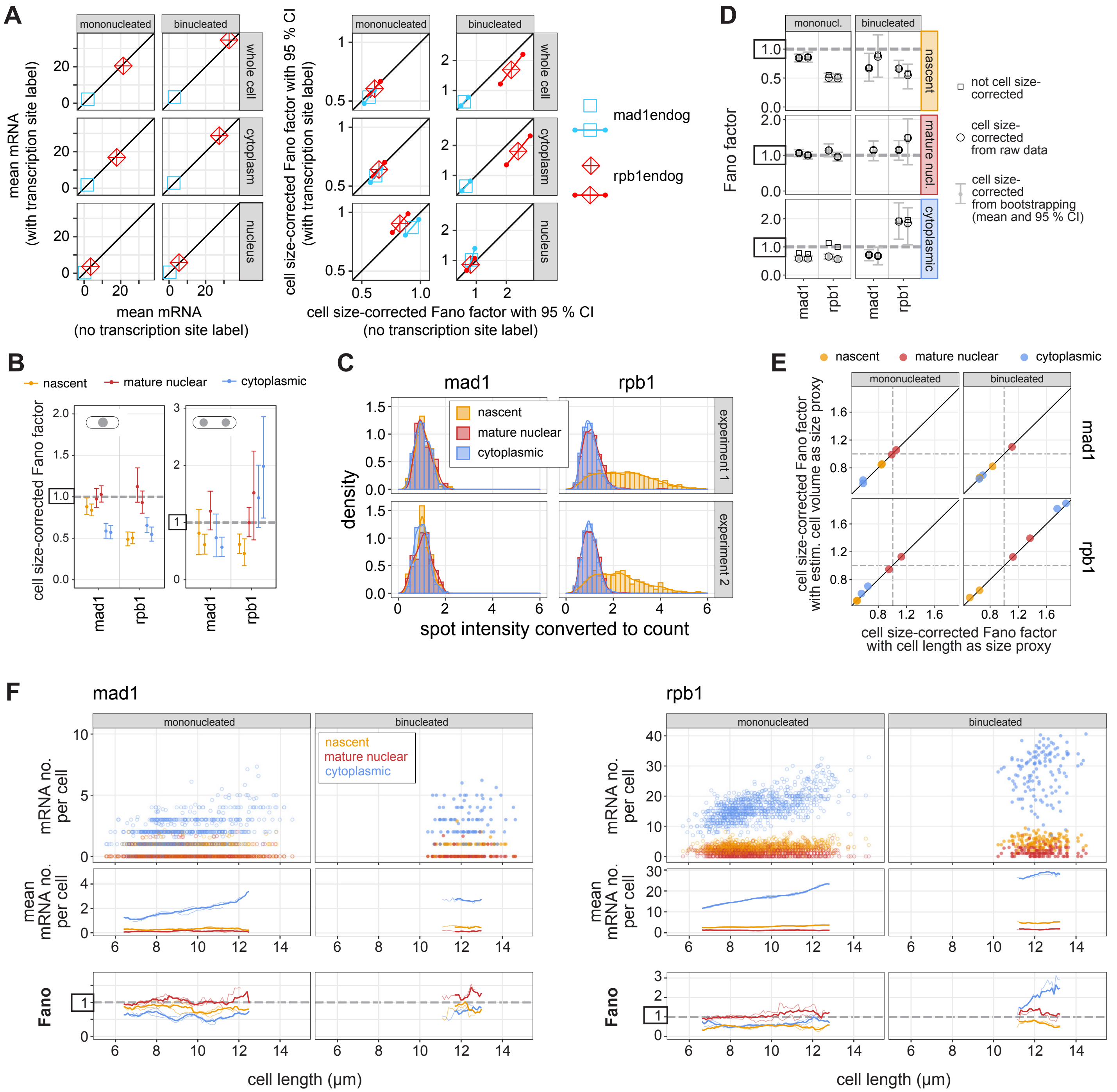


**Figure S9 - Additional data from cells with transcription site labeled.**
**(A)** Comparison of mean mRNA number and cell size-corrected Fano factor for mad1 and rpb1 measured in cells without transcription site label (x-axis) or with transcription site label (y-axis). **(B)** Same analysis as in Fig. 7D, except that cells in which the GFP-labelled transcription start site was not within the segmented nucleus were excluded. This removed between 11 % (mad1, mononucleated) and 36 % (rpb1, binucleated) of cells, yet yielded highly similar results. **(C)** Histogram and density distribution of FISH spot intensity at the transcription site (nascent), at other positions in the nucleus (mature nuclear), or in the cytoplasm. Data were normalized to the median of the spot intensity in the cytoplasm for each image (n = 1,411 and 806 spots for mad1; n = 11,244 and 9,496 spots for rpb1). Spots identified as nascent by their proximity to the GFP-labeled rpb1 gene contain a higher number of mRNAs, which (i) reflects strong transcription of rpb1, and (ii) suggests that the identification of transcription sites by proximity to GFP is accurate. **(D)** Comparison between not cell size-corrected and cell size-corrected Fano factors for mad1 and rpb1 in different compartments. The cell size-corrected Fano factors from bootstrapping and their 95 % confidence interval (also shown in Fig. 7) are shown in gray; 2 independent experiments for each gene. **(E)** Comparison of cell size-corrected Fano factors determined by either using cell length or cell volume as proxy for cell size. Cell volume is calculated from cell length, cell width, and the idealized assumption that an *S. pombe* cell is a cylinder with hsalf-spheres at each end. Individual experiments are shown as dots. **(F)** Top: Scatter plot of cell length versus mRNA number at the transcription site (nascent, orange), in the nucleus, but not associated with the transcription site (mature nuclear, red), or in the cytoplasm (blue). Mono- and binucleated cells are shown separately. Middle: The mean mRNA number in the three different compartments was determined in a sliding window spanning 1 µm of cell length. Bottom: The Fano factor in the three different compartments was determined in a sliding window spanning 1 µm of cell length. For the sliding window data, single replicates are shown as thin lines; pooled data as thick line.
